# Molecular Analysis and Computational Modeling Reveal Temporally Separable Responses triggered by DENV-Induced Soluble Factors in Endothelial Cells

**DOI:** 10.64898/2026.03.20.713013

**Authors:** Jenny Paola Alfaro-García, Julieta M. Ramírez-Mejía, Paola Rojas-Estevez, Diego A. Álvarez-Díaz, Geysson Javier Fernández, Carlos Alberto Orozco-Castaño, Boris Anghelo Rodríguez-Rey, Juan Carlos Gallego-Gómez, Miguel Vicente-Manzanares

## Abstract

Dengue virus (DENV) represents a growing global health challenge with billions of people at risk. Severe Dengue (SD), a complication of DENV infection that involves generalized hemorrhage, is driven, at least in part, by endothelial dysfunction. Endothelial dysfunction refers to increased permeability due to inflammation, mechanical injury and/or modification of the genetic program of endothelial cells. Previous work showed that exposure of endothelial cells to conditioned media from DENV-infected cells (CMDV) increased permeability and cellular stiffness, repressed endothelial markers and induced mesenchymal genes. However, the generality, extent, mechanism and ultimate impact of these events in the onset of SD remain elusive. Here, we integrate analysis from in vitro infection of endothelial cells with computational modeling to investigate the key features of CMDV-induced endothelial alterations and their potential impact on endothelial dysfunction. We found that CMDV increased *SNA1* and *CDH2* expression, while suppressing endothelial genes *OCLN* and *CDH5*. Global transcriptomics analysis revealed that CMDV triggered a transient pro-inflammatory response, followed by induction of selected tissue repair genes and matrix remodeling. A non-directed asynchronous network model (NDAM-CMDV) identified IL6 and FN1 as central nodes of DENV-induced endothelial trans-differentiation, providing new molecular insights that predict the evolution of the disease and identify potential therapeutic targets.

## 1. Introduction

Dengue virus (DENV) is currently the most prevalent arbovirus in the world. The largest outbreak of dengue was recorded between 2023 and 2024, underscoring its growing prominence as a global threat to human health. Estimates suggest that half of the human population may be at risk [1–3]. Of particular concern is the progression of dengue to severe dengue (SD), a rare but life-threatening condition characterized by generalized hemorrhage. There is no specific treatment for SD, only palliative care [4, 5].

Hemorrhage in SD is due to several, poorly characterized, factors. A central feature of SD is endothelial dysfunction, which seems caused by activation of the innate immune response; expression of pro-inflammatory mediators that activate endothelium; antibody-dependent enhancement of viral infection [6, 7]; and/or extracellular matrix (ECM) remodeling in response to cytopathic damage [8]. Notably, SD typically arises during the defervescence stage [4], when viremia is declining [9]. Conversely, viral proteins, e.g., NS1, and DENV-induced host factors such as IL-6, TNF-α, NF−κB, TGF-β and others persist during this stage [10–12].

The expression and effect of those soluble factors can be recapitulated in vitro using conditioned media from DENV-infected cells (CMDV). In this model, media are irradiated with UV light, which inactivates the virions but does not affect viral proteins and host factors. Previous reports have indicated that incubation of epithelial and endothelial cells with CMDVs activate migratory pathways involving c-Abl and RhoA [13–15]. Additionally, CMDVs induce a transient reorganization of the cytoskeleton. They also decrease the expression of selected endothelial markers such as ZO-1 and VE-Cadherin [14, 16], while increasing expression of mesenchymal markers such as SNAIL, TWIST1, α-SMA and N-Cadherin [16]. These effects were concurrent with increased endothelial permeability, which is partially reversed by treatment with imatinib [14].

These data strongly suggest that the host soluble factors induced by DENV infection trigger increased permeability due to the weakening of cell-cell junctions and the acquisition of migratory features. These changes are also consistent with a process known as Endothelial-to-Mesenchymal Transition (EndMT)[14, 16]. EndMT is defined as a reversible trans-differentiation process that occur in healthy and pathological processes, such as wound healing and cancer, respectively [17]. Interestingly pathogens such as bacteria [18–20] and viruses [21–23] may also trigger EndMT. It is important to note that EndMT is not a binary switch in which cells remain completely endothelial or turn completely mesenchymal. Instead, different studies highlight the existence of multiple intermediate stages [24–26]. In the present study, we pose that CMDVs may induce changes in endothelial cells that recapitulate specific aspects of EndMT. Such events would constitute a suitable explanation for the conspicuous absence of significant post-mortem evidence of inflammatory damage in patients who succumbed to SD [27–29]. This confirms previous studies from the group that had provided partial evidence for such possibility [14, 16]. However, the kinetics of the process and the potential overlap and/or synergy with the inflammatory response remain unknown. In addition to recapitulating and quantifying the effect of CMDV in selected markers of endothelial and mesenchymal lineages, global transcriptomics have revealed a genetic link to the biphasic response observed in severe dengue patients, that is, high viremia associated with elevated fever (where we observe an induction of pro-inflammatory mediators), followed by more severe symptoms during the defervescing phase, where we detect induction of endothelial plasticity. Together, these data enabled us to generate an unbiased model based on transcriptomic data from endothelial cells treated with CMDV compared to untreated cells and cells treated with TGF-β, a bona fide inducer of EndMT. Computational modeling and perturbation analysis predict a biphasic response to DENV infection in endothelial cells, characterized by an initial period dominated by the inflammatory response (when viremia is reportedly high), followed by a repair/remodeling response later on, during the defervescence stage, when the viremia is reportedly declining.

In summary, our findings provide in vitro and in silico support to the hypothesis that partial EndMT induced by soluble factors stemming from DENV infection plays a crucial role in the pathogenesis of SD, forming the conceptual basis of novel approaches that could identify critical targets to treat SD patients.

## 2. Results

### 2.1. CMDV induces expression of mesenchymal markers and morphological alterations in endothelial cells

To evaluate the effect of CMDV on endothelial trans-differentiation, HMEC-1 (endothelial) cells were exposed for 48 hours or 120 hours to CMDV or 5 ng/mL TGF-β1. The latter was used as a positive control that induces EndMT [30]. In agreement with previous results, CMDV had no cytopathic effect [16]. At a protein level, CMDV neither induced expression of N-cadherin (*CDH2* gene) nor reduced expression of VE-cadherin (*CDH5* gene) at 48 hours **(Figure 1A-B)**. However, N-cadherin induction and VE-cadherin reduction by CMDV were significant after 120h **(Figure 2A-B)**. Similarly, CMDV did not affect Snail (*SNA1* gene) or occludin (*OCLN* gene) expression after 48h **(Figure 1C)**, but reduced occludin and increased SNAIL expression significantly after 120h **(Figure 2C)**. Conversely, N-cadherin expression was induced more rapidly (48h) by TGF-β1 **(Figure 1A-B)**. Interestingly, TGF-β1 slightly increased VE-cadherin expression **(Figure 1B)** and had no significant effect on the expression of occludin and SNAIL **(Figure 1C)** at 48h. However, TGF-β1 reduced VE-cadherin and occludin expression and increased SNAIL levels after 120h, as expected **(Figure 2B)**, suggesting that the loss of endothelial markers and the acquisition of mesenchymal markers during TGF-β1-induced EndMT is not a linear process.

**Figure 1.**
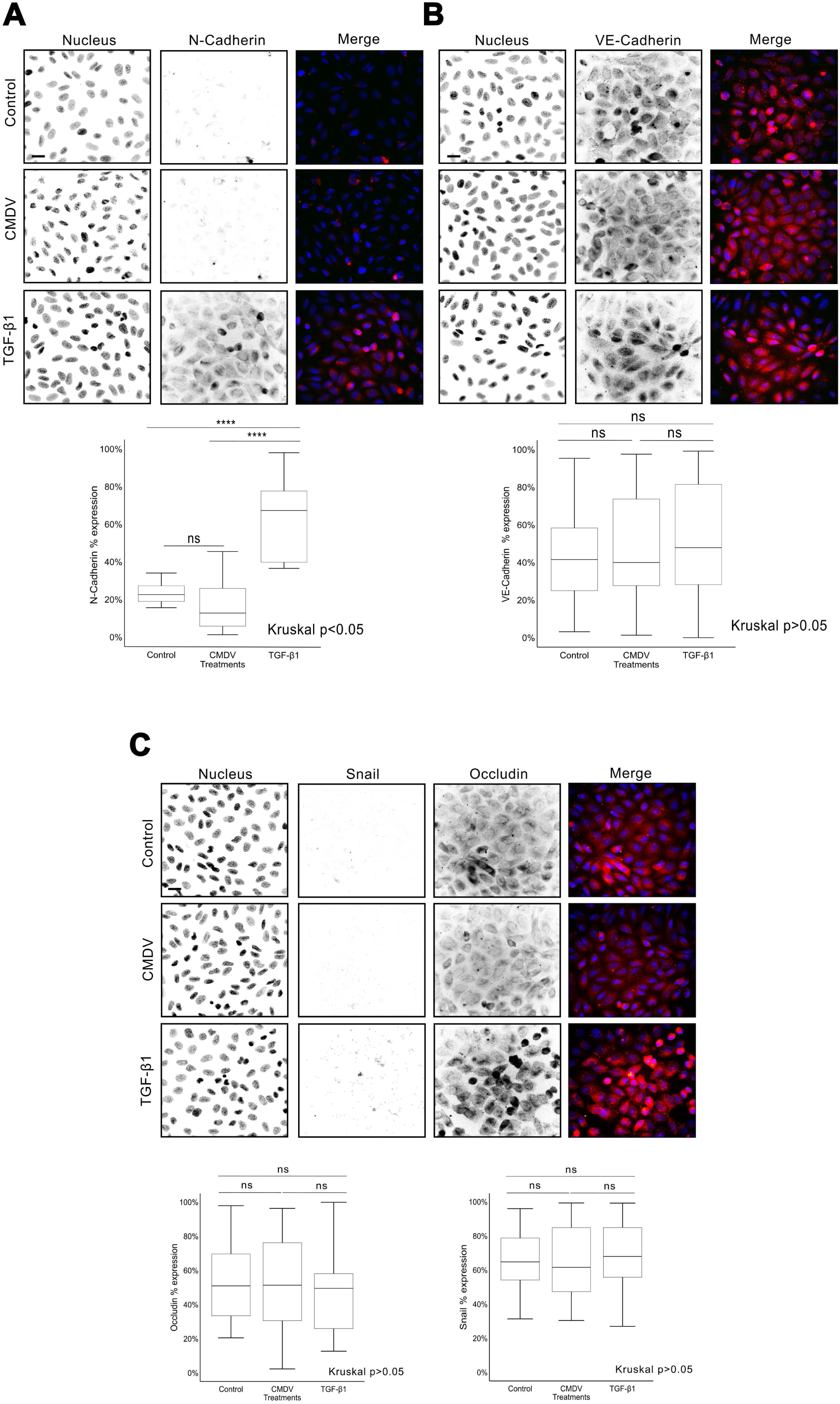
Effect of CMDV and TGF-β1 on the expression of selected endothelial and mesenchymal markers in HMEC-1 endothelial cells at 48h. Representative images of HMEC-1 cells treated as indicated and stained for DNA (DAPI) and N-Cadherin (A), VE-Cadherin (B) and occludin and Snail (C). In overlays, red is N-cadherin (A), VE-cadherin (B) or occludin (C) as indicated, and green is Snail (C only); blue represents DAPI staining in all overlays. Scale bar=20 µm. Bottom, quantitative analysis of images as in top. See Material and Methods for details. Data are presented as mean ± SEM from >100 fields examined in three independent experiments. Statistical tests are indicated, and significance is as follows: *p < 0.05, **p < 0.01, ***p < 0.001, ****p < 0.0001.

**Figure 2.**
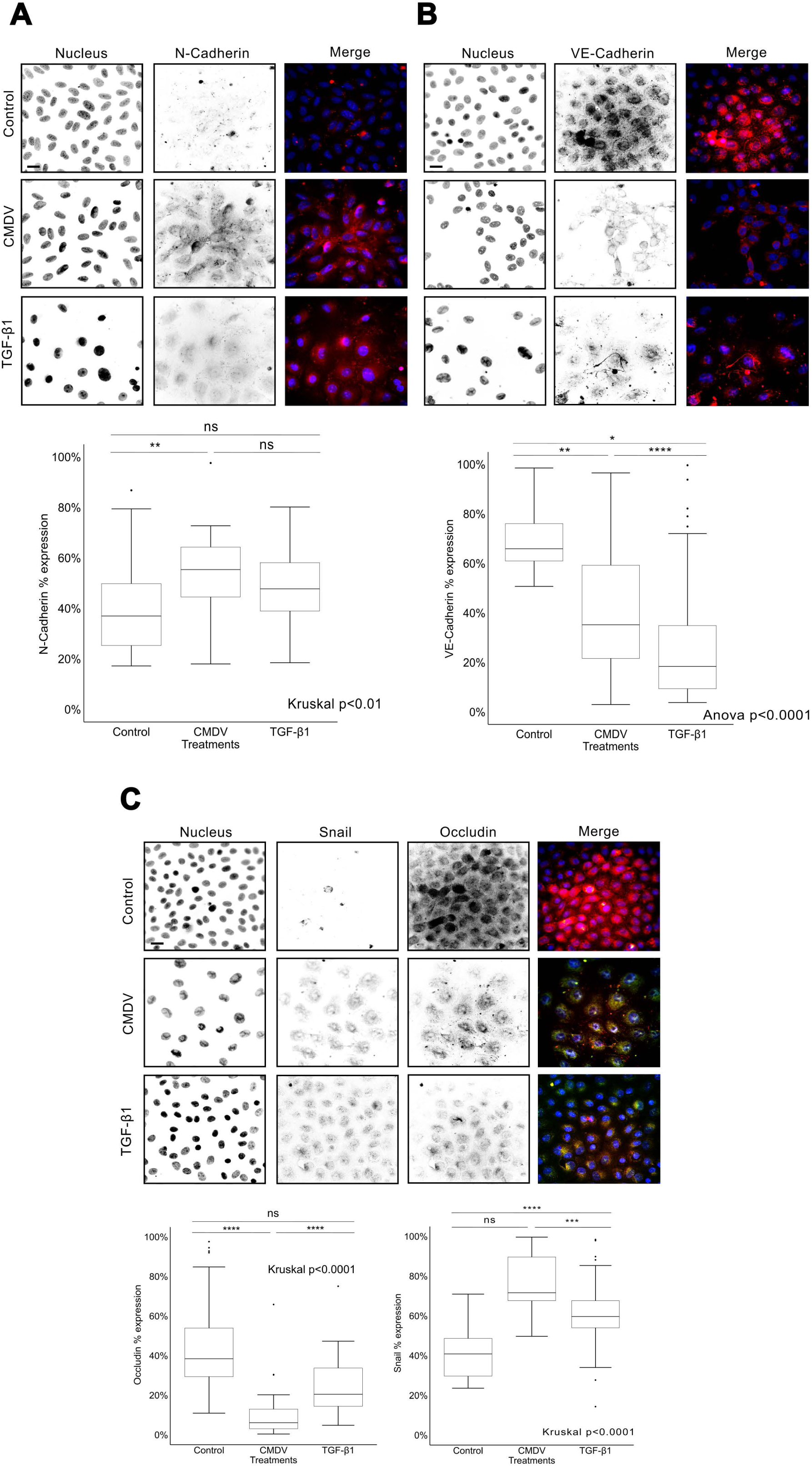
Effect of CMDV and TGF-β1 on expression of selected endothelial and mesenchymal markers in endothelial cells at 120h. Representative images of HMEC-1 cells treated as indicated and stained for DNA (DAPI) and N-Cadherin (A), VE-Cadherin (B) and occludin and Snail (C). In overlays, red is N-cadherin (A), VE-cadherin (B) or occludin (C) as indicated, and green is Snail (C only); blue represents DAPI staining in all overlays. Scale bar=20 µm. Under the microphotographs, quantitative analysis of images as in top. See Material and Methods for details. Data are presented as mean ± SEM from >100 fields examined in three independent experiments. Statistical tests are indicated, and significance is as follows: *p < 0.05, **p < 0.01, ***p < 0.001, ****p < 0.0001.

Imatinib, a c-Abl inhibitor previously shown to counter some of the effects of exposure to CMDV [14, 16], curbed the increased expression of Snail and N-cadherin and prevented the decrease of VE-cadherin and occludin **(Supplementary Figures 1 and 2**). However, imatinib-treated cells did not recover their morphological integrity, likely due to direct effects of the inhibitor on the actin cytoskeleton [31].

### 2.2. CMDV transiently enhances pro-inflammatory gene expression

We next performed RNAseq of CMDV– and TGF-β1-treated endothelial cells compared to untreated cells, in conditions similar to those in **Figure 1-2**. 48 hours post-treatment, we found 98 DEGs (differentially expressed genes), 75 upregulated and 23 downregulated in response to CMDV. After 120 hours of incubation with CMDV, the total DEG number increased to 742, 401 upregulated and 341 downregulated. Conversely, TGF-β1 treatment for 48 hours produced 840 DEGs, 401 upregulated and 439 downregulated. After 120 hours, we found 1506 DEGs, 827 upregulated and 679 downregulated.

Analysis of DEGs (differentially expressed genes) expressed by HMEC-1 cells in response to CMDV revealed that the treatment triggered two distinct gene expression patterns: an initial pro-inflammatory response at 48 hours, including less than 100 genes, for example IL-6, CXCL1, CCL1/6 and several growth factors. At 120h, we observed an endothelial tissue repair program containing over 800 DEGs, including repair genes such as MYC, TENM2 and specific forms of collagen, e.g. COL5A2 **(Figure 3A-B and Supplementary Figure 3A-B)**. This response is markedly different from that induced by TGF-β1 at 48h **(Figure 3C-D and Supplementary Figure 3C-D)**, which induced over 800 DEGs including EndMT-related genes and cell junctions. At 120h, >1500 DEGs that included EndMT and cell cycle-related proteins. This indicates that the effects of CMDV are not solely due to TGF-β1 induced during DENV infection, as seen in peritoneal macrophages [32].

**Figure 3.**
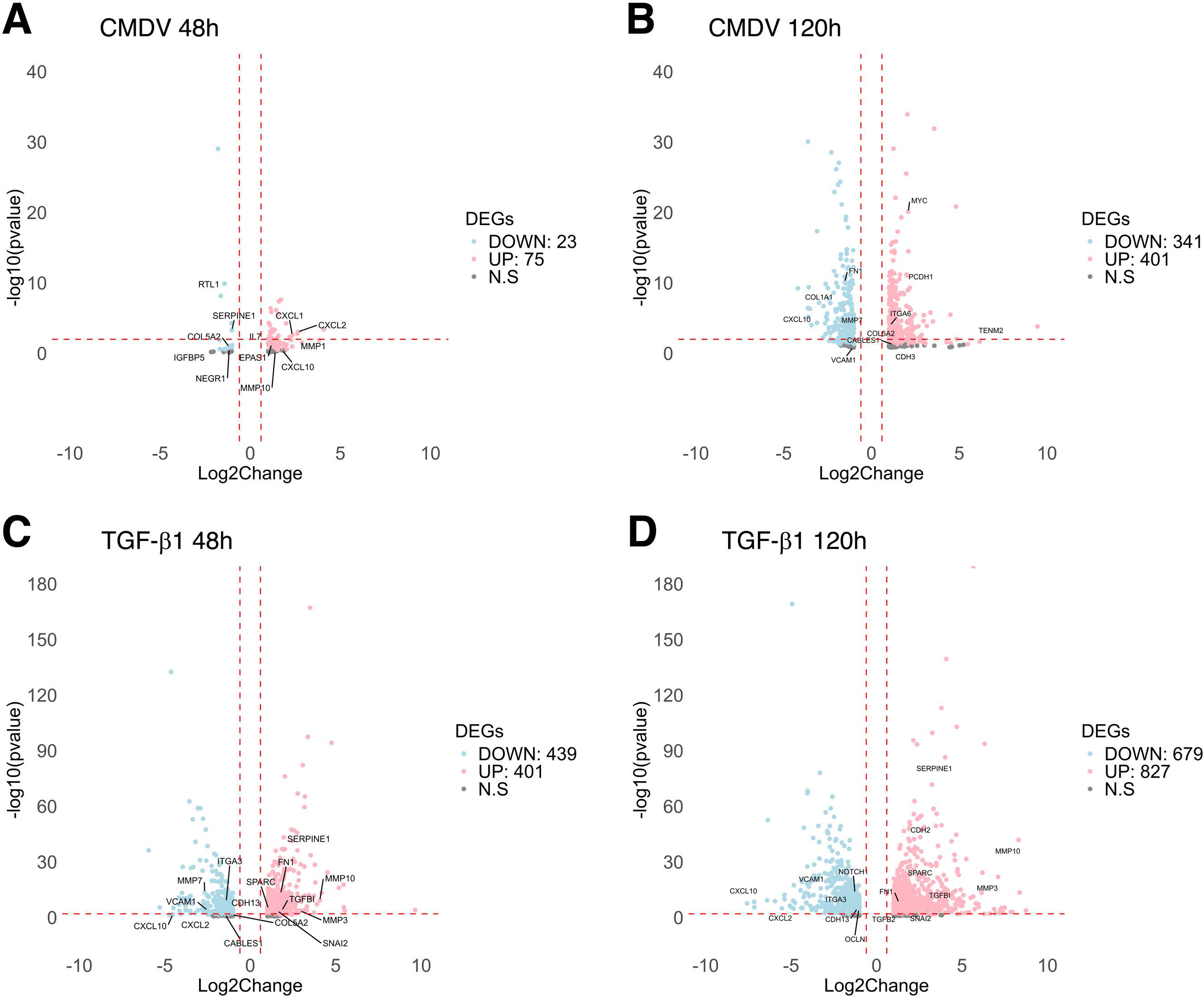
Volcano plots depicting DEGs at 48h and 120h after exposure to CMDV (A-B) and TGF-β1. (**C-D**). See Material and Methods for technical details on sample collection and analysis. In the graphs, red means upregulated and blue means downregulated. Grey represents no significance (genes are located beyond the red dotted lines, denoting the significance threshold). Experiment is representative of three independent replicates.

As expected, TGF-β1 did not trigger the expression of inflammatory genes at 48h, whereas CMDV mainly induced inflammatory cytokines such as CXCL1, CXCL2, CCL2 and IL7; other inflammatory mediators such as IRAK3 and NRP2; and metalloproteases such as MMP1 and MMP10. Pharmacological interaction analysis (DGIdb) identified potassium alum as a potential modulator of IL-7 in this phase **(Supplementary Table 1)**. TGF-β1 triggered mesenchymal genes such as *SNAI2* (Slug protein), *SMAD7*, *TGF-β1*, *ITGA2* and others, while inflammatory markers such as *CXCL6*, *CXCL8* or *STAT4* were repressed. Interestingly, additional pharmacological potential targets emerged in this phase, including Anakinra, which is an antagonist of IL1R2 (Supplementary Table 1). Conversely, 120h of CMDV treatment induced a response focused on cell repair, including genes such as MYC. At 120h, most of the inflammatory genes induced at 48h were repressed **(Figure 3B)**.

On the other hand, 120h exposure to TGF-β1 promoted expression of mesenchymal markers such as *TGF-β1/2/3*, *SNAI2*, *FN1* and *VCL*, among others **(Figure 3D)**. Interestingly, both TGF-β1 and CMDV modulated genes involved in ECM remodeling, including collagens, integrins, and metalloproteases. This suggests a potential reorganization of cell–cell and cell–matrix interactions, directing endothelial cells toward a more mesenchymal-like phenotype.

We validated the expression of the *SNA1* (Snail), *TWIST1* and *VIM* (vimentin) genes observed in the RNAseq analysis by RT-qPCR, using ACTB as housekeeping gene. The results revealed that CMDV increased the transcription of those three genes after 48 hours **(Figure 4A)** and 120 hours of treatment **(Figure 4B)**.

**Figure 4.**
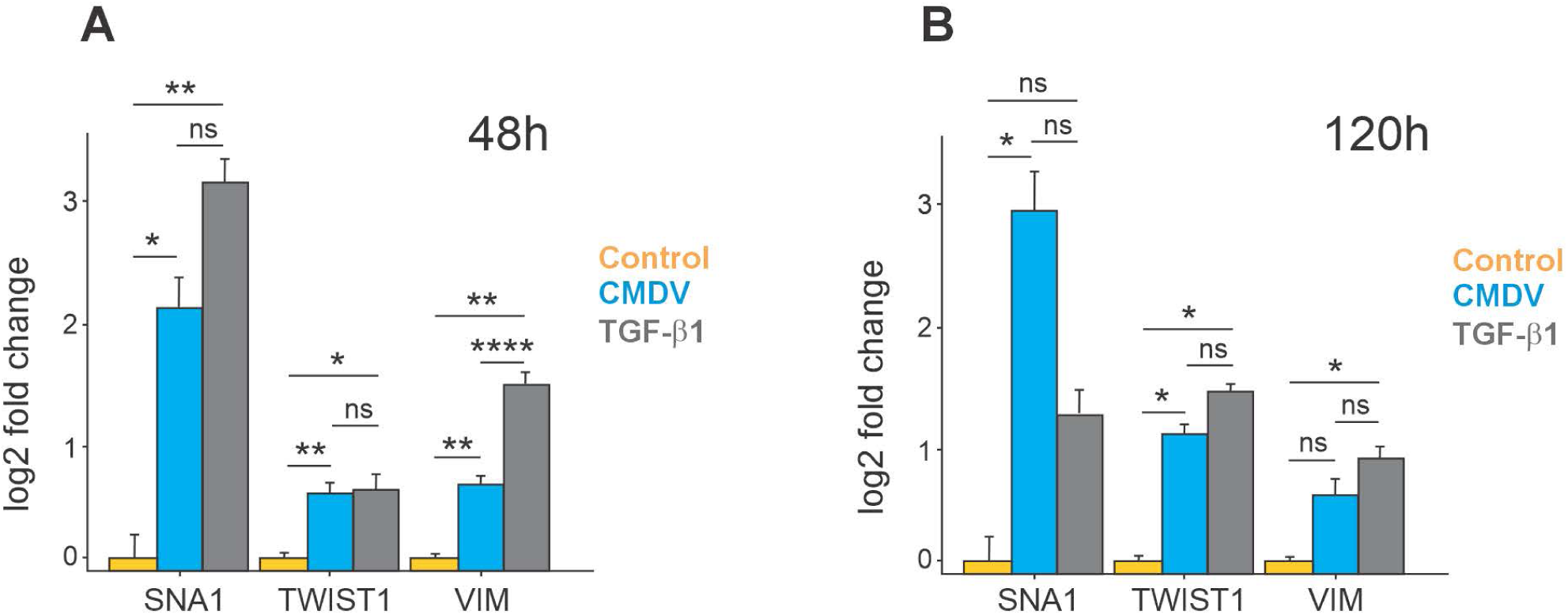
Semi-quantitative PCR of SNA1, TWIST1, and VIM gene expression at 48 hours (A) and 120h (B) post-treatment with CMDV and TGF-β1. Data is relative to untreated controls. Results are presented as mean ± SEM of three independent experiments, and statistical significance was determined using T test or Wilcoxon test (p<0.05). Significance between samples against untreated control is shown on top of each comparison. *p < 0.05, **p < 0.01, ***p < 0.001, ****p < 0.0001.

We then performed GO and pathway enrichment analysis on the RNAseq data **(Supplementary Figure 4)**. The main biological process (BP) induced after 48h treatment with CMDV was immune response **(Supplementary Figure 4A)**. Molecular functions (MF) clustered around receptor ligand activity and cytokine pathways, but the Cellular component (CC) was not significant. In contrast, the main biological processes triggered by 48h treatment with TGF-β1 were cellular migration and differentiation, with cellular components (CC) involved in cytoskeleton-extracellular matrix and adhesion. Interestingly, the major alterations to MF induced by TGF-β1 at 48h were calcium ion binding and sensory perception of mechanical stimulus **(Supplementary Figure 4C)**.

The main BPs induced by 120h treatment with CMDV were mitosis, including mitotic sister chromatid segregation, and cellular differentiation **(Supplementary Figure 4B)**. Meanwhile, CC were linked to cytoskeleton and cellular adhesion, with emphasis on focal adhesions **(Supplementary Figure 4B)**. This pattern was similar to that induced by TGF-β1 at 120 hours **(Supplementary Figure 4D)**. The CC and MF induced by 120h of TGF-β1 were associated with cellular morphology changes involving the cytoskeleton, tubulin binding and related components. In summary, CMDV induces an initial pro-inflammatory response that turns into an endothelial repair response at later time points, whereas TGF-β1 purely induces trans-differentiation with a late repair response that is similar to that promoted by CMDV.

### 2.3. Common cellular alterations induced by CMDV at 48h and 120h

The endothelial response induced by CMDV at 48 hours is different from that at 120 hours, although some common genes were identified at both timepoints. This suggests potential biological processes linking the two cellular responses associated to the effects of soluble factors on the cells over time. To assess this hypothesis, the DEGs from CMDV at both 48 hours and 120 hours were used to create a protein-protein network for each time-point **(Supplementary Figure 5A-B)**. At 48 hours, four main clusters were identified: one was disconnected from the others, and related to cellular differentiation, while the other three were interacting with each other and were associated with the immune response. At 120 hours **(Supplementary Figure 5B)**, 57 clusters were found, including large clusters that control cell cycle progression **(Supplementary Figure 5B, dashed red box)**. Additionally, one cluster was associated with cytokine signaling and several neighbor clusters were involved in ECM deposition and cell migration processes **(Supplementary Figure 5B, dashed blue ellipse)**. These observations were made using two independent cluster enrichment tools, StringdB and Enrichr.

To evaluate the possible connection between the three clusters identified at 48 hours and the cytokine signaling cluster along with its neighbors at 120 hours, a new interactome was created using the upregulated and downregulated genes with a p-value <0.05 in Cytoscape and StringDB. The merged network **(Supplementary Figure 6)** confirmed the connection between the two cellular responses, with four overlapping genes. The average local clustering coefficient (0.7) indicated that these interactions were not random. Notably, IL6 and FN1 emerged as central nodes. Furthermore, the MCL algorithm from String DB revealed additional clusters in the merged network. These clusters could be grouped into three major biological processes: immune/inflammatory response (cytokine-mediated signaling pathway), cell migration (positive regulation), and differentiation (neural crest cell migration). These processes were highly significant, with a False Discovery Rate (FDR) <6.76e-07, although the strength of the signals varied. Molecular functions were consistent with immune functions (growth factor binding, cytokine activity, and CXCR chemokine receptor binding, with high signal-to-noise ratio, strength and an FDR <3.5e-07). Cellular component analysis identified components involved in cellular migration and differentiation (endoplasmic reticulum lumen, collagen-containing extracellular matrix, and extracellular matrix, also with a high signal-to-noise ratio, variable strength, and FDR <2.5e-09).

Overall, these in silico observations were consistent with cellular and molecular assays, in which we observed alterations in markers related to morphological and cellular plasticity. Such consistency allowed us to select this network for further modeling.

### 2.4. Building a Non-Directed Asynchronous Model of the effect of CMDV on endothelial cell behavior (NDAM-CMDV)

The interactome described earlier was used to formalize a set of Boolean rules (Supplementary Table 1) regarding the fluctuation in expression and connections, mostly based on bibliographic data due to the scant experimental data available **(Figure 5)**. This section is described in technical detail in the Supplemental Section. Briefly, the resulting network consists of 41 nodes and 99 edges. Centrality analysis (Supplementary Table 3A and **Figure 5**) identified IL6 and FN1 as key nodes, with other relevant nodes were CXCL1, IL1α, FGF2, and CCL2. Since the ultimate goal of this model is enable simulations that reproduce the real situation, we implemented thresholds and modulators that modify the network generally or locally [33]. The number of iterations that would be required to reach a minimum threshold of FN1 degradation were determined using a first-order modified model with regular catabolic degradation rate [34]. FN1 required 23 iterations to reach the minimum threshold. The maximum thresholds and modulators are 24, establishing 23 FN1-dependent interactions. This was used to induce changes in specific nodes. The only exception was IL1R2, which required a threshold of 22 iterations to remain in the model.

**Figure 5.**
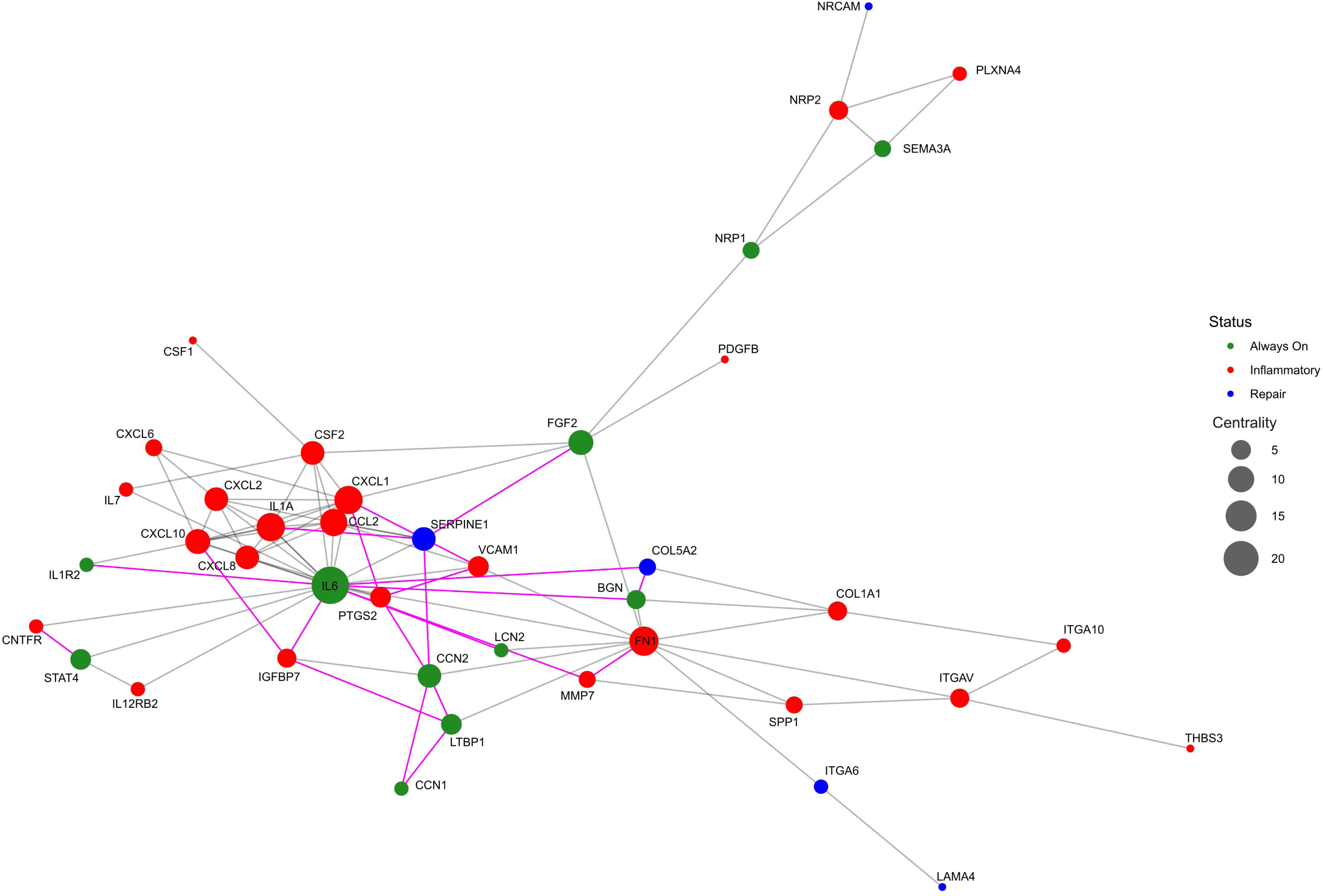
Graphic representation of the non-directed model depicting the effect of CMDV. The network depicts the nodes found in the transcriptomic analysis of CMDV-treated cells. Nodes are color-coded as inflammatory only (red), reparative only (blue) and always on (green). The size of the nodes represents centrality (node size) as indicated, with focus on IL6 and FN1. Gray lines represent edges. Magenta lines represent new interactions.

Additional regulators that influence the network in initial stages are defined by the transcription, translation and/or metabolic synthesis rates. For modeling, transcription and translation are assumed to be rapid and comparable to each other [35]. Conversely, metabolic synthesis involves multiple steps of different duration. This is best illustrated by VCAM1, an adhesive receptor produced in response to inflammation [36, 37]. This introduces an asynchronous behavior component in the system, which better simulates the experimental results and rounded up a new non-directed asynchronous model of the effect of CMDV on endothelial cells (NDAM-CMDV).

### 2.5. NDAM-CMDV robustness, accuracy and attractors

Next, we examined NDAM-CMDV accuracy and robustness. We performed 25000 simulations, each consisting of 100 iterations with asynchronous activation **(Figure 6 and Supplementary Table 4)**. NDAM-CMDV exhibited the expected behavior: pro-inflammatory nodes were activated during the first 25 iterations and then turned off; the remaining 75 iterations corresponded to the reparative/plasticity response.

**Figure 6.**
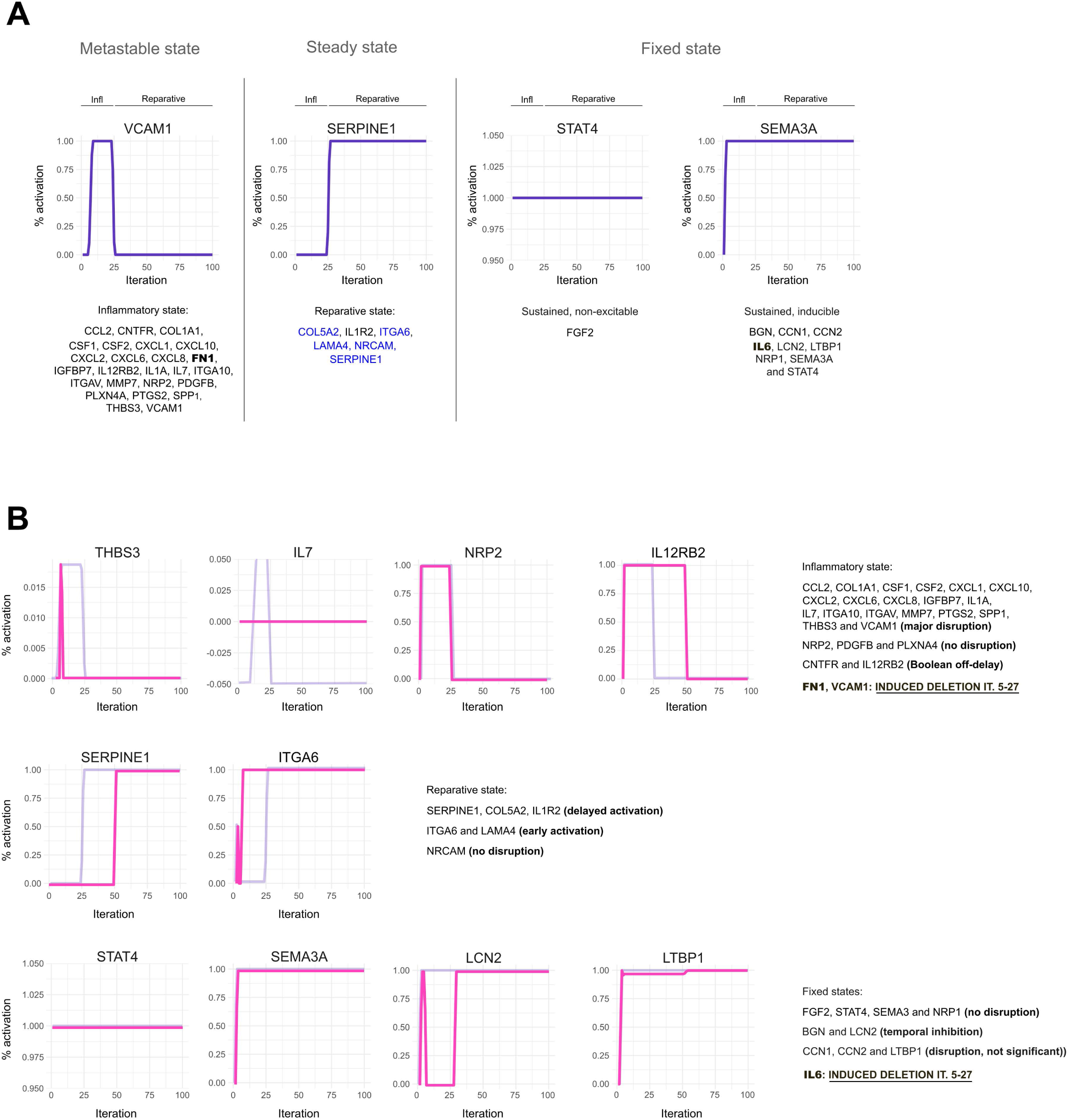
Model simulations predict gene responses in response to CMDV. A. Simulations of the nodes illustrated in Figure 5 display four main behaviors: inflammatory nodes are activated from the beginning of the simulation until approximately iteration 25, decaying thereafter (metastable state). Reparative genes are induced around iteration 25 and remain elevated for the rest of the iterations (steady state). Some genes, e.g. STAT4 and FGF2 do not become modified in response to stimulation (fixed, non-responsive), whereas other genes are induced early and remain elevated over the entire course of the simulation. B. Simulation of the effect of knocking out FN1, IL6, and VCAM1 nodes. Pale magenta lines represent the state of each node in untreated conditions (as in A). Steady state is reached after 50 iterations.

The activation percentage, media, and standard deviation from each node was highly predictable because they exhibit high activation but low variance (STAT4, FGF2, IL6, NRP1, SEMA3A, BGN, and LCN2). This means these nodes are consistent across the entire system. Another subgroup of nodes displays low means and variance (IL1A, CCL2, CSF2, CSF1, and IL7). Finally, another high predictability group includes those with high means and medium variance (CCN1, CCN2, LTBP1, and IL1R2), indicating that they are activated and tend to be consistent. However, other nodes tend to be more unpredictable due to their high variance and average means (SERPINE1, COL5A2, NRCAM, LAMA4, and ITGA6), or high variance and low means (IL12RB2, NRP2, CNTFR, PDGFB, FN1, and PLXNA4). Every node (gene) fell into four possible categories as shown in **Figure 6A**: i) induced during the inflammatory response (iterations 1-25); ii) induced only during the reparative response (iterations 26-100); iii) non-responsive; and iv) induced during the inflammatory response and sustained during the reparative phase.

On the other hand, we also determined the system attractors (Supplementary Table 5). The analysis identified them using various approaches, including synchronic and asynchronous methods, thereby identifying synchronic and asynchronous attractors. In the first two analyses, attractors were consistent (STAT4, IL6, FGF2, NRP1, SEMA3A, LCN2, BGN, ITGA6, LAMA4, LTBP1, CCN1/2, SERPINE1, IL1R2, COL5A2, and NRCAM), suggesting that these nodes influence the dynamic behavior of the system. However, this analysis identified 31 highly recurrent nodes. This strongly indicates that the model may be misaligned. Nevertheless, eight attractors (IL6, SEMA3A, LCN2, BGN, LTBP1, CCN1, CCN2, and IL1R2) were common in all the analyses. These indicate that the network has a moderate degree of complexity, high stability, and predictability, which are strong indicators of robustness. Additionally, the accuracy of the CMDV model compared to the experimental data is good, indicating its potential usefulness to study endothelial dysfunction in dengue using a cellular approach.

### 2.6. NDAM-CMDV can be used to predict endothelial responses to Dengue infection in silico

To test the performance of NDAM-CMDV under different conditions, we introduced observer-directed perturbations involving specific nodes in iterations 5-27. In the first perturbation, we deleted PTGS2 **(Supplementary Figure 7A)**. PTGS2 suppression caused a partial inhibition of the pro-inflammatory response, abrogating the induction of CCL2, CSF1, CSF2, CXCL10, CXCL8, IGFBP7, IL1A, IL7, and VCAM1. It also decreased the levels of CXCL10, CXCL8, IL1A, CSF1/2, IGFBP7, and IL1R2 **(Supplementary Figure 7A** and Supplementary Table 6).

Next, we deleted IL6 **(Supplementary Figure 7B** and Supplementary Table 7). This perturbation prevented the induction of CCL2, COL1A1, CXCL1, CXCL2, CXCL10, CXCL6, CXCL8, IGFBP7, IL1A, IL7, ITGA10, ITGAV, PTGS2, VECAM1, THBS3, and SPP1, while CSF1, CSF2, and MMP7 displayed a brief activation that declined rapidly. Additionally, IL6 deletion impairs expression of BGN, LCN2, IL1R2, SERPINE1, IL12RB2, and CNTFR.

Next, we deleted BGN and LCN2 simultaneously (data not shown). We found that this perturbation does not greatly affect the system. The main defect was a delay in the induction of COL5A2, IL1R2 and SERPINE2 until iteration number 50, explaining why their activation percentage decreased. Meanwhile, IL12RB2 and CNTFR increased because they remain present in the system until iteration 50.

We also tested the effects of a triple deletion scenario affecting IL6, FN1, and VCAM1 **(Figure 6B** and Supplementary Table 8). Simulations predicted a complete inhibition of CCL2, CXCL10, IGFBP7, IL1A, and IL7 expression, with brief activation of COL1A1, CSF1, CSF2, CXCL1, CXCL2, CXCL8, CXCL6, FN1, ITGA10, ITGAV, PTGS2, THBS3, MMP7, and SPP1. The triple knockout disrupted the behavior of BGN and LCN2. Their expression outside the system recovered after iteration 27. CCN1, CCN2, and LTBP1 displayed decreased activation that is restored after iteration 27, while CNTFR, IL12RB2, IL1R2 and SERPINE1 behaved as reported when only IL6 was deleted **(Supplementary Figure 7B** and Supplementary Table 7). Conversely, ITGA6 and LAMA4 activated faster than in control conditions.

In addition, we deleted FN1 but overexpressed FGF2 **(Supplementary Figure 7C** and Supplementary Table 9). CCL2, IL7, and VECAM1 were inhibited, whereas COL1A1, CSF1, CSF2, ITGA10, MMP7, SPP1, and THSB3 displayed a brief activation. Additionally, CCN1/2, IL1R2, ITGAV, and LTBP1 were modestly reduced at the beginning, recovering thereafter. Similar to the previous simulation, ITGA6 and LAMA4 were activated early.

Finally, we analyzed the effect of a single node knockout **(Figure 7A)** and single node overexpression **(Figure 7B)**. FGF2 knockout increased the levels of VCAM1, IL7, CXCL1, CCL2, CXCL10, CXCL8, IL1A, CXCL2, PTGS2, CXCL6, CSF1, CSF2, NRP2, PLXNA4, FN1, COL1A1, ITGAV, THBS3, ITGA10, SPP1, MMP7, IGFBP7, and PDGFB, while it decreased the levels of NRCAM, COL5A2, LAMA4, ITGA6, and NRP1. Additionally, we found that most nodes exhibit a similar expression profile, except for NRP1.

**Figure 7.**
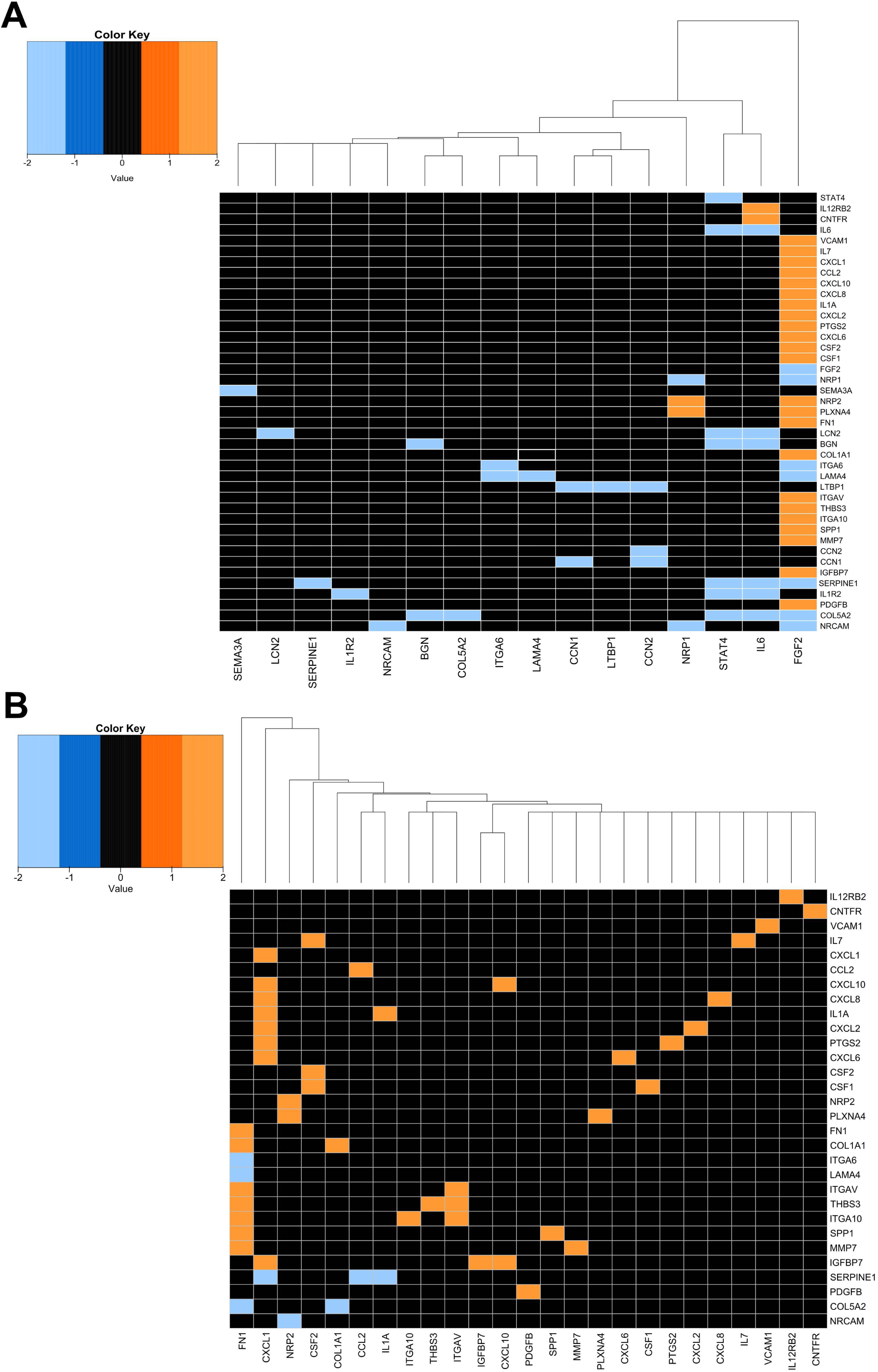
Hierarchical clustering of perturbation analysis of the effects of a single mutated node in the CMDV network. (A) Knockout of each node and their closeness. FGF2 knockout increases the expression of pro-inflammatory nodes. (B) Overexpression of each node and their closeness. FN1 enhances expression in some pro-inflammatory nodes such as MMP7. This is similar for CXCL1, but its effect is local and modifies the CXCL cluster.

Conversely, overexpression of FN1 increases MMP7, COL1A1, ITGAV, THBS3, ITGA10, and SPP1, while reducing COL5A2. Additionally, CXCL1 enhances IGFBP7, CXCL6, PTGS2, CXCL2, IL1A, CXCL8, and CXCL10, while decreasing PDGFB. Most nodes exhibit similar expression levels, although there are slight differences. Interestingly, NRP2 and CSF2 share similar expression patterns, while FN1 and CXCL1 are related, yet display completely different expression profiles compared to the other nodes. Increasing the sensitivity of the analysis decreased variation and masked mutational effects per node **(Supplementary Figure 7D-E)**.

## 3. Discussion

Soluble factors released during DENV infection induce a transient and partial EndMT-like state in endothelial cells. In our model, exposure to CMDV caused morphological changes, intercellular junction disruption, and upregulation of some EndMT markers, leading to increased endothelial permeability [16]. Notably, this response was biphasic and reversible: at 48 hours, CMDV-exposed cells displayed a strong pro-inflammatory profile, whereas by 120 hours they had shifted toward an endothelial repair and angiogenic profile, similar to what has been reported for EMT [38]. This pattern (an initial inflammatory surge followed by a possible reparative phase) may explain why endothelial dysfunction in severe dengue is often transient and does not always lead to irreversible vascular injury [9, 11, 12]. We posit that CMDV triggers an acute, partial EndMT that compromises the endothelial barrier during the peak infection period that may be naturally resolved in the absence of additional inflammatory stimuli and/or additional factors that sustain inflammation or exacerbates the repair response debilitating endothelial cell junctions. Importantly, the four serotypes of dengue produce the same disease; the difference is in the onset of the symptoms or their intensity [39–42]. While disease produced by any serotype can evolve to SD, DENV2 has been extensively associated with this complication and a higher mortality rate [43].

These findings align with and extend previous observations of dengue pathogenesis. Prior studies have shown that pro-inflammatory cytokines and the viral NS1 protein can disrupt endothelial junctions and increase vascular permeability [6, 10–12]. Our results support this framework and further indicate that other soluble mediators present during infection (beyond NS1, which is not completely absent from media in similar conditions [44]) also play a critical role in endothelial dysfunction.

The analysis of dengue virus–induced cytokines in our model reveals a strong pro-inflammatory signature—dominated by IL-6, CXCL1, CXCL2, CXCL8, and CCL2—which mirrors the central role these mediators play in endothelial activation and vascular barrier disruption. Although NS1 was not directly measured in this specific set of experiments, its presence in DVC is expected due to its well-documented secretion and persistence during DENV infection [10]. Secreted flavivirus NS1, including DENV2 NS1, can directly alter endothelial permeability in vitro in a cell type-dependent manner [45], experimental evidence suggests this effect is largely independent of inflammatory cytokines in HMEC-1 cells, as they do not produce TNF-α, IL-6, or IL-8 upon NS1 exposure alone. Instead, NS1 primarily acts via the activation of glycocalyx-degrading enzymes such as sialidases, cathepsin L, and heparanase [46].

Clinically, the relationship between specific cytokines and severe dengue remains complex. While reports for TNF-α, IL-1β, and CXCL8/IL-8 are inconsistent, IL-6 levels are consistently associated with a pathogenic role in patients [10, 11]. Furthermore, high levels of NS1 in plasma do not always correlate temporally with vascular leakage in vivo, often appearing after NS1 levels have declined [46]. Our model suggests that the synergistic effect of infection-induced pro-inflammatory cytokines and NS1 may act through converging pathways to promote junctional remodeling and vascular dysfunction [10].

Unlike the permanent EndMT seen in chronic fibrotic diseases or cancer [17, 47], the partial EndMT-like changes induced by CMDV appear incomplete and transient. Endothelial cells acquired some mesenchymal traits (e.g., elevated Snail, N-cadherin, vimentin) without fully losing their endothelial identity [48–52]. This suggests a partial EndMT that may facilitate immune cell recruitment, rather than leading to irreversible mesenchymal conversion. In fact, the early secretion of chemokines (CXCL1, CXCL2, CCL2) by CMDV-treated cells would promote leukocyte adhesion and transmigration, consistent with cytokine-driven EndMT in inflammatory settings [53–56] and the known role of these chemokines in neutrophil recruitment [57, 58].

Mechanistically, CMDV appears to activate both inflammatory signaling and mechanotransduction pathways in endothelial cells. The surge of IL-6 and other cytokines at 48 h indicates activation of canonical inflammatory cascades (e.g., JAK/STAT) that drive endothelial activation and EndMT [49, 59, 60]. In parallel, we observed upregulation of TWIST1 and transient focal adhesion disassembly, implicating pathways associated with shear stress responses [47–49, 61]. It is possible that CMDV-induced extracellular matrix degradation and cytoskeletal remodeling provide cues resembling mechanical stress, thereby promoting a migratory, EndMT-like phenotype. By 120 h, as expression of inflammatory stimuli declines, endothelial cells upregulate genes related to cell proliferation and tissue repair [50–52, 54], which may eventually lead to the normalization of the cells. Whether such return to normalcy is due to the transcriptomic changes seen in this phase or by the absence of stimulation in the media remains to be investigated.

This transient, partial EndMT-like burst – followed by eventual recovery – may be an adaptive mechanism to limit long-lasting vascular damage [24]. Computational modeling further underscores IL-6 and fibronectin (FN1) as critical drivers of the effect of the CMDV. In silico removal of either diminished the inflammatory and ECM responses. This correlates well with the crucial role of IL-6 in endothelial dysfunction [16, 59] and FN1 in maintaining vascular integrity [53]. Targeting IL-6 could attenuate the early cytokine-mediated damage; indeed, IL-6 inhibitors such as tocilizumab merit exploration in severe dengue [55]. Likewise, preventing FN1-driven extracellular matrix breakdown might help preserve endothelial junctions and barrier function [53]. Another potential strategy is to stabilize endothelial cells during infection. We have shown that the tyrosine kinase inhibitor imatinib partially restored endothelial junction integrity after CMDV exposure. Imatinib has protective effects on the endothelium during DENV infection by stabilizing VE-cadherin and the actin cytoskeleton [14]. Therefore, imatinib could address both the inflammatory and mechanical aspects of dengue-induced endothelial dysfunction, potentially yielding a synergistic benefit [14, 55]. Beyond these specific examples, our integrative approach provides a framework to screen additional compounds: by targeting key nodes like IL-6 or FN1 in the network, future studies can identify and test novel interventions to mitigate dengue-associated vascular leakage.

This study has identifiable limitations. The in vitro model used here, while informative, cannot fully replicate the complexity of DENV infection in vivo. In vivo, endothelial cells are influenced by interactions with immune cells and hemodynamic forces, which were absent in our cell culture system [17, 26]. Second, while the effect of CMDV on vascular leakage can be inferred from previous studies, including our own [14–16], we did not measure endothelial permeability in real time. Finally, the Boolean network model is an abstraction that would benefit from incorporation of kinetic data and further experimental validation to improve its predictive power [33, 56]. As it is, it is a useful hypotheses generator that provides a justifiable conceptual framework for future prediction, investigation and experimental validation in vitro and in vivo. Future studies, including animal models and patient samples, are required to confirm the transient partial EndMT-like mechanism *in vivo* and to assess the proposed therapeutic strategies under physiological conditions. In this context, identifying IL-6 and FN1 as key drivers of this process should be an experimental priority that would situate these two genes as concrete targets for therapies aimed at preserving vascular integrity in severe dengue. In conclusion, our data and model sustain that soluble factors from dengue infection promote a reversible, partial EndMT-like mechanism in endothelial cells, linking the acute inflammatory surge to transient vascular leakage. These findings advance our understanding of dengue pathophysiology and lay the groundwork for targeted interventions to prevent life threatening vascular complications. Moreover, the concept of a regulated, transient EndMT-like process may have broader relevance for other acute inflammatory conditions involving endothelial dysfunction, such as sepsis [57, 58].

## 4. Materials and Methods

### 4.1. Cell lines and viral infections

Human Microvascular Endothelial Cells (HMEC-1) (ATCC Cat# CRL-3243) were maintained at 37°C with 5% CO2 in RPMI supplemented with 10% FBS, 10mM L-glutamine, 100 U/mL penicillin/100 mg/mL streptomycin (P/S), 10ng/mL Epidermal Growth Factor (hEGF), and 1 µg/mL Hydrocortisone. Cells were used up to passage 10. Human Epithelial Kidney Cells (HEK-293) (ATCC Cat# CRL-1573) were maintained at 37°C with 5% CO2 in RPMI supplemented with 10% FBS, 10mM L-glutamine, 100 U/mL penicillin/100 mg/mL streptomycin (P/S), in the same way that HeLa Cells (ATCC Cat# CCL2), but in DMEM media. Aedes albopictus clone C6/36 HT cells (ATCC Cat# CRL-1660) were cultured at 34°C with 5% CO2 in L-15 media supplemented with 10% FBS and 1X P/S. Dengue virus Serotype 2 (DENV-2, New Guinea) was amplified in C6/36 HT cells grown in L-15 medium supplemented with 10% FBS, 100 U/mL P/S in T75 flasks at 70% confluence and incubated at 34°C [14]. The virus was titrated in BHK-21 hamster kidney cells (ATCC Cat# CCL-10) as described [14, 62].

### 4.2. Generation of conditioned media

CMDV was obtained using the protocol described in [13]. Briefly, HMEC-1 cells at 75% confluence were infected with Dengue virus serotype 2 (DENV-2) obtained from viral amplification in C6/36 cells and titrated in BHK-21 cells as described [16]. HMEC-1 infection was carried out at a relative Multiplicity of Infection (MOI) of 5 for 2 hours. Subsequently, the cells were rinsed with Phosphate-Buffered Saline (PBS) and cultured in RPMI supplemented with 2% Fetal Bovine Serum (FBS) for 48 hours. Cell culture supernatants were collected and transferred to Petri dishes for complete virus inactivation using ultraviolet (UV) irradiation for 15 minutes. UV inactivation was confirmed by the absence of cytopathic effects in HMEC-1 cells for 6 days [16] and lack of PFU in BHK-21 cells. Inactivated supernatants were stored at –80°C for further use [14]. For experimental treatments, a ratio of 80:20 was used (80% CCM or CMDV and 20% fresh medium).

### 4.3. Fluorescence microscopy and image quantification

For the 48-hour assays, 1.5×10^5^ HMEC-1 cells were seeded into glass coverslips previously treated with 4% porcine gelatin and cultured for four days in 2% FBS + RPMI. Cells were then treated with CMDV in an 80:20 ratio [14], with conditioned medium from healthy cells (negative control), or with 5 ng/mL TGF-β1 (positive control). Cells were re-stimulated after 24 hours, then fixed after 48 h from the beginning of the assay. All conditions maintained a final concentration of 0.1% FBS in RPMI media.

For the 120-hour and imatinib assays, 4.0×10^4^ cells were seeded as above. After 24h, medium was switched to 0.1% FBS to favor the formation of cell-cell junctions and cells remained in these conditions for another 48h. Then, cells were treated with CMDV in the presence or absence of 6.25 µM imatinib (this concentration was determined experimentally as outlined in [14]) and fixed after 48h (120h total time).

After the cells were exposed to CMDV per 48 h or 120h, or to CMDV and then to imatinib per 48h or 120h, they were washed three times with 1× Cytoskeletal Buffer with sucrose (CBS)[14], thereafter, they were fixed with 3.8% paraformaldehyde (PFA) in CBS for 20 minutes at 37°C. The cells were treated with NH4Cl (50 mM), permeabilized (0.02% Triton), and blocked (PBS + 5% FBS) for 1 hour at 37°C. Subsequently, the fixed cells were labeled at a 1:400 dilution with primary antibodies (mouse anti-SNAIL/VE-Cadherin /N-Cadherin or Occludin (OCLN)) for 1 hour at 37°C. After three thorough PBS washes, the cells were incubated with species-matched secondary antibodies coupled to AlexaFluor-594 or AlexaFluor488 as indicated, and Hoechst (1:5000) for 45 minutes at 37°C and finally mounted on slides using FluorSave mounting medium. For image acquisition, we used a Leica Thunder Tissue Analyzer (Leica Microsystems, GmbH) microscope equipped with a multi-LED using a 63x objective. Data processing and quantification were performed using ImageJ. To minimize observer bias, all image files were randomized and blinded before analysis. Data processing was performed using ImageJ version 1.54f, where the Integrated Density (IntDen) for both red and green channels was recorded. To maintain data quality, an outlier removal step was implemented using the Interquartile Range (IQR) method (values outside Q1-1.5\times IQR and Q3+1.5\times IQR were excluded). Final fluorescence intensity values were expressed as a percentage of expression relative to the total intensity to account for variations in cell density across treatments. Analysis was conducted using R software (version 4.3.0) with at least three biological replicates per condition. Given that the data did not meet the assumptions of normality (Shapiro-Wilk test) or homogeneity of variance (Bartlett’s test), non-parametric Kruskal-Wallis test was used, followed by a Dunn’s post-hoc test with Bonferroni correction. Multiple comparisons were controlled using Benjamini–Hochberg where relevant.

### 4.4. RNAseq-bulk sample generation and processing

HMEC-1 cells were seeded at 5×10^5^ per well (6-multiwell plate) in 10% RPMI at 37°C with 5% CO2 overnight to promote adherence. The cells were treated with 5 ng/mL TGF-β1 (positive control), CMDV, or conditioned medium from untreated cells (control) at a final concentration of 0.1% FBS. Medium was changed every 24h, and cells were collected for RNA extraction at the indicated time points. Three technical replicates and three biological replicates were performed per assay. Total RNA extraction was performed using the easyRNA kit (QiaGen) according to the manufacturer’s protocol. RNA quantity was assessed using Nanodrop and Qubit. Additionally, the quality was assessed according to the RNA Integrity Number (RIN) using Tapestation 4150 (Agilent) to select samples with a RIN ≥7 for library preparation. Library construction was carried out using MGI technology (MGISP-100 system, MGI. Shenzhen, China) in accordance with the manufacturer’s instructions and paired-end sequencing (2×100 bp). High-throughput sequencing was performed using the DNBSEQ-G50RS system (MGI, Shenzhen, China) with an SE 100 flowcell, spanning 120 cycles. The sequencing was performed at the National Genomics Laboratory of Colombia. Sequencing was conducted to achieve a depth of approximately 40 million reads per sample.

### 4.5. Sequencing Data Processed and Differential expressed genes (DEGs) analysis

Raw sequencing reads were processed to remove adapters and low-quality bases using Trimmomatic v0.39 [63] with the settings LEADING:3, TRAILING:3, MINLEN:50 CROP: 92,HEADCROP:8, SLIDINGWINDOW:4:20. After trimming (Trimmomatic) and read-level QC (FastQC), we assessed sample-level QC using variance-stabilized counts (DESeq2) and performed principal component analysis and hierarchical clustering to confirm replicate concordance and to identify potential outliers. No samples met exclusion criteria based on low library complexity or abnormal mapping/QC metrics. Quality control of the trimmed reads was performed with FastQC v0.11.9 [64], and clean reads were aligned to the reference genome GRCh38 (GCA_000001405.15 GCF_000001405.26) using STAR v2.7.10a [65] with the parameters –-outFilterType BySJout –-outFilterMultimapNmax 20 –-alignSJoverhangMin 8 –-alignSJDBoverhangMin 1 –-outFilterMismatchNmax 999 –-outFilterMismatchNoverLmax 0.04 –-alignIntronMin 20 –-alignIntronMax 1000000 –-seedSearchStartLmax 30. Gene-level read counts were quantified using featureCounts v2.0.1 [66] with annotation files from GENCODE v39 [67]. Differential expression was performed with DESeq2 v1.36.0 [68]; p-values were adjusted using the Benjamini–Hochberg procedure and genes were considered significant at FDR < 0.05 with |log2FC| ≥ 1. All downstream RNA-seq-derived gene sets used for modeling were restricted to this FDR-controlled list. Data visualization, including heatmaps and volcano plots, was performed using the R package ggplot2 v3.3.6.

### 4.6. Gene ontology

Results from DEGs were used for functional enrichment analysis utilizing the Enrichr web tool version 2016 update [69]. We used Gene Ontology (GO) annotation to identify biological process enrichment, molecular functions, and cellular components associated with these DEGs. Only up regulated genes with p<0.05 were used, and the accepted match are those with a adjusted p-value<0.01 and the combined score was high because the odds ratio varied for each suggested biological process, pathway or function. Additionally, the DEGs from CMDV at 48h and 120h were used to identify potential pharmacological targets through The Drug Gene Interaction Database. The results were filtered based on the interaction score (>4) and FDA approval. Additionally, the DEGs from CMDV at 48h and 120h were used to perform an interactome analysis in the StringDB database. The following parameters were applied: H. sapiens as the biological model, a high confidence level of 0.9, 1% FDR stringency, and clustering using MCL with an inflation factor of 3, which were classified in three biological phenomena: immune/inflammatory response, cellular migration and cellular plasticity/differentiation. The non-interacting nodes were excluded. The merge network between the two cellular responses was isolated and analyzed in cytoscape v 3.10.1, using Dynet Analyzer plugin to detect overlaps.

### 4.7. RT-qPCR

Expression levels of SNA1, TWIST,1 and VIM were assessed. ACTB (beta-actin gene) was used as housekeeping control. Primer sequences used are shown in Table 1. cDNA was synthesized using M-MLV reverse transcriptase (Promega) from 500 ng of starting RNA. The reaction mix included random primers at a concentration of 125 ng/µL, 10 mM dNTP Mix, 50 mM Mg2+ 5x First-Strand Buffer, 100 nM DTT, RNase Out at 40 U/µL and MilliQ water, with a final volume of 20 µL. The thermal profile was as follows: 65°C for 5 minutes (incubation), 37°C for 2 minutes, 25°C for 10 minutes, 7°C for 50 minutes, and 70°C for 15 minutes (enzyme inactivation).

**Table 1.**
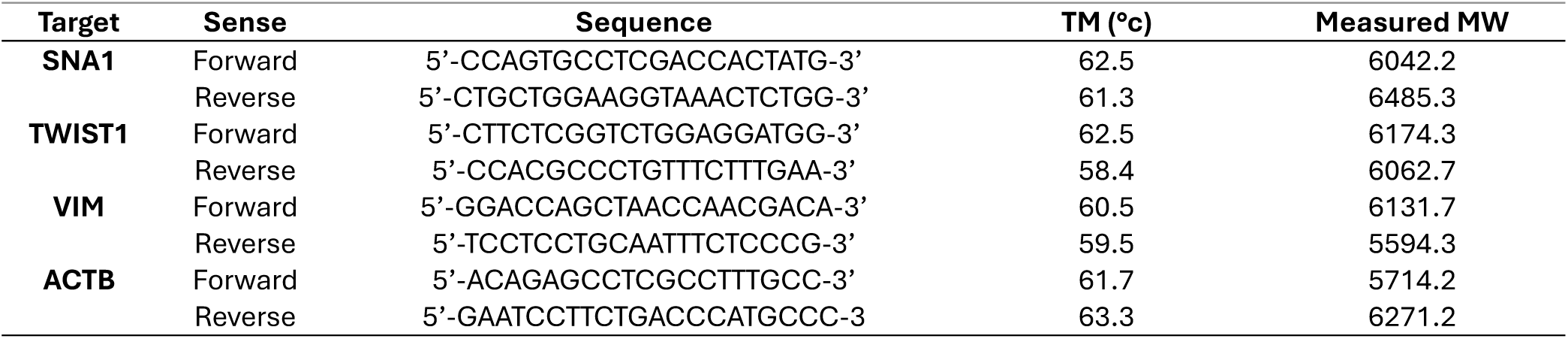
qPCR Primer sequences.

qPCR was conducted using SsoFastTM EvaGreen® Supermix (Biorad). Samples contained 400 nM primers, 1 µL of cDNA in a final volume of 9 µL. Amplification was performed in a CFX96TM TOUCH REAL-TIME PCR (Biorad) with the following thermal profile: 95°C for 30 seconds, followed by 40 cycles of 95°C for 5 seconds, 60°C for 5 seconds, and a gradual increase from 65°C to 95°C at a rate of 0.5°C every 5 seconds in an optical 96 multiwell plate. All reactions were performed in triplicate, and the mean Ct (Cycle threshold – fluorescent cycle threshold) was calculated for each sample.

mRNA differential expression was calculated using the 2−ΔΔCt (Livak method), where ΔCt represents the difference between the Ct of the gene of interest and that of the housekeeping control ACTB). ΔΔCt is the comparison between the treated (CMDV and TGF– β1) samples and the controls (C-). The results were expressed as Log2 (LogFC). An increase in expression is indicated when LogFC > 1 and a decrease when LogFC < –1, following the methodology described by Coebergh Van Den Braak [70]. Differential expression, statistical analysis and graphics were performed in RStudio, with the packages tidyr v.1.31, dplyr v.1.14, qpcR v.1.4-1, stringr v.1.5.1, purrr v.1.0.2 and qPCRtools v.1.0.1. Statistical tests were carried out after assessing distributional assumptions (Shapiro–Wilk for normality; Levene’s test for homoscedasticity). Because these assumptions were not met, we used non-parametric tests (Mann–Whitney or Kruskal–Wallis with Dunn’s post hoc). Multiple comparisons were controlled using Benjamini–Hochberg where relevant.

### 4.8. Non-directed Asynchronous Boolean Network

A subnetwork was obtained from the interactomes using the CMDVs DEGs created in StringDB to model a non-directed network. Therefore, bibliographic data was compiled to infer the node state where it was absent at one point and to understand its role in the biological processes and the model. For this reason, some interactions (edges) were included in the network to enhance its dynamism and to clarify the biological results.

The Boolean rules were formalized using Boolean algebra and were transcribed into a format suitable for use in the SPIDDOR v.1 RStudio packag. The number of edges and nodes, as well as centralities, the clustering coefficient, communities, and network metrics, were determined using the igraph v.2.1.2 package in RStudio v. 2024.12.0 (R v.4.3.0). To achieve the iterations necessary to activate the switch that changes between the 48-hour and 120-hour periods produced by the CMDV, a modified first-order decay model (1) was used [71], to find a solution to n (2).

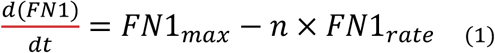

with the condition that FN1 ≥ FN1min. FN1 represents the amount of FN1 at a given iteration, FN1*max* is the maximum amount of FN1, n is the number of iterations and FN1rate is the degradation rate of FN1

The equation (1) can be solved as follows:

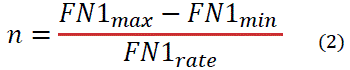

which is formulated considering the role of FN1 in the system and the biological process. The minimum and maximum values were arbitrary (0.1 and 1 (100%), respectively). Meanwhile, the reported degradation rate taken was 4.8% per 1h (it was taken as 0.04) which is the reported for normal catalysis [82]. The equation ignores the recovery rate. The advanced Boolean networks used in SPIDDOR defined the thresholds and modulators as described in Table 2.

**Table 2.** Thresholds and modulators in the advanced Boolean rules to describe the effect of CMDV on endothelial cells.

| Type | Name | Value |
| --- | --- | --- |
| **Thresholds** | THR_IL6 | 2 |
|  | THR_IL6 | 22 |
|  | THR_IL6 | 24 |
|  | THR_FGF2 | 24 |
|  | THR_NRP1 | 24 |
| **Modulators** | MOD_IL12RB2 | 1 |
|  | MOD_CNTFR | 1 |
|  | MOD_IL6 | 1 |
|  | MOD_VCAM1 | 3 |
|  | MOD_IL7 | 2 |
|  | MOD_CCL2 | 2 |
|  | MOD_CXCL10 | 2 |
|  | MOD_CXCL8 | 2 |
|  | MOD_IL1A | 2 |
|  | MOD_CXCL2 | 2 |
|  | MOD_PTGS2 | 2 |
|  | MOD_CXCL6 | 2 |
|  | MOD_CSF2 | 2 |
|  | MOD_SEMA3A | 1 |
|  | MOD_PLXNA4 | 1 |
|  | MOD_LTBP1 | 2 |
|  | MOD_THBS3 | 2 |
|  | MOD_ITGA10 | 2 |
|  | MOD_SPP1 | 2 |
|  | MOD_MMP7 | 2 |
|  | MOD_CCN2 | 2 |
|  | MOD_IGFBP7 | 2 |
|  | MOD_SERPINE1 | 24 |
|  | MOD_IL1R2 | 3 |
|  | MOD_COL5A2 | 24 |

The parameters to perform the average simulations, as well as to find the synchronous and asynchronous attractors and some interesting punctual perturbations such as knockouts or overexpression, were 100 iterations with 25000 repetitions in an asynchronous way. Meanwhile, to identify all the attractors, 6 CPUs were used with startStates set to 1000. The perturbation analysis evaluated the effects of single-node knockouts and single-node overexpression on the network. Sensitivity was disregarded due to its impact on the results. The model was exported to SBML (https://github.com/bioweaver/CMDV_nondirectModel_v.1.0). We deposited it in GitHub (https://github.com/bioweaver/CMDV_nondirectModel_v.1.0). Model visualization was created using ggplot v.3.5.0 packages.

### 4.9. Non-directed Asynchronous Boolean Network

For the microscopy assays, data normality was assessed using the Shapiro-Wilk test and homogeneity was evaluated using the Bartlett test. Depending on the data distribution, one-way analysis of variance (ANOVA) was performed for normally distributed data, while the Kruskal-Wallis test was used for non-normally distributed data. Post-hoc analyses were conducted using Bonferroni in Dunnett tests or Tukey tests as appropriate. For the qPCR validation data, T test or Wilcoxon test were performed depending on the data distribution. Additionally, the mean and standard error of the mean (SEM) were calculated. For the non-directed network, the mean and standard deviation were calculated for the simulations and perturbation analyses performed.

#### Node Selection and Edge Support

The nodes included in our model were prioritized from the differentially expressed genes (DEGs) identified at 48 and 120 hours post-exposure to CMDV. The selection and initial interaction mapping were performed using Cytoscape and STRING-db, with the latter providing the primary experimental and bibliographic support for each initial edge. While most nodes originate directly from our DEGs, some nodes presented “unknown” or inconsistent states at specific time points (e.g., at 48h vs. 120h). In these cases, we performed an extensive bibliographic search to determine the most probable biological state, ensuring the model remained consistent with the observed cellular response. Additionally, we manually verified interactions not initially detected by STRING-db or Cytoscape through literature review. These suggested interactions, which refine the molecular mechanisms affected by CMDV, are explicitly highlighted as predictions that require future experimental validation to increase the model’s robustness.

#### Biological Justification for the Asynchronous Model

We selected a non-directed asynchronous paradigm because it more accurately reflects real biological scenarios compared to synchronous models. In a biological system, it is extremely rare for all molecular events to occur simultaneously. Asynchronous modeling assumes that variables are updated one at a time, capturing the stochastic nature of cellular processes. This is particularly relevant for describing multi-stability—the simultaneous presence of multiple state-space attractors (e.g., epithelial vs. mesenchymal states)—which describes how a single viral stimulus can lead to different cellular fates. Then, biological functions operate on vastly different timescales (e.g., fast protein-protein interactions vs. slower protein synthesis). Asynchronous and stochastic updating help mitigate the temporal distortions of synchronous models, allowing the representation of hierarchical dependencies, such as the requirement for a protein to be synthesized before it can be phosphorylated. In this models “complex attractors” emerge as overlapping loops in the state graph. This complexity is better suited for capturing system-wide biological behavior and cellular plasticity, as the trajectory of these graphs defines stable states (attractors) that represent biological equilibrium under specific conditions [72, 73].

## Supplementary Materials

The following supporting information can be downloaded at: xxxxxxxxxxxxxx, Figure S1 and S2: 4.3 Immunofluorescence results from cells exposed to CMDV per 120h and then treated with imatinib per 48h or 120h. Figure S3: DEGs at 48h and 120h from CMDV and TGF-β1. Figure S4: Enrichment analysis of TGF-β1 at 48 hours and 120 hours. Figure S5: Interactomes from CMDV at 48h and 120h. Figure S6: Perturbation scenarios simulated in CMDV non-directed network. Table S1: Potential Pharmacological Targets detected in CMDV DEGs. Table S2: Genes State at 48h and 120h for CMDV model. Table S3: Centralities analysis results and Network Metrics. Table S4: CMDV network Simulation Average. Table S5: Attractors Analysis. Table S6: PTGS2 knockout scenario in CMDV network. Table S7: IL6 knockout scenario in CMDV network. Table S8: Triple Knockout simulation AVG in CMDV network. Tables S9: FGF2 overexpression and FN1 knockout scenario AVG.

## Acknowledgments

The authors thank María Elena Peñaranda and Eva Harris (University of California, Berkeley) for their generous donation of reagents. We also thank Engineer Hernández Talero C.A. for consulting and peer review of the code and the Ana I. García-Vega (Advanced Cellular Analysis Unit, CIC, Salamanca) for training and assistance with ImageJ code for image analysis.

## Author Contributions

Conceptualization, J.C.G-G, C.A.O-C, and M.V-M; methodology, J.P.A-G, C.A.O-C, G.J-F, B.A.R.-R, D.A.A-D, J.C.G-G and M.V-M; software, J.P.A-G, C.P.R-E, J.M.R-M, B.A.R-R, G.J-F and C.A.O-C; formal analysis, J.P.A-G, C.A.O-C, G.J-F, B.A.R-R, M.V-M and J.C-G-G; writing—original draft preparation, J.P.A-G, C.A.O-C, J.C.G-G. and M.V-M; writing—review and editing, J.P.A-G, C.A.O-C, C.P.R-E, D.A.A-R, J.M.R-M, G.J-F, J.C.G-G and M.V.-M; visualization, J.P.A.-G, C.A.O-C, C.P.R-E, J.M.R-M, G.J-F and M.V.-M; supervision, J.C.G.-G. and M.V-M.; project administration, J.C.G-G; funding acquisition, J.C.G-G. All authors have read and agreed to the published version of the manuscript.

## Data Availability Statement

CMVD non-directed network code: https://github.com/bioweaver/CMDV_nondirectModel_v.1.0

RNAseq sequences: https://dataview.ncbi.nlm.nih.gov/object/PRJNA1331842?reviewer=lu4r1nun9376f1c15sp7dekepv

The rest of the data, including original images, is available from the corresponding authors upon reasonable request.

## Supplemental information to

## Extended Results

### 2.4. Building a Non-Directed Asynchronous Model of the efect of CMDV on endothelial cell behavior (NDAM-CMDV)

The interactome described earlier was formalized into a set of Boolean rules (Table 1). To achieve this, we used a bottom-up approach to determine the expression of unknown genes at 48 hours or 120 hours (Supplementary Table 2). The data had to describe the function of the gene (node). Additionally, it considered the neighboring clusters and their interactions. Potential edge directionality was inferred from bibliographic data, which is reflected in the basic model rules. The rationale behind this choice is the limited amount of experimental data. Validation of the use of such approach as a fully predictive tool will require additional experimental validation to confer full significance to the edges. However, to ensure that the model corresponds to the experimental data, some new interactions between selected nodes were incorporated.

**Table 1.**
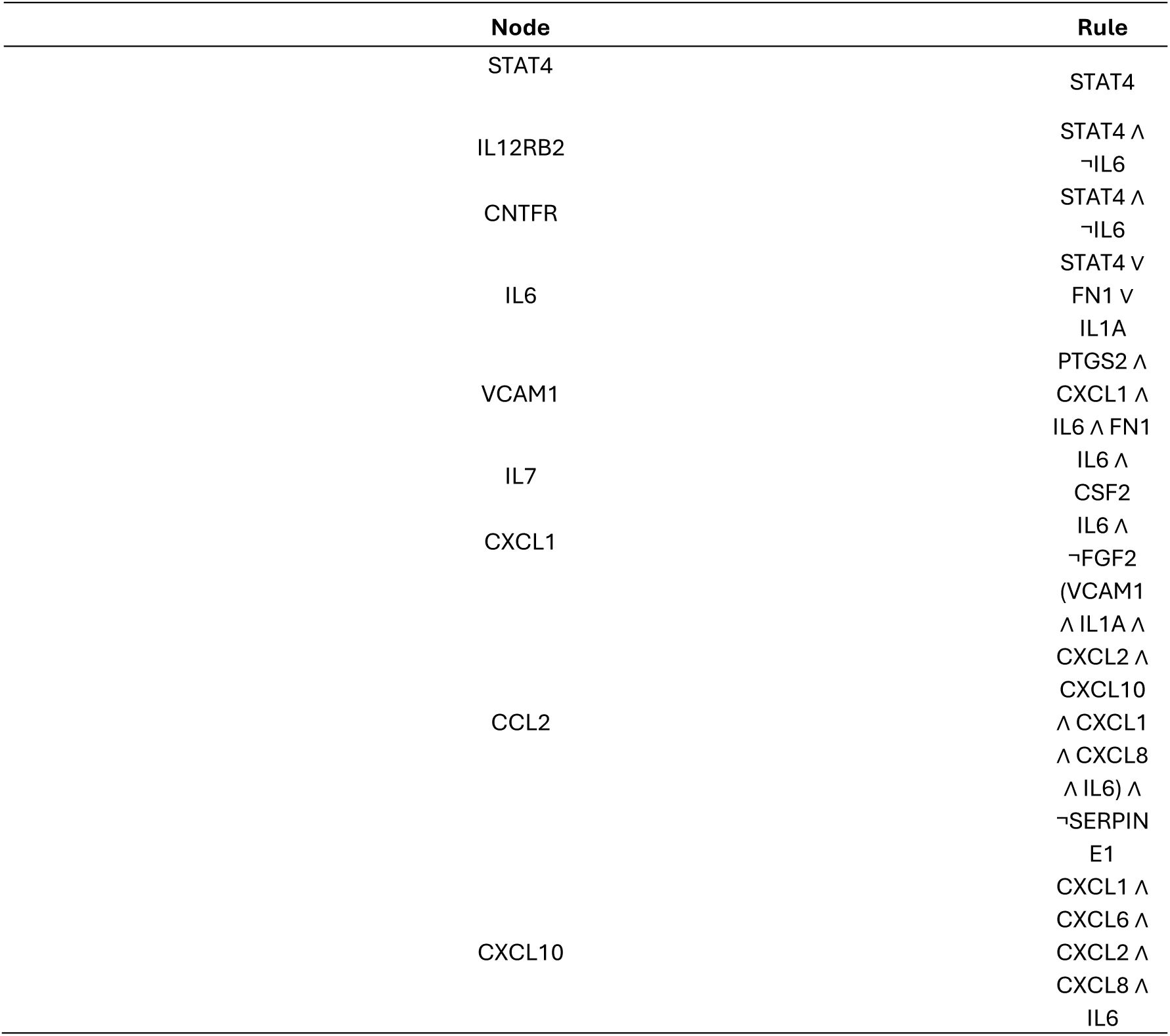
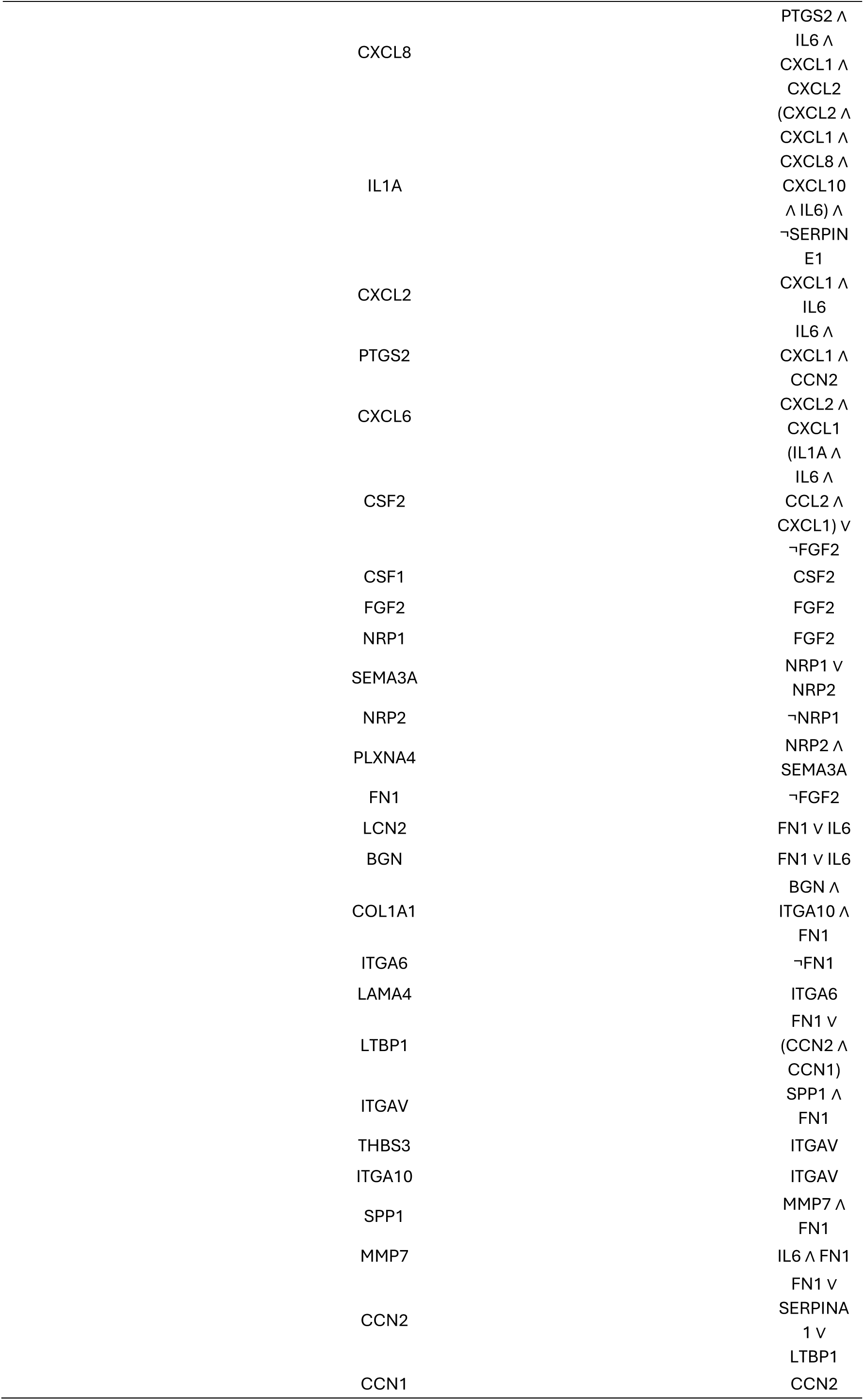
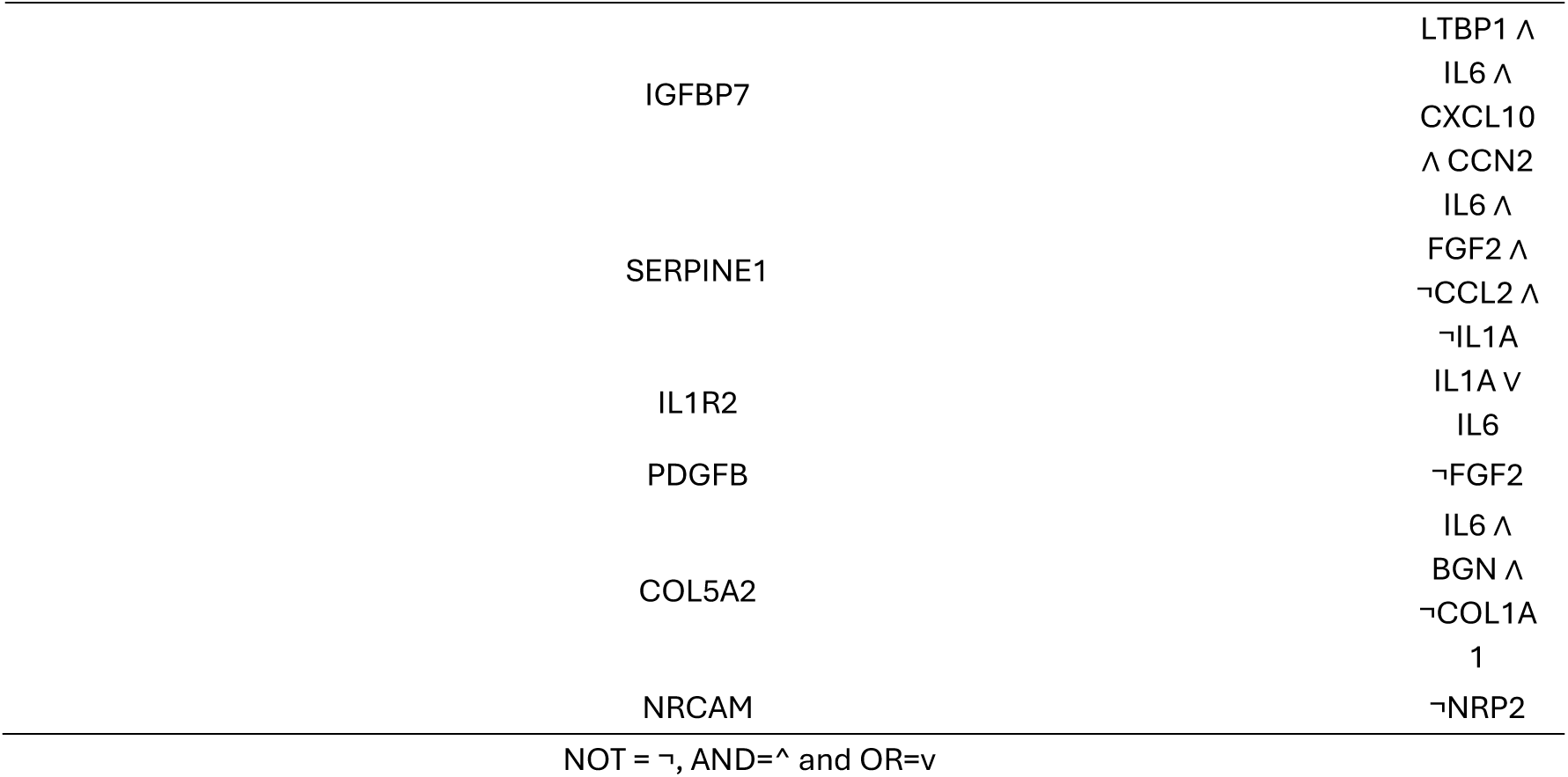
Algebra Boolean Rules for CMDV Non-directed network.

These new interactions included STAT4 with CNTFR [33, 34]; IL6 with IL1R2 [35–37], MMP7 [38], SERPINE1 [39], IGFBP7 [40, 41], COL5A2 [42–44], LCN2 [45, 46] and BGN [47]. We also included the newly described interaction between BGN and COL5A2 [48]; and of CXCL1 with VCAM1 [49] and PTGS2 [50]. Additionally, we created an edge between FN1 and MMP7 [51] and a loop among LTBP1, CCN1 and CCN2 [52]. The latter also interacts with PTGS2 [53]. Additionally, an edge was defined among LTBP1, IGFBP7 [53] and SERPINE1 [54]. SERPINE1 also interacts with IL1 [55] and FGF2 [56]. We also included the interaction of IGFBP7 with CXCL10 [57]. The resulting network consists of 41 nodes and 99 edges; however, the average clustering coeficient decreases to 0.56. The longest path (diameter) in the network between two nodes encompasses 6 edges, while the shortest path (radius), from the central node to a peripheral one, consists of 3 edges (Supplementary table 3B). Community analysis identified six highly interconnected clusters (see Table 2). However, those communities are not well-defined (Mod=0.44) because they are not isolated; rather, there is a uniform interaction among them. This is due to the low connectivity between nodes (density=0.1207), which range from highly connected to poorly connected (assortativity=-0.1176).

**Table 2.**
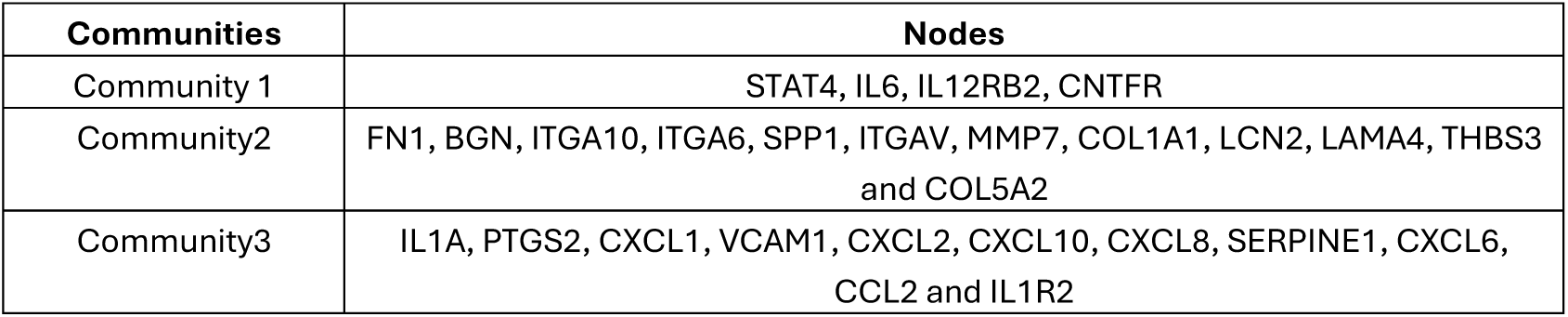
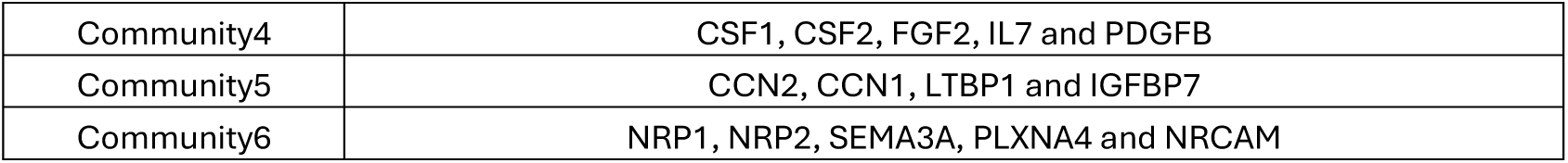
CMDV network communities.

| Communities | Nodes |
| --- | --- |
| Community 1 | STAT4, IL6, IL12RB2, CNTFR |
| Community2 | FN1, BGN, ITGA10, ITGA6, SPP1, ITGAV, MMP7, COL1A1, LCN2, LAMA4, THBS3 and COL5A2 |
| Community3 | IL1A, PTGS2, CXCL1, VCAM1, CXCL2, CXCL10, CXCL8, SERPINE1, CXCL6, CCL2 and IL1R2 |
| Community4 | CSF1, CSF2, FGF2, IL7 and PDGFB |
| Community5 | CCN2, CCN1, LTBP1 and IGFBP7 |
| Community6 | NRP1, NRP2, SEMA3A, PLXNA4 and NRCAM |

On the other hand, centrality analysis (Supplementary Table 3A) confirmed that IL6 is a central node because it interacts with 22 nodes, occurs 312.78 times in the shortest path between nodes (betweenness), and has a maximum eigenvector (1), even though its closeness in the network is low (0.13).

A second central node is FN1, but its interacting nodes (12) are fewer than IL6. Betweenness was 290.81 but its eigenvector was low (0.33). However, closeness was similar to that of IL6. Other relevant nodes include CXCL1, IL1α, FGF2, and CCL2. However, centralities vary. For example, FGF2 interacts with 8 nodes, and its betweenness is 224.10, but its eigenvector (0.33) is lower than that of IL1α, which interacts with 11 nodes but has a betweenness of 13.8 and an eigenvector of 0.8. This indicates that IL1α is more relevant in the present framework.

In order to enable simulations in the model, we implemented thresholds and modulators using the SPIDDOR R Studio package. These regulators can be used to modify the network generally or locally [58]. Before imposing arbitrary values, we needed to determine how many iterations would be required to reach a minimum threshold of FN1 degradation. To do this, we used a first-order modified model with regular catabolic degradation rate [59].

FN1 required 23 iterations to reach the minimum threshold. The maximum thresholds and modulators are 24, establishing 23 FN1-dependent interactions. This was regularly used to induce a state change in a particular node. The only exception was IL1R2, which displayed a threshold of 22, allowing the node to maintain its presence in the model.

Other regulators influence the network in initial stages and are defined by how much time a cell takes to perform transcription, translation, or metabolic synthesis. The first two processes can be assumed to be relatively quick and comparable to each other [60]. Also, CMDV contains some proteins corresponding to specific nodes, for example IL6 [15], which increases the cellular response rate. However, metabolic synthesis involves multiple steps of diverse duration. An example is VCAM1, which is produced by endothelial cells in response to inflammation [61, 62]. This introduces an asynchronous behavior component in the system, which better simulates the experimental results.

### 2.5. 2.5. NDAM-CMDV robustness, accuracy and attractors

The accuracy and robustness of the model in describing the experimental data were then evaluated, taking the above-described conditions and parameters into consideration. 25000 simulations, each consisting of 100 iterations with asynchronous activation, were performed **(Figure 6A and Supplementary Table 4)**. The system exhibited the expected behavior: the pro-inflammatory and endothelial dysfunction nodes were activated during the first 25 iterations and then turned of. Some nodes (CSF1, CSF2, ITGA6, LAMA4, and NRCAM) displayed a mildly nonspecific (noisy) signal during the first three iterations. However, these did not exceed an activation state of 0.5, thereby did not impair the proper evolution of the system. The model is built to correct such disturbances by stabilizing the state of these nodes ([dx]((t))/dt=0) until they become activated. In this manner, such behavior preserves the dynamics of the network.

The activation percentage, media, and standard deviation from each node is highly predictable because they exhibit high activation but low variance (STAT4, FGF2, IL6, NRP1, SEMA3A, BGN, and LCN2). This means these nodes are consistent across the entire system. Another subgroup of nodes displays low means and variance (IL1A, CCL2, CSF2, CSF1, and IL7). Finally, another high predictability group includes those with high means and medium variance (CCN1, CCN2, LTBP1, and IL1R2), indicating that they are activated and tend to be consistent.

However, other nodes tend to be more unpredictable due to their high variance and average means (SERPINE1, COL5A2, NRCAM, LAMA4, and ITGA6), or high variance and low means (IL12RB2, NRP2, CNTFR, PDGFB, FN1, and PLXNA4).

On the other hand, we also determined the system attractors (Supplementary Table 5). The analysis identified them using various approaches, including synchronic and asynchronous methods, thereby identifying synchronic and asynchronous attractors. In the first two analyses, attractors were consistent (STAT4, IL6, FGF2, NRP1, SEMA3A, LCN2, BGN, ITGA6, LAMA4, LTBP1, CCN1/2, SERPINE1, IL1R2, COL5A2, and NRCAM), suggesting that these nodes have influenced the dynamic behavior of the system. However, this analysis identified 31 nodes with a high recurrence. This strongly indicates that the model may be misaligned. Nevertheless, eight attractors (IL6, SEMA3A, LCN2, BGN, LTBP1, CCN1, CCN2, and IL1R2) were common in all the analyses. These indicate that the network has a moderate degree of complexity, high stability, and predictability, which are strong indicators of robustness. Additionally, the accuracy of the CMDV model against the experimental data is good, indicating its potential to study endothelial dysfunction in dengue using a cellular approach.

## Legends to the Supplementary Figures

**Supplementary Figure 1.**
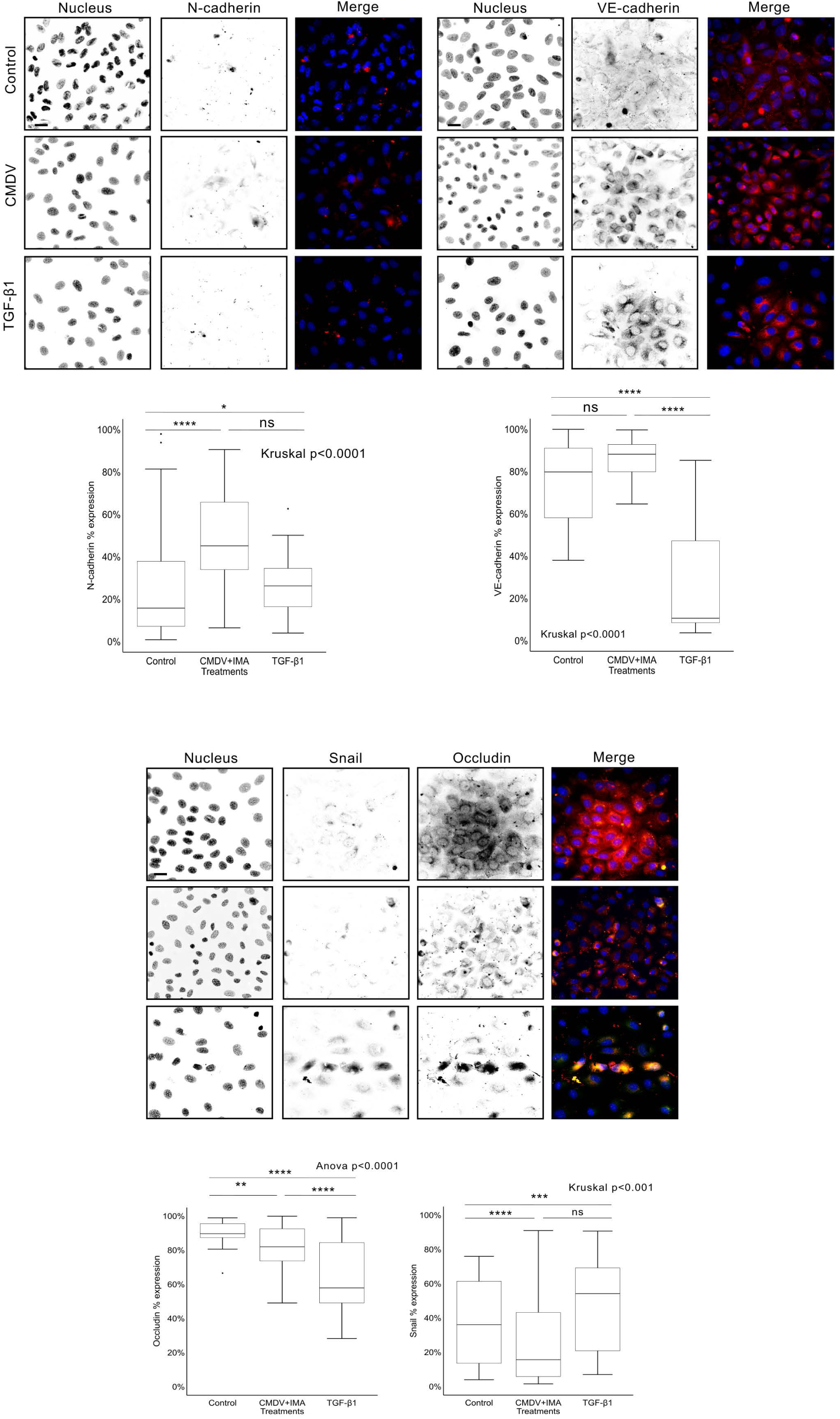
Efect of exposure to imatinib for 48h in the expression of expression of selected endothelial and mesenchymal markers in endothelial cells in response to CMDV. Cells were treated with imatinib (10 µM) for 48h after 120h exposure to CMDV. Representative images of HMEC-1 cells treated as indicated and stained for DNA (DAPI) and N-Cadherin (A), VE-Cadherin (B) and occludin and Snail (C). In overlays, red is N-cadherin, VE-cadherin or occludin as indicated, and green is Snail; blue represents DAPI staining. TGF-β1 was used as a positive control. Scale bar=20 µm. Under the microphotographs, quantitative analysis is shown. Data are presented as mean ± SEM from >100 fields examined in three independent experiments. See Material and Methods for additional details. Statistical tests are indicated, and significance is as follows: *p < 0.05, **p < 0.01, ***p < 0.001, ****p < 0.0001.

**Supplementary Figure 2.**
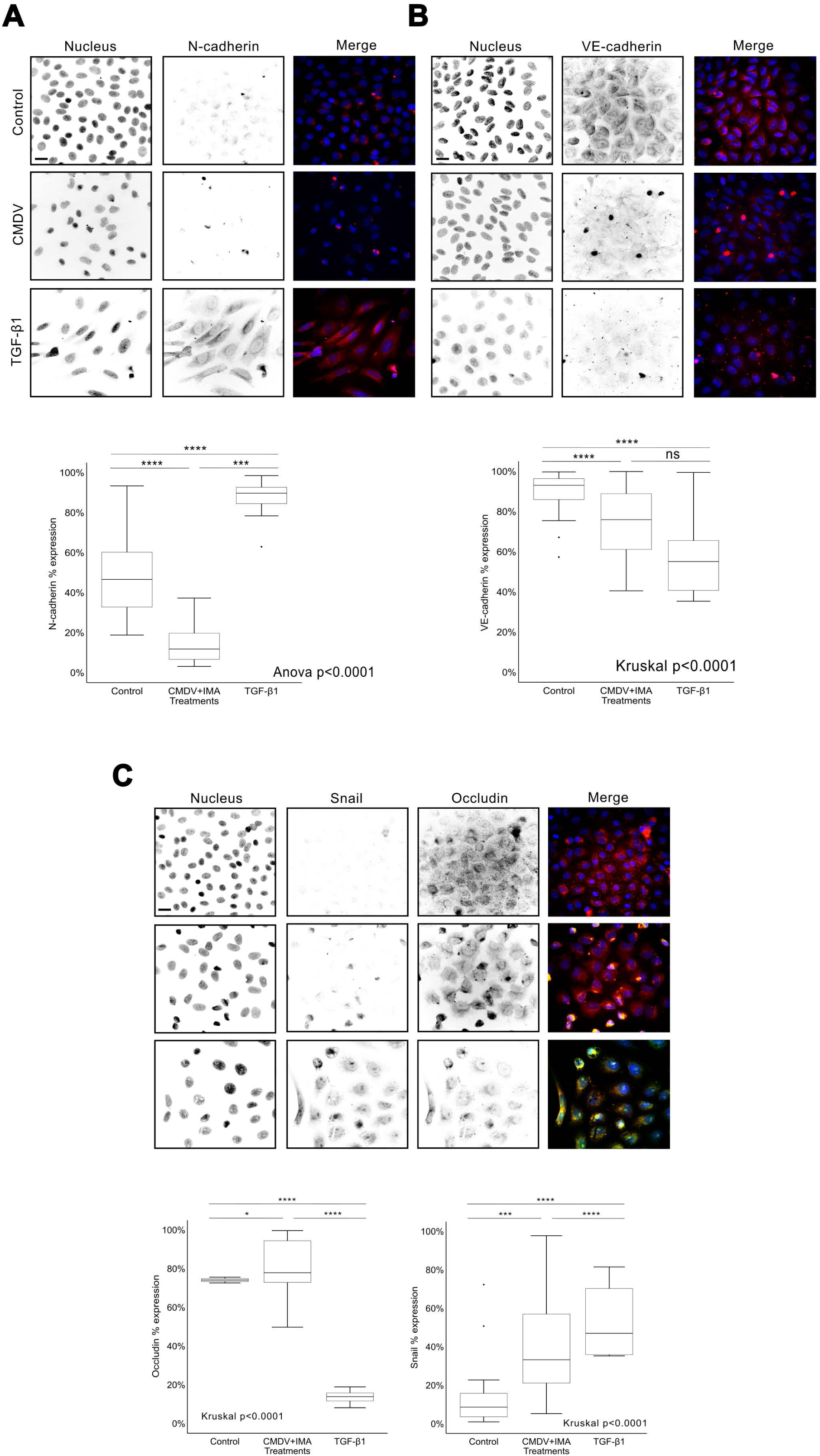
Efect of exposure to imatinib for 120h in the expression of selected endothelial and mesenchymal markers in endothelial cells in response to CMDV. Cells were treated with imatinib (10 µM) for 120h after 120h exposure to CMDV. Representative images of HMEC-1 cells treated as indicated and stained for DNA (DAPI) and N-Cadherin (A), VE-Cadherin (B) and occludin and Snail (C). In overlays, red is N-cadherin, VE-cadherin or occludin as indicated, and green is Snail; blue represents DAPI staining. TGF-β1 was used as a positive control. Scale bar=20 µm. Under the microphotographs, quantitative analysis of images as in top. See Material and Methods for details. Data are presented as mean ± SEM from >100 fields examined in three independent experiments. Statistical tests are indicated, and significance is as follows: *p < 0.05, **p < 0.01, ***p < 0.001, ****p < 0.0001.

**Supplementary Figure 3:**
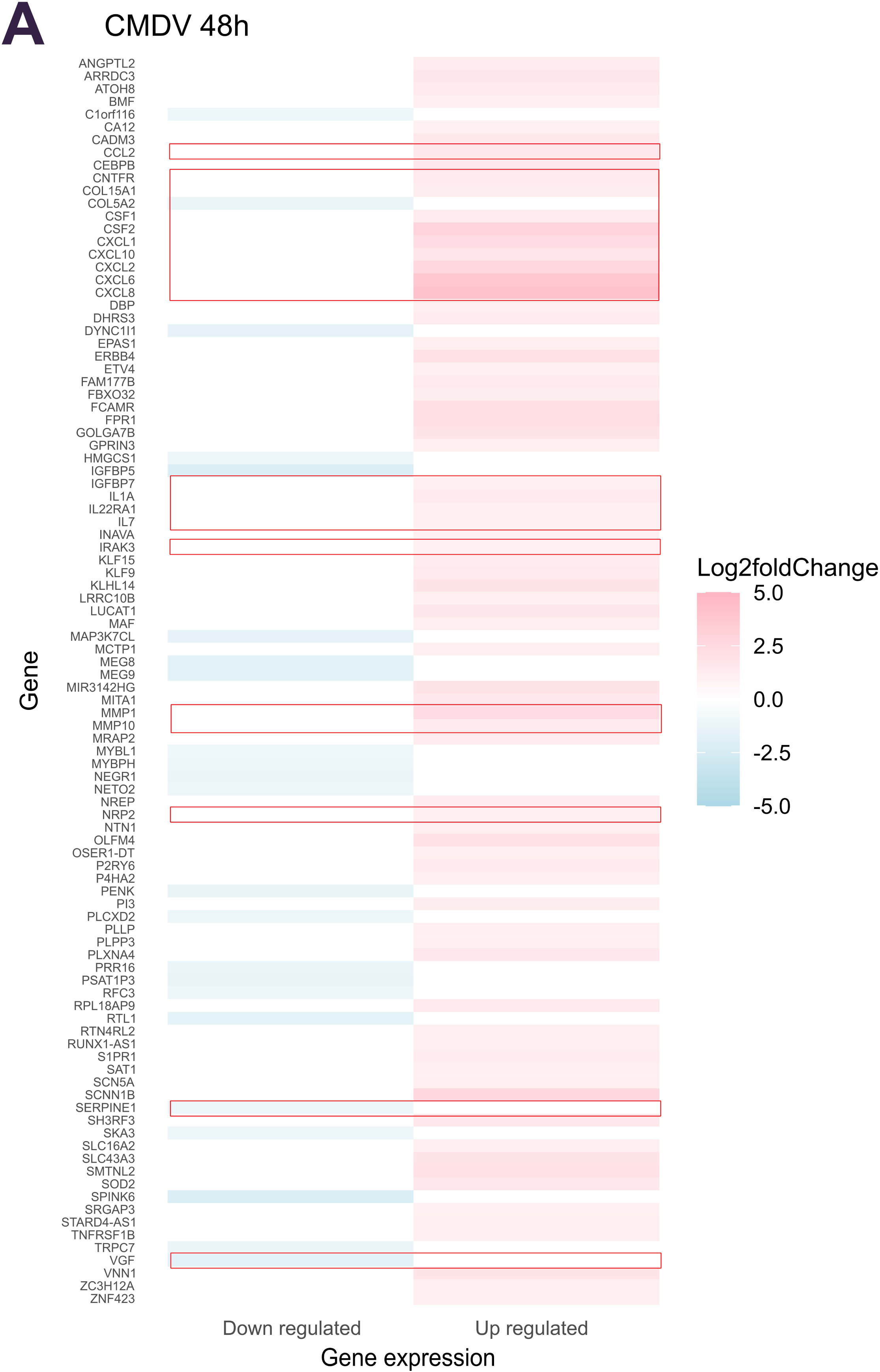
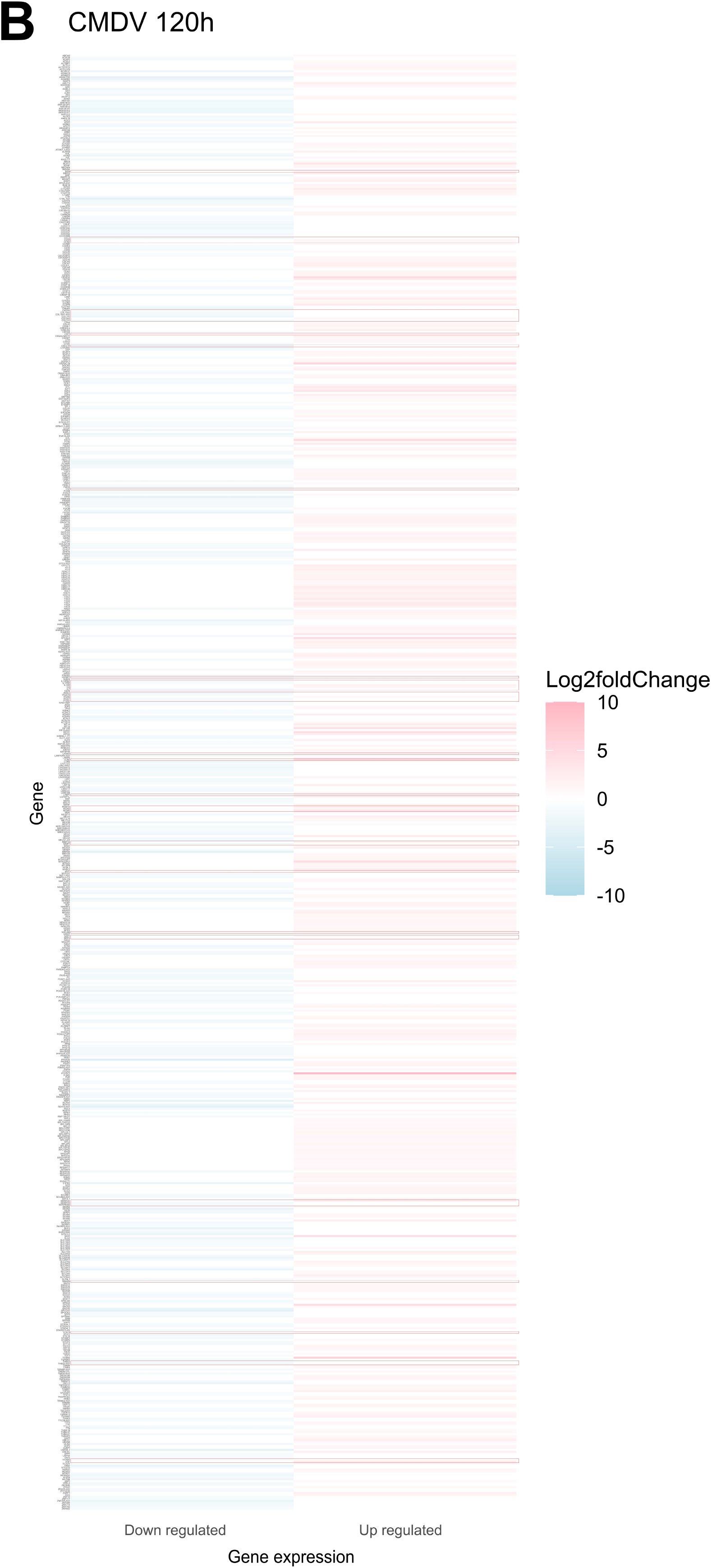
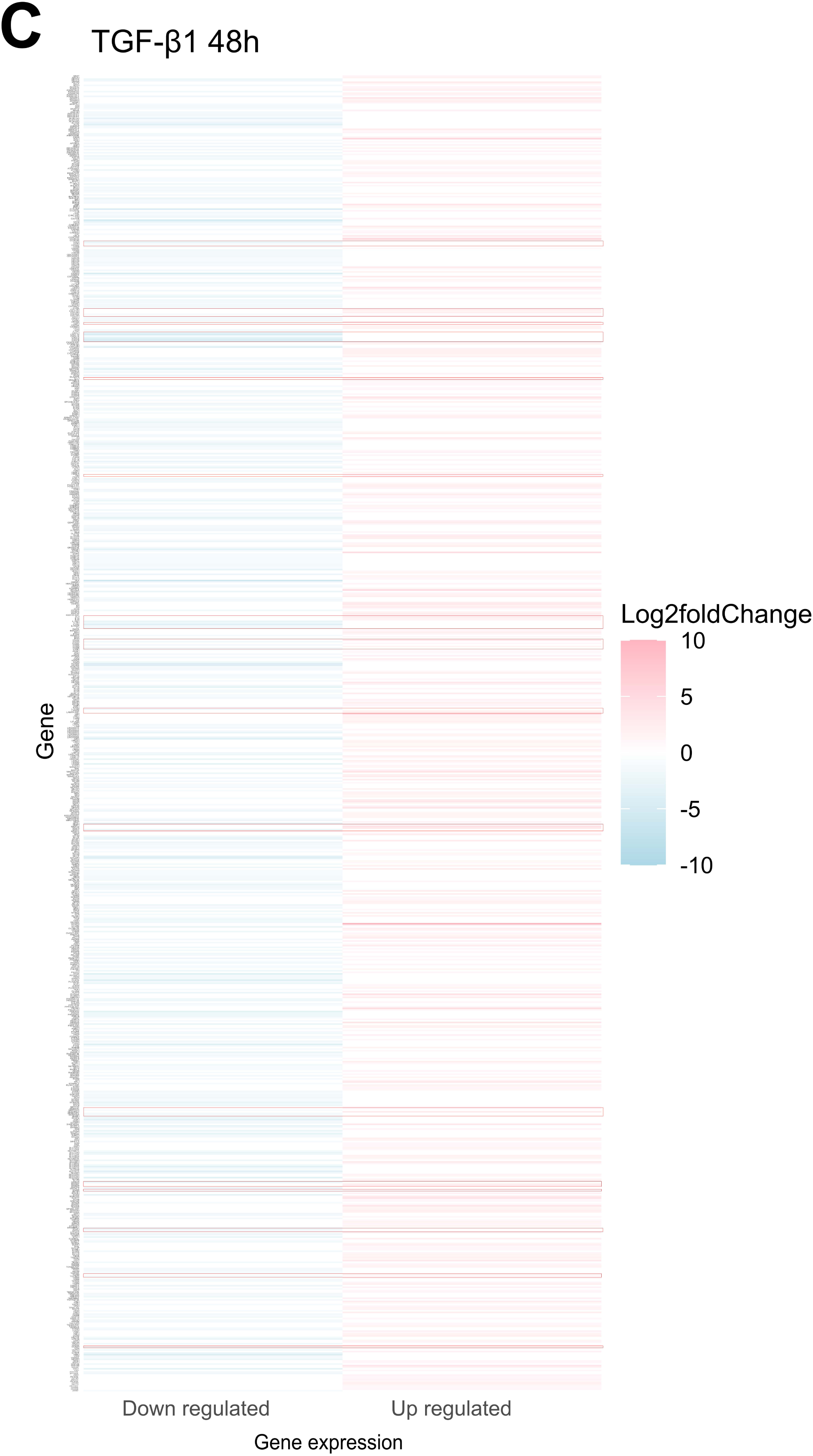
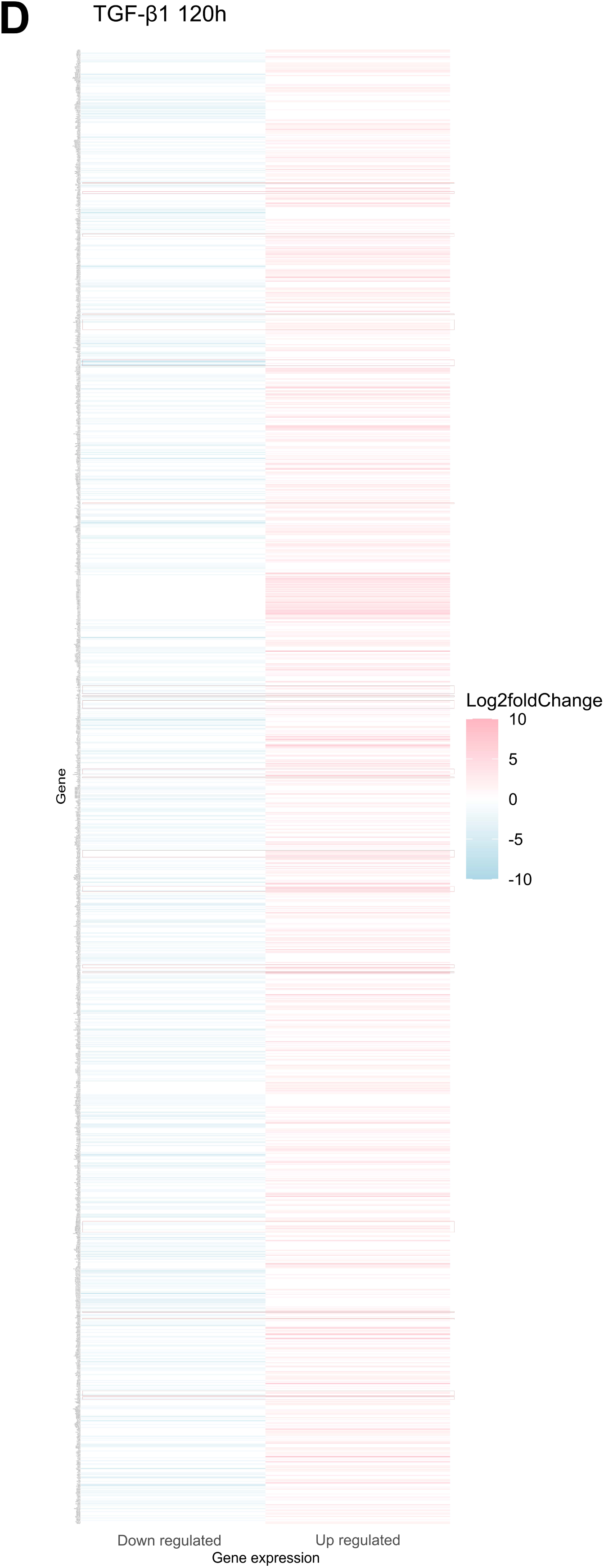
Heat maps of DEGs in HMEC-1 cells treated with CMDV or TGF-β1. Figures represent cells treated with CMDV for 48h (A), CMDV for 120h (B), or TGF-β1 for 48h (C) and TGF-β1 for 120h (D). Heatmaps corresponding to the CMDV treatments illustrate the numeric diferences between the two timepoints.

**Supplementary Figure 4.**
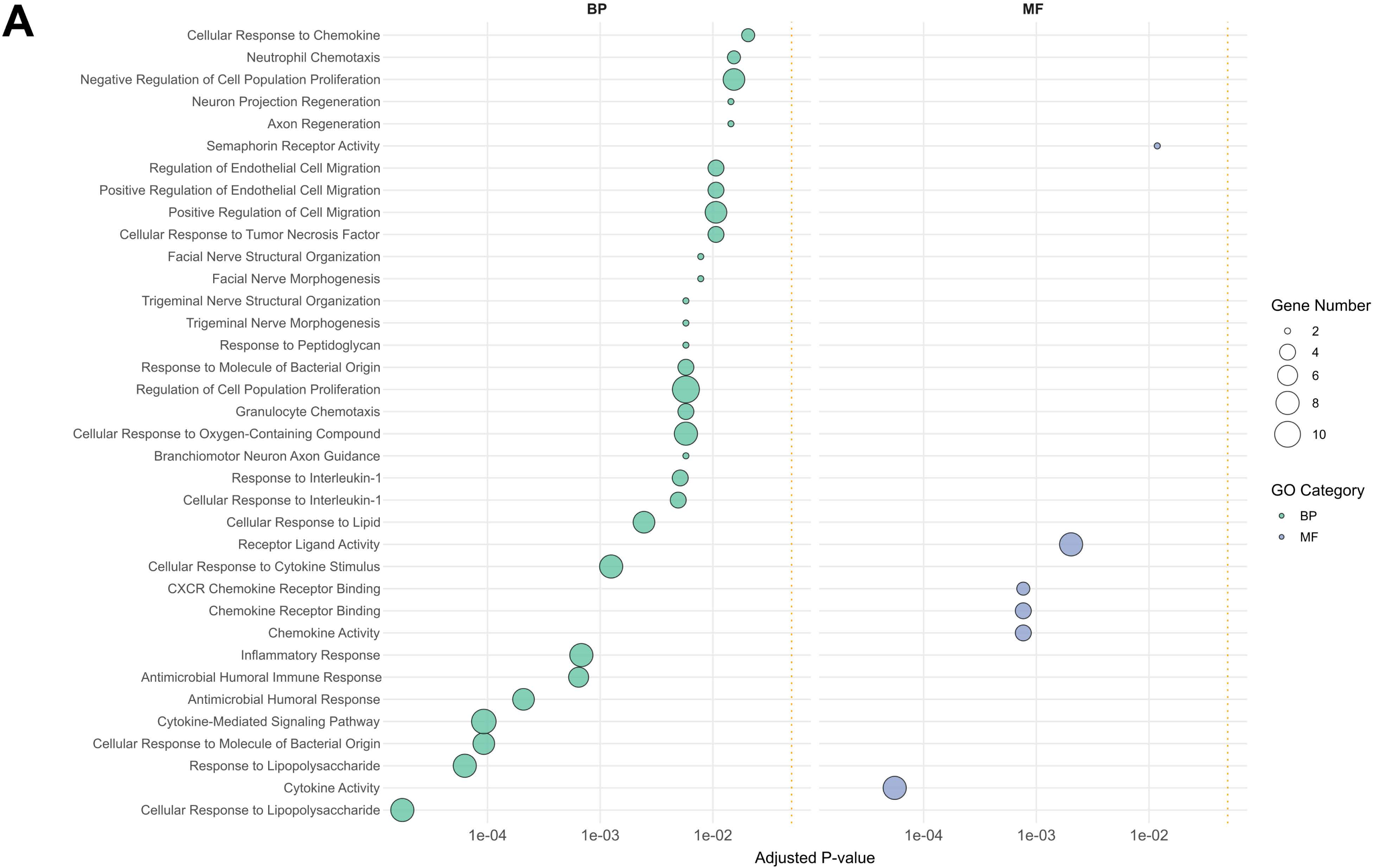
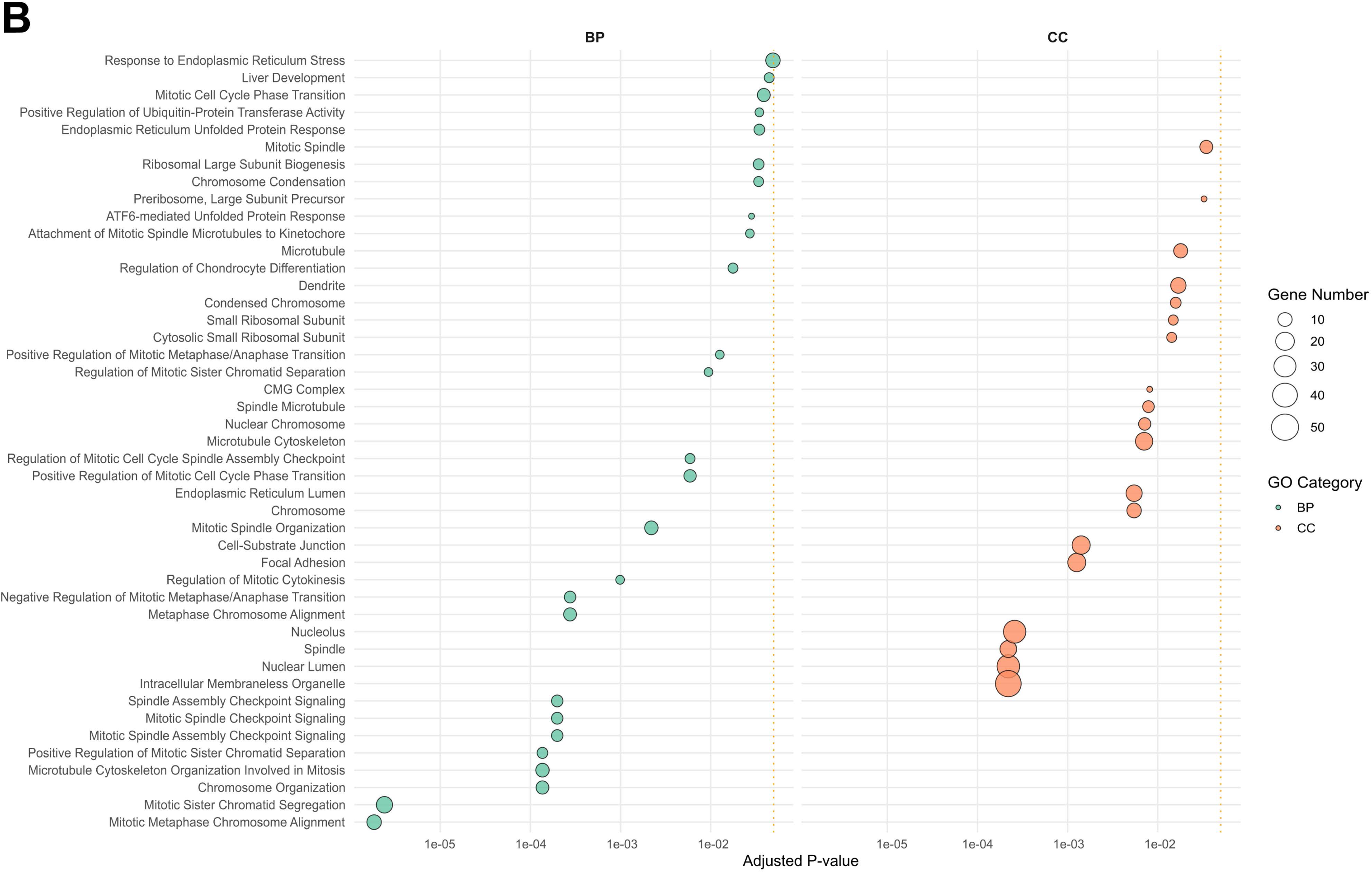
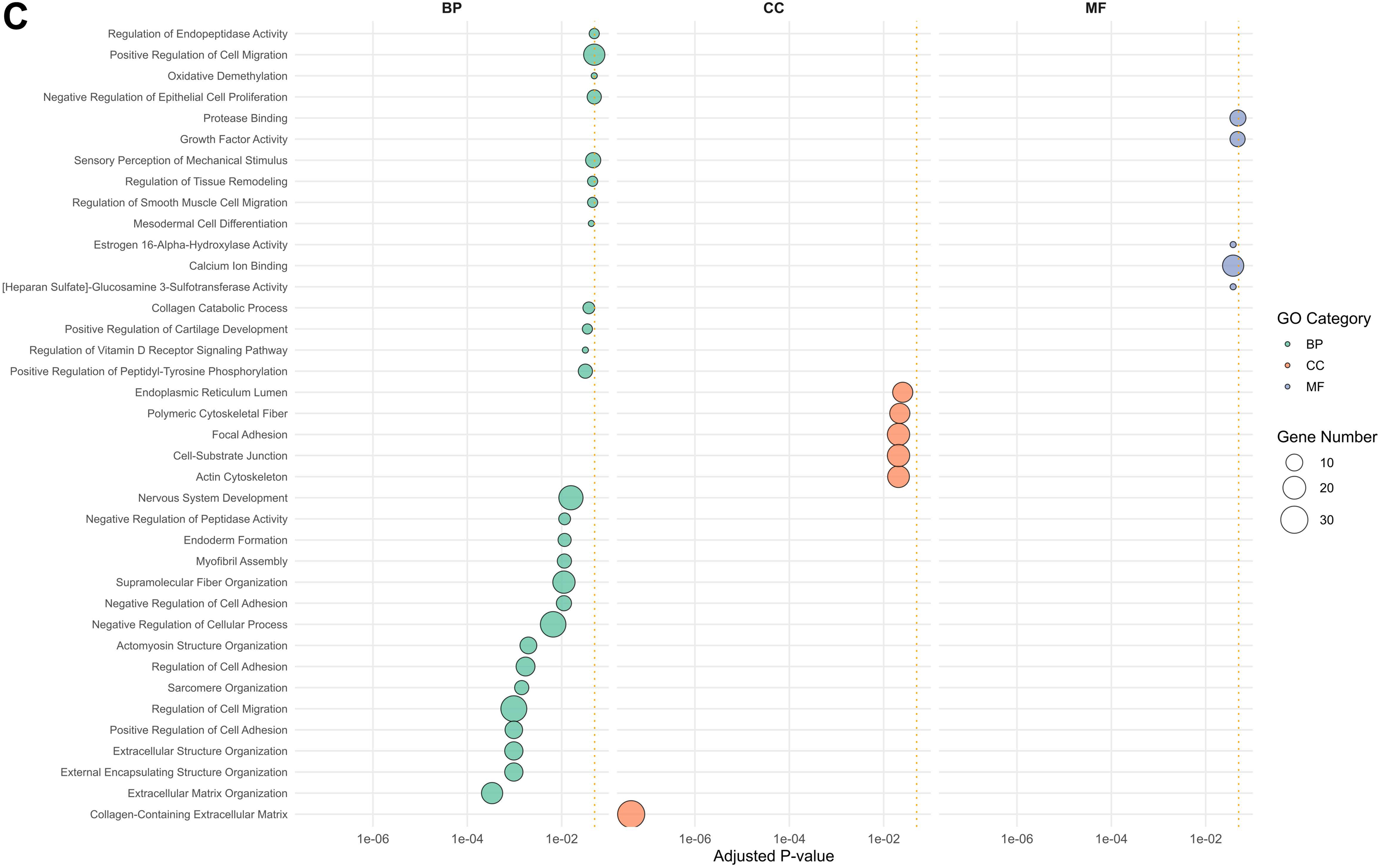
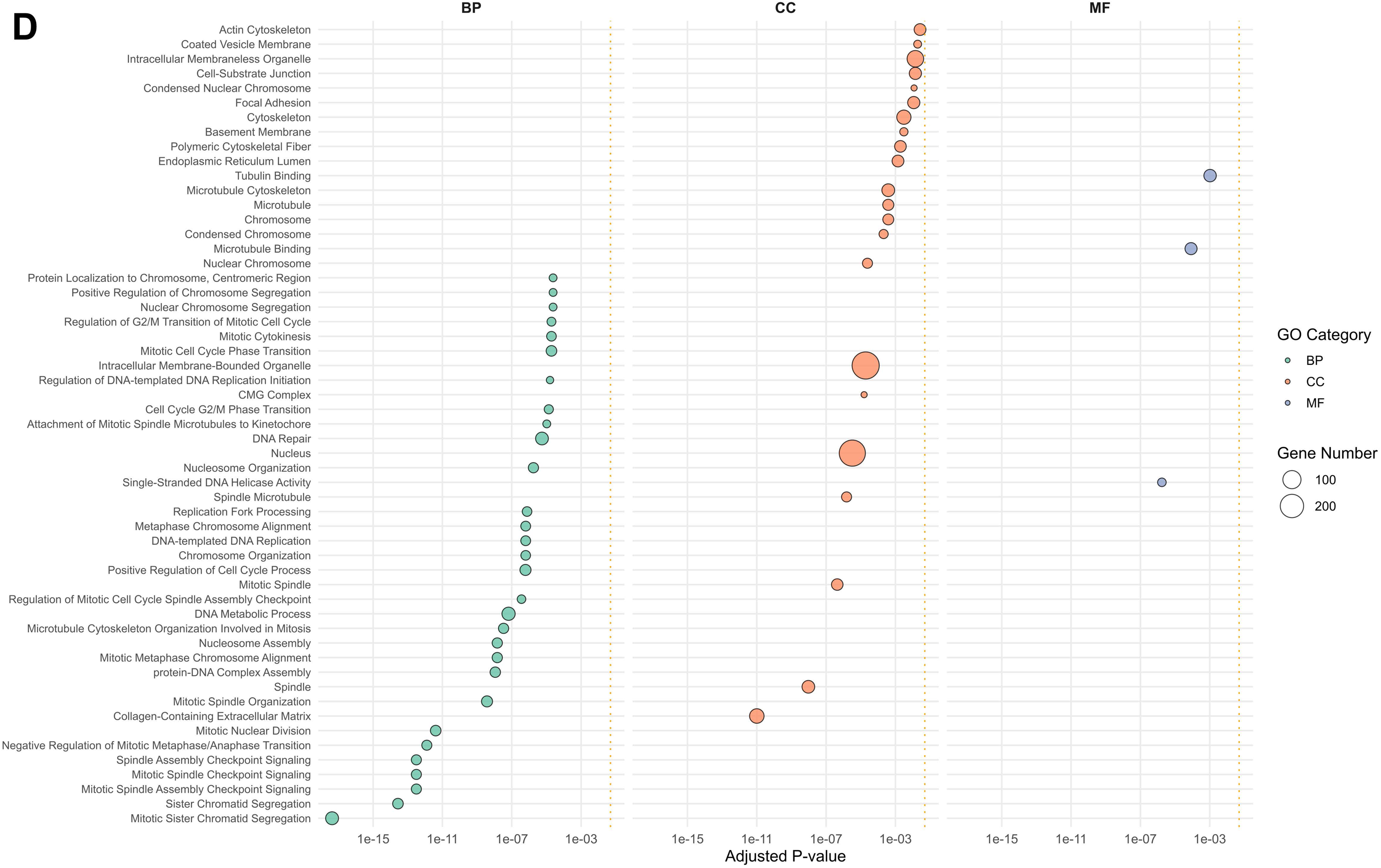
Gene enrichment analysis of HMEC-1 cells treated with CMDV or TGF– β1. Only data points with p-values < 0.05 are shown. (A-B) Biological Processes (BP) and Molecular Functions (MF) altered by CMDV after 48 h. (B) BP and Cellular Components (CC) altered by CMDV after 120 h. (C-D) BP, CC and MF altered by TGF-b1 after 48 (C) and 120 (D) hours.

**Supplementary Figure 5.**
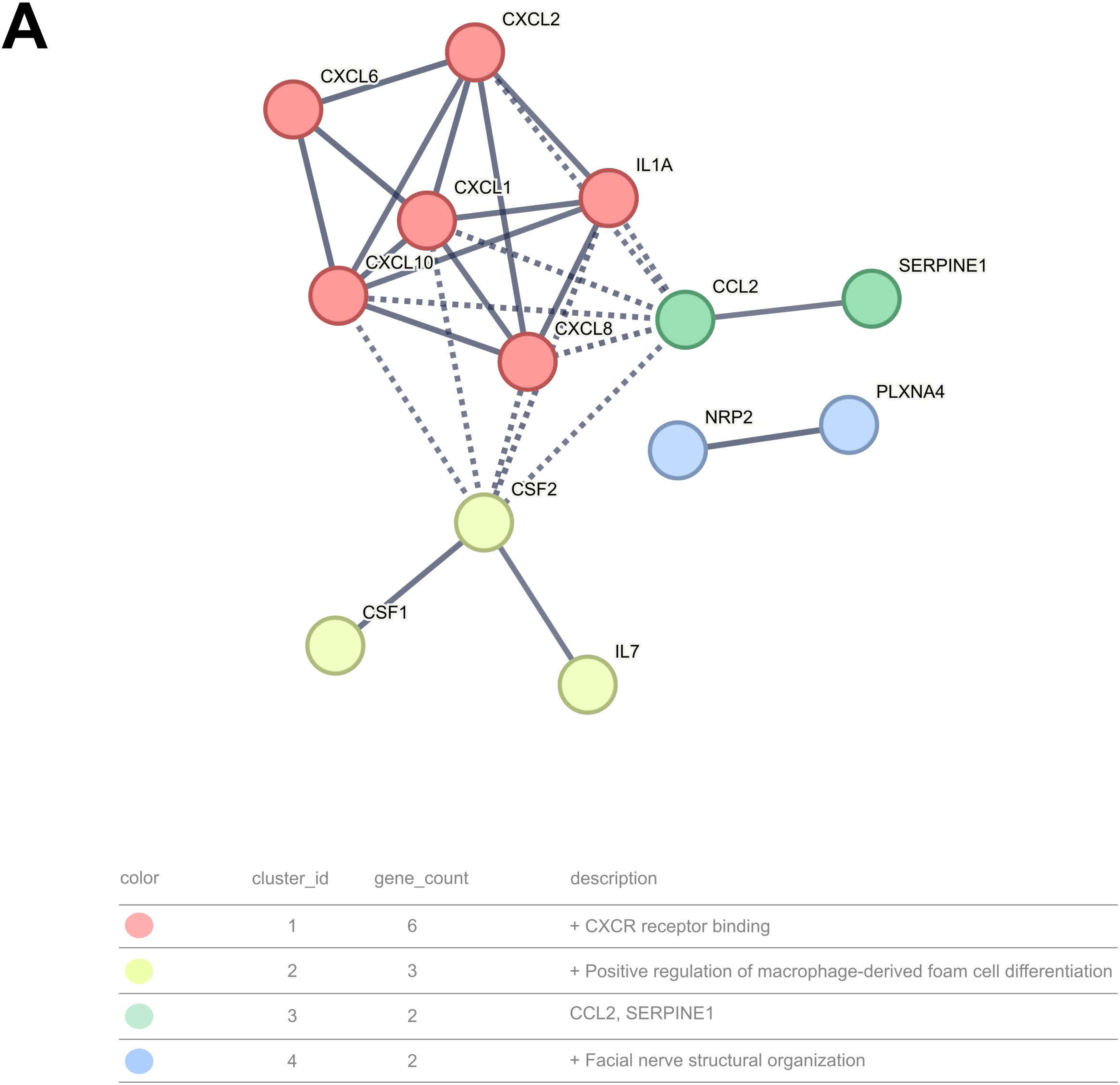
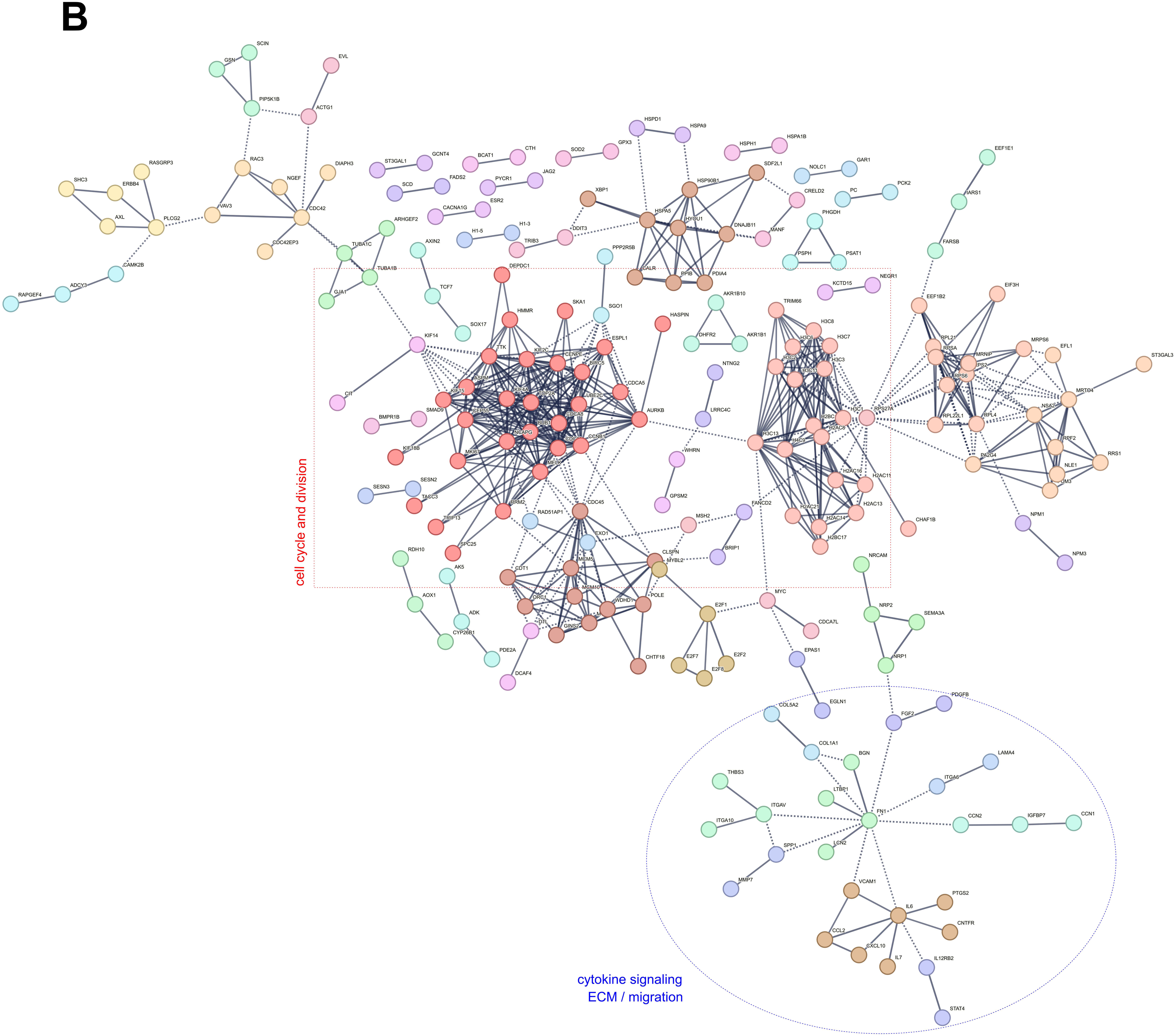
Interactomes of HMEC-1 cells treated with CMDV for 48h and 120h. CMDV interactome at 48h (A) and 120h (B). In (B), dashed red box denotes clusters involved in cell cycle and division (mostly clusters 1 and 2), whereas blue oval highlights cytokine signaling (cluster 7) and neighbors involved in ECM remodeling and cell migration (clusters 13, 15, 29

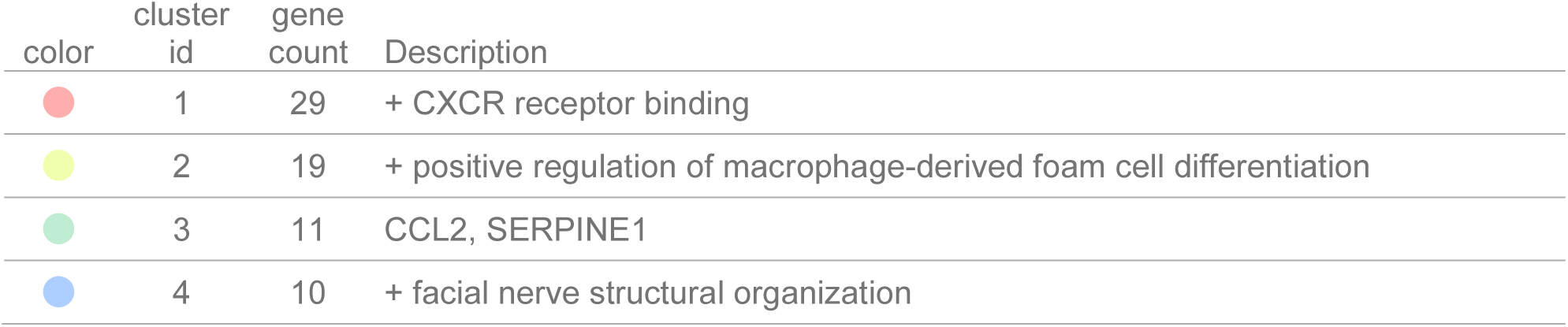
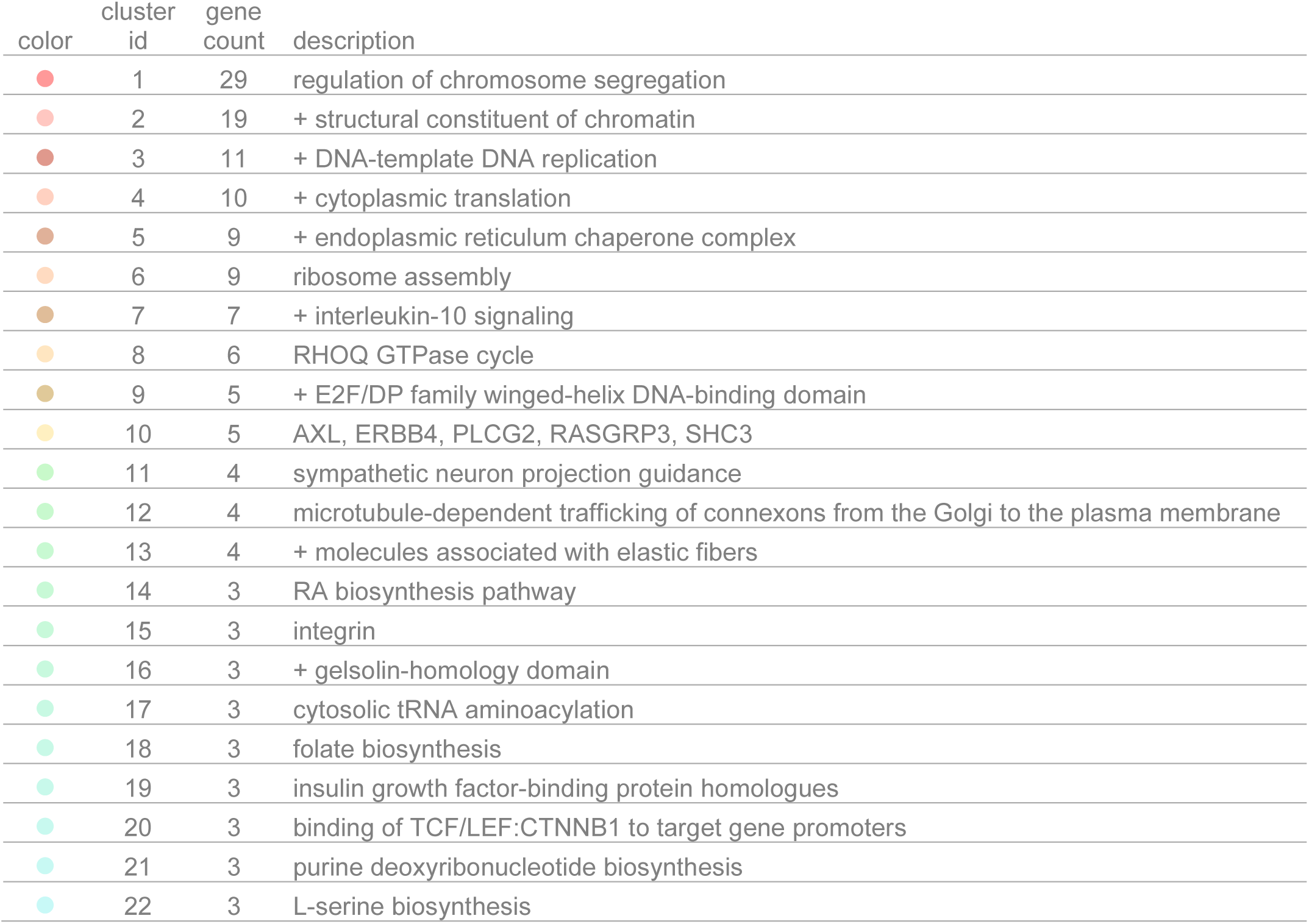
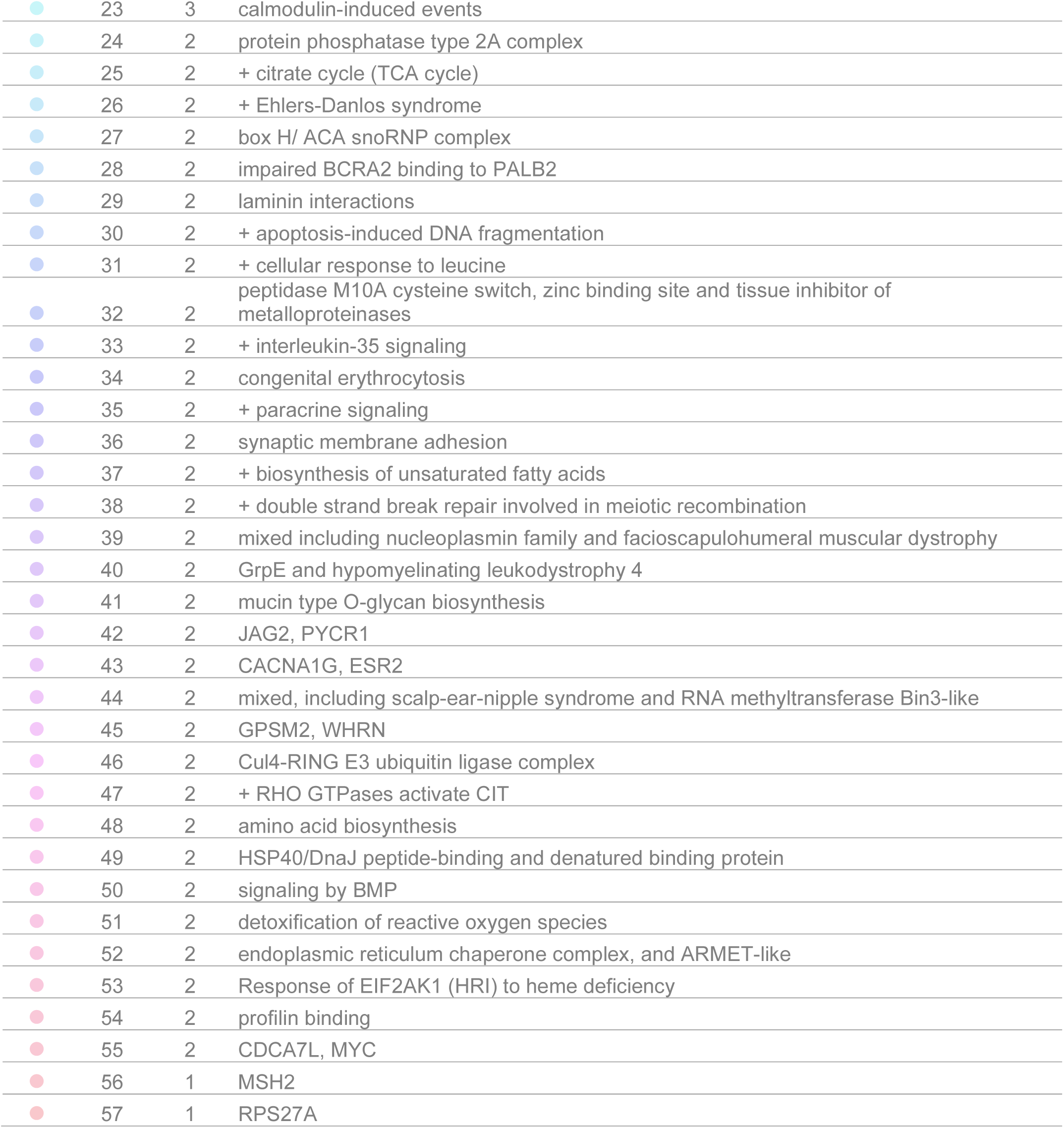

**Supplementary Figure 6.**
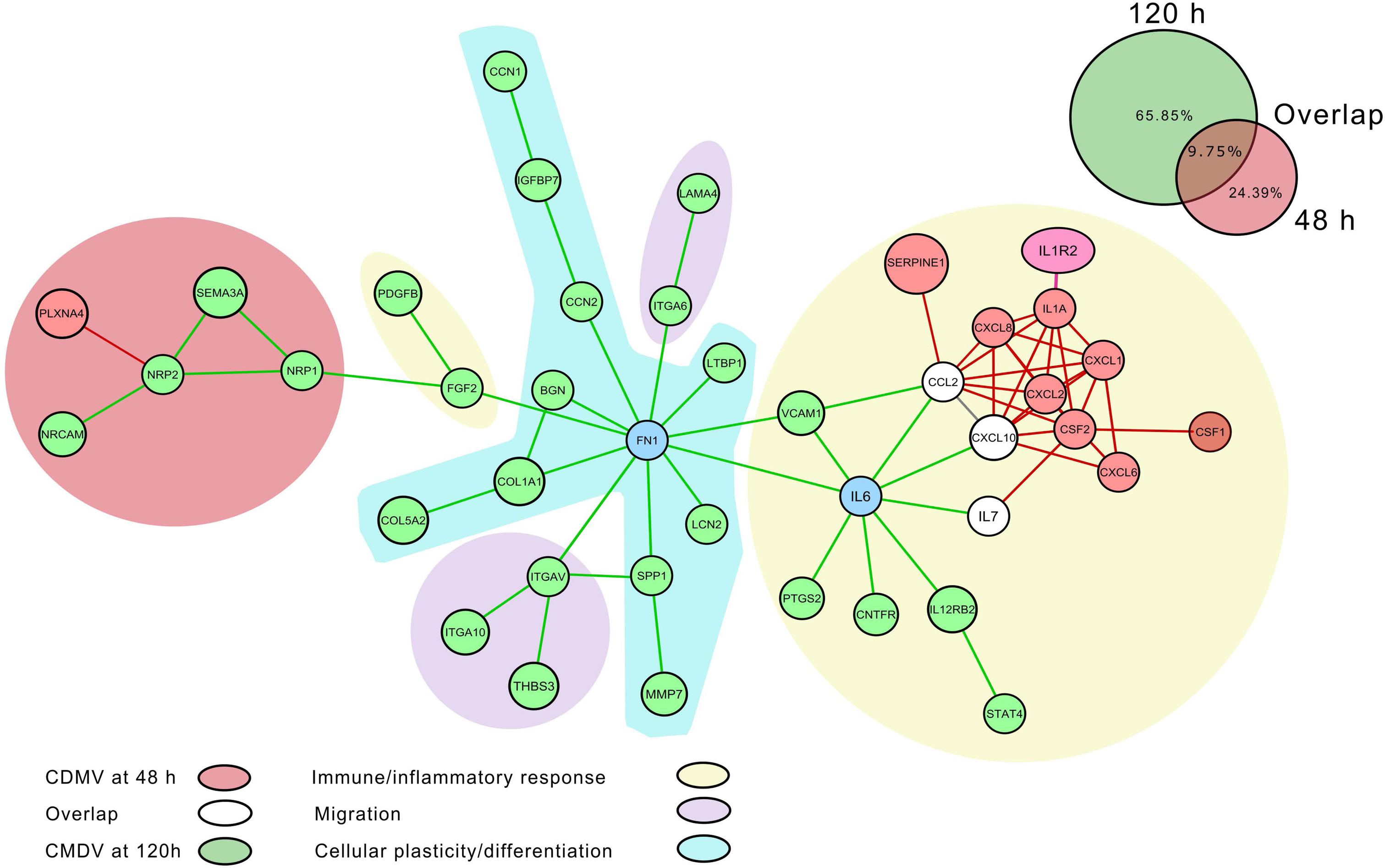
Merged network using DEGs from CDMV at 48h (A) and 120h (B). The cellular response between the two times is not isolated and are biologically involved. The networks were grouped using the MCL clustering algorithm, representing functional groups and were classified in immune/inflammatory response (yellow clusters), migration (purple clusters) and cellular plasticity/diferentiation (blue cluster). The red nodes and edges are from the cellular response at 48 h to CMDV, the green ones are from 120h, and the whites/grey are the overlap nodes and edges between the two times. Blue denotes the central nodes in the network.

**Supplementary Figure 7:**
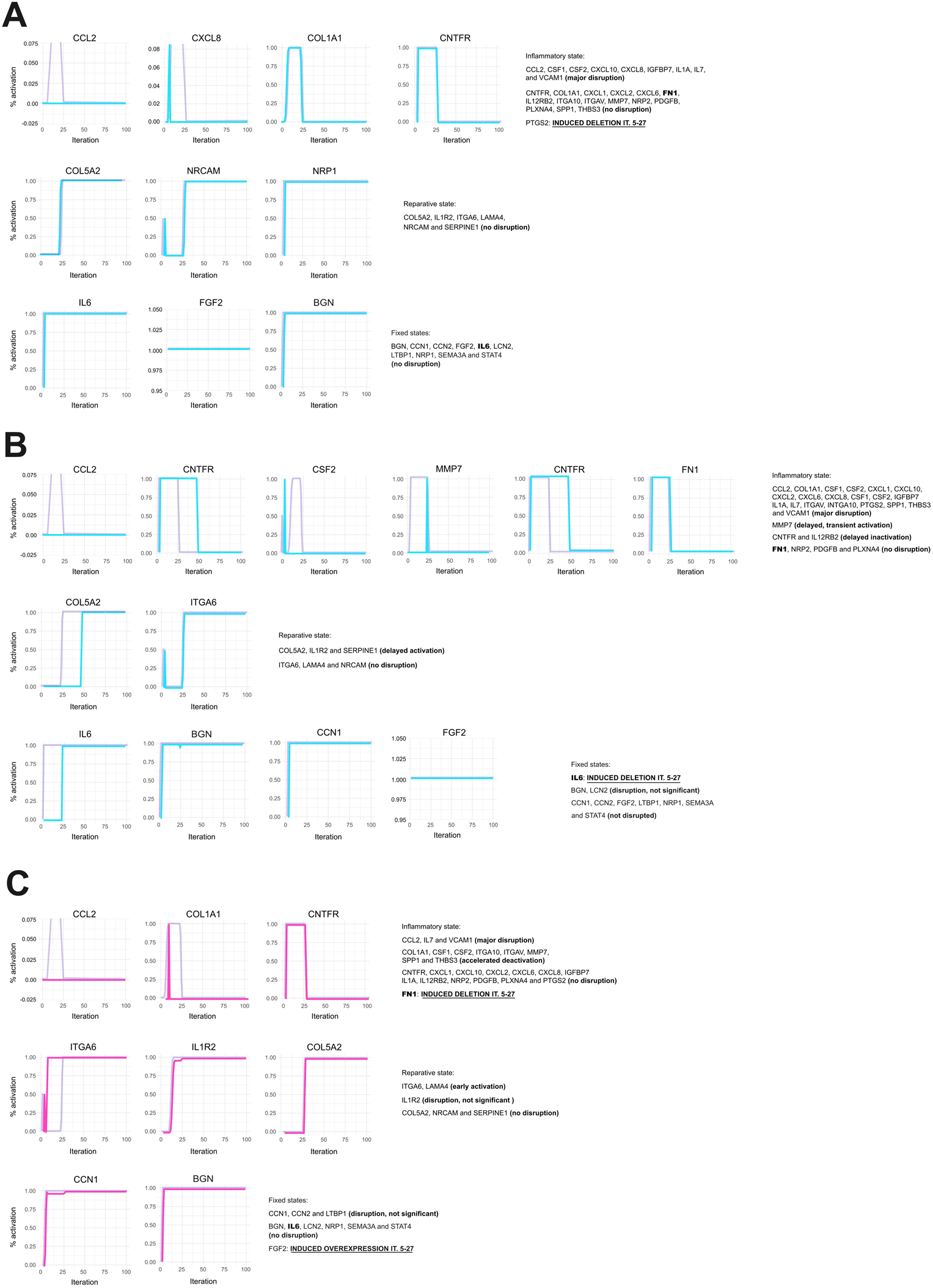
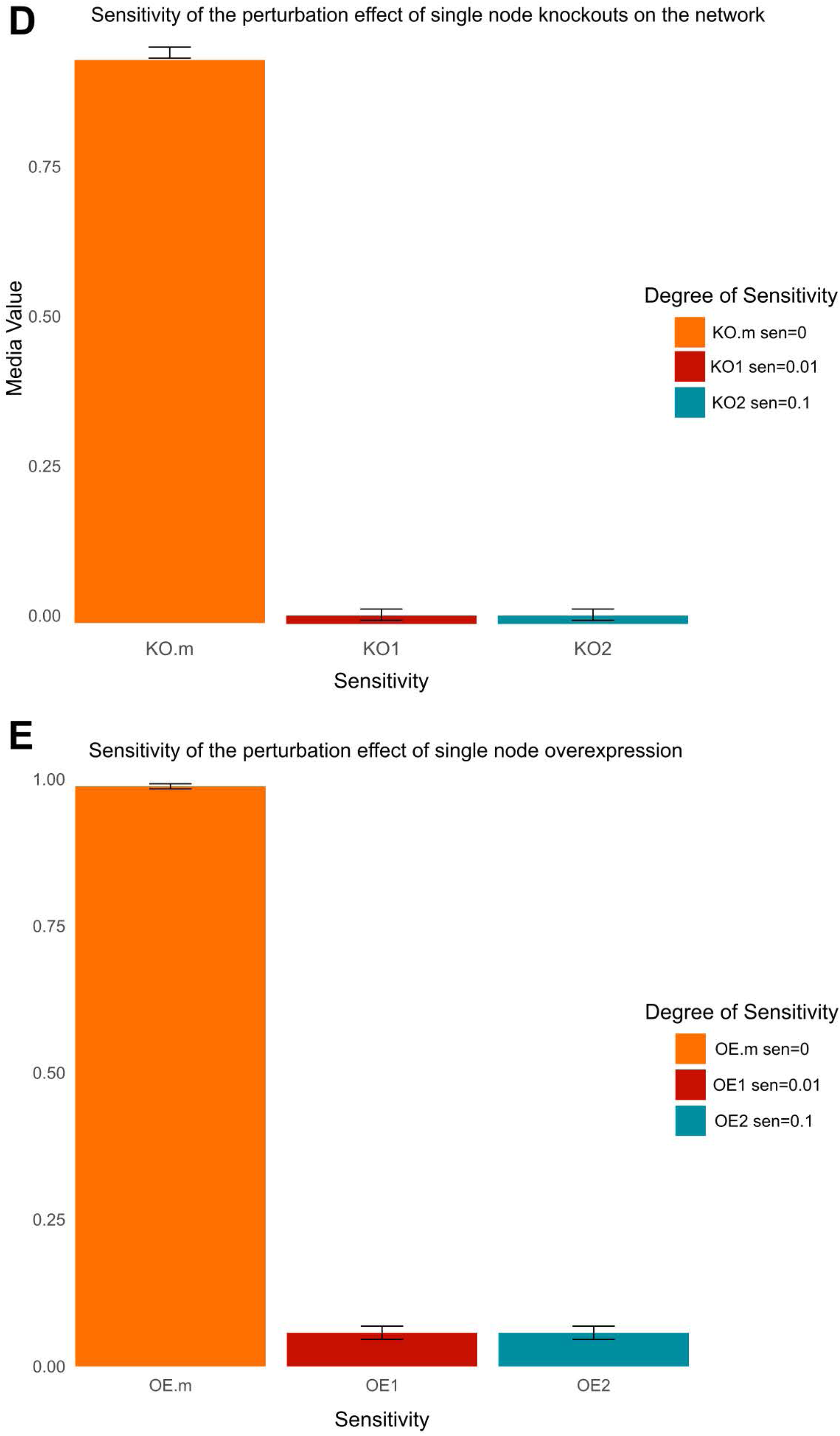
Simulations of perturbation scenarios in response to CMDV in the non-directed network. Modulation of gene modulation in response to (A) PTGS2 knockout; (B) IL6 knockout; (C) FGF2 overexpression+FN1 knockout. For the analysis of single-gene perturbations (A-C), 25000 repetitions were performed. Pale magenta lines represent the state of each node in untreated conditions (as in Figure 6A). (D) Sensitivity over Perturbation in single node knockouts. (E) Sensitivity over Perturbation in single node overexpression.

**Supplementary Table 1.** Potential Pharmacological Targets detected in CMDV DEGs.

| gene | drug | regulatory approval | indication | interaction score | time |
| --- | --- | --- | --- | --- | --- |
| IL7 | ALUM, POTASSIUM | Approved |  | 4.38 | 48h |
| GPR27 | ELOTUZUMAB | Approved | antineoplastic agent | 26.25 | 120h |
| PSPH | METHIONINE | Approved |  | 21.00 | 120h |
| NRP1 | PEGAPTANIB | Approved | for treatment of age-related macular degeneration | 17.50 | 120h |
| SLC1A5 | GLUTAMINE | Approved |  | 13.13 | 120h |
| PPIB | PROLINE | Approved |  | 13.13 | 120h |
| ST6GAL1 | FLOXACILLIN | Approved |  | 11.67 | 120h |
| HYOU1 | DABIGATRAN | Approved | anticoagulant | 11.67 | 120h |
| HSPD1 | CETRORELIX | Approved | antineoplastic agent | 11.67 | 120h |
| GALNT18 | TOCILIZUMAB | Approved | DMARD,antiinflammatory agent | 10.50 | 120h |
| ATP8B1 | ODEVIXIBAT | Approved |  | 8.75 | 120h |
| H1-3 | MITAPIVAT | Approved |  | 8.75 | 120h |
| TRIB3 | INDAPAMIDE | Approved | Antihypertensive Agents; Diuretics | 7.00 | 120h |
| NPR3 | NESIRITIDE | Approved | cardiovascular agent,for treatment of congestive heart failure | 6.56 | 120h |
| IL1R2 | ANAKINRA | Approved | DMARD,antiinflammatory agent | 5.83 | 120h |
| TRIB3 | PERINDOPRIL | Approved | Antihypertensive Agents | 5.83 | 120h |

**Supplementary Table 2.**
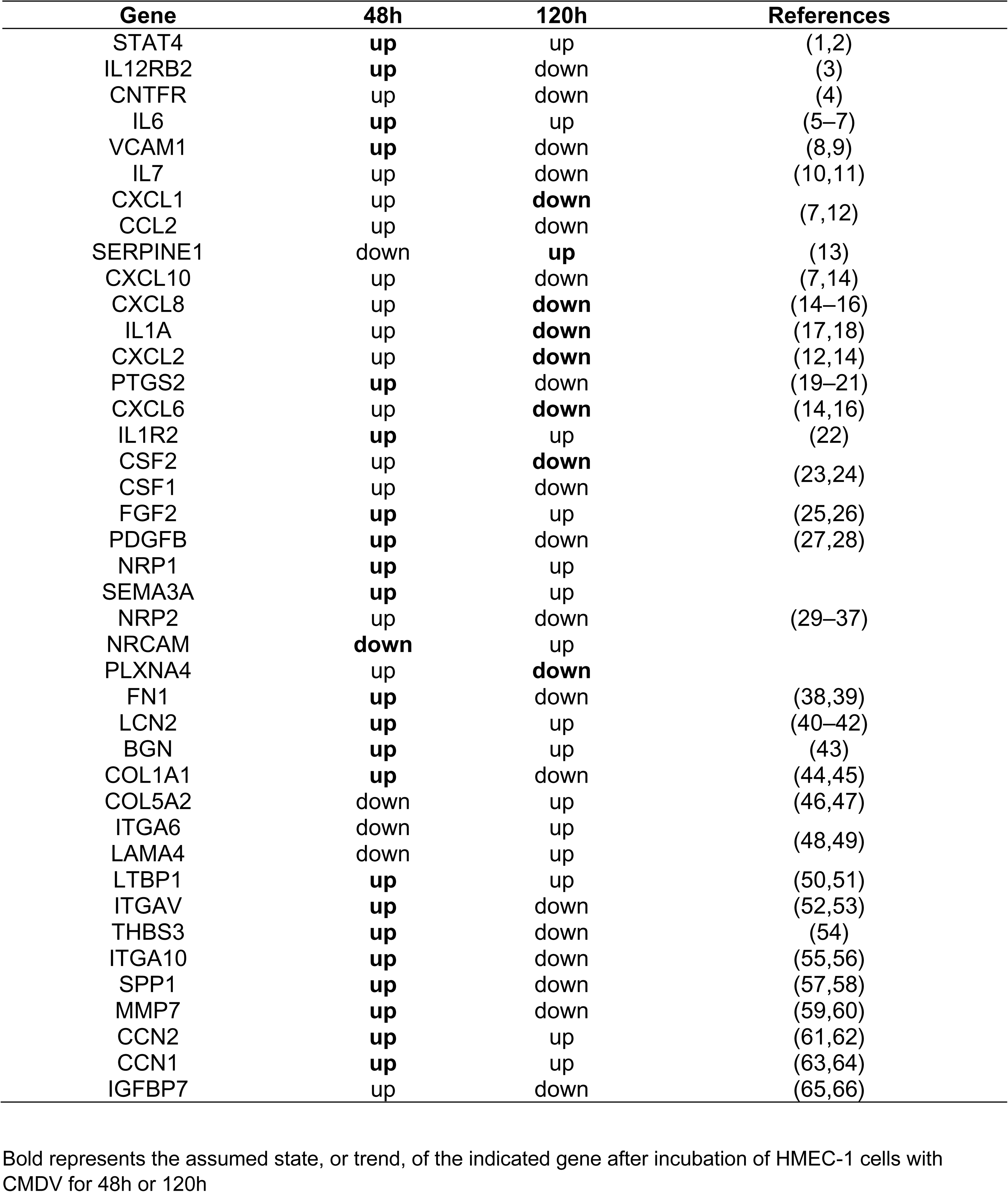
Genes State at 48h and 120h for CMDV model.

---

**Supplementary Table 3A.** Centralities analysis results.

| **Node** | **Degree** | **Betweenness** | **Closeness** | **Eigenvector** |
| --- | --- | --- | --- | --- |
| STAT4 | 5 | 0.5 | 0.00892857 | 0.19954602 |
| IL6 | 22 | 312.779131 | 0.01333333 | 1 |
| FN1 | 12 | 290.807504 | 0.01315789 | 0.32518526 |
| IL1A | 11 | 13.799026 | 0.01052632 | 0.80699843 |
| PTGS2 | 5 | 10.3804113 | 0.01020408 | 0.33312293 |
| CXCL1 | 11 | 73.8165584 | 0.01149425 | 0.69420125 |
| CSF2 | 7 | 61.875 | 0.01075269 | 0.4464607 |
| FGF2 | 8 | 224.071807 | 0.01162791 | 0.32409437 |
| VCAM1 | 5 | 10.0087662 | 0.01086957 | 0.36382442 |
| CXCL2 | 7 | 6.33522727 | 0.01 | 0.53868731 |
| CXCL10 | 8 | 13.8562951 | 0.01041667 | 0.56230585 |
| CXCL8 | 7 | 1.32900433 | 0.01020408 | 0.55143923 |
| SERPINE1 | 7 | 36.3041847 | 0.01075269 | 0.54202478 |
| CXCL6 | 3 | 0 | 0.00819672 | 0.2127898 |
| CCN2 | 7 | 39.5206349 | 0.01 | 0.20091607 |
| CCL2 | 10 | 7.54902597 | 0.01052632 | 0.71688373 |
| NRP1 | 3 | 144 | 0.00854701 | 0.03973524 |
| NRP2 | 4 | 57.5 | 0.00662252 | 0.00560052 |
| SEMA3A | 3 | 18.5 | 0.00657895 | 0.00553017 |
| BGN | 4 | 12.3768939 | 0.01075269 | 0.18431969 |
| ITGA10 | 2 | 1.5 | 0.00694444 | 0.01523884 |
| ITGA6 | 2 | 39 | 0.00884956 | 0.03909447 |
| CCN1 | 2 | 0 | 0.00724638 | 0.03769081 |
| SPP1 | 3 | 1.33333333 | 0.00900901 | 0.06382146 |
| ITGAV | 4 | 53.4791667 | 0.00909091 | 0.04859927 |
| MMP7 | 3 | 7.625 | 0.01052632 | 0.16464315 |
| LTBP1 | 5 | 15.7011905 | 0.00943396 | 0.11706129 |
| COL1A1 | 4 | 26.5541667 | 0.00909091 | 0.07996273 |
| IL12RB2 | 2 | 0 | 0.00884956 | 0.14218581 |
| CNTFR | 2 | 0 | 0.00884956 | 0.14218581 |
| IL7 | 2 | 0 | 0.00952381 | 0.17145335 |
| CSF1 | 1 | 0 | 0.00757576 | 0.05292033 |
| PLXNA4 | 2 | 0 | 0.00529101 | 0.00131935 |
| LCN2 | 2 | 0 | 0.01041667 | 0.1570782 |
| LAMA4 | 1 | 0 | 0.00657895 | 0.00463399 |
| THBS3 | 1 | 0 | 0.00671141 | 0.00576062 |
| IGFBP7 | 4 | 15.6541126 | 0.00943396 | 0.22287564 |
| IL1R2 | 2 | 0 | 0.00884956 | 0.21418897 |
| PDGFB | 1 | 0 | 0.008 | 0.03841588 |
| COL5A2 | 3 | 8.84356061 | 0.00917431 | 0.14985921 |
| NRCAM | 1 | 0 | 0.00526316 | 0.00066385 |

**Supplementary Table 3B.** Network Metrics.

| Supplementary Table 3B. Network Metrics |  |
| --- | --- |
| Metric | Value |
| Clustering Coefficient | 0.56290484 |
| Number of Communities | 6 |
| Diameter | 6 |
| Radius | 3 |
| Density | 0.12073171 |

**Supplementary Table 4.**
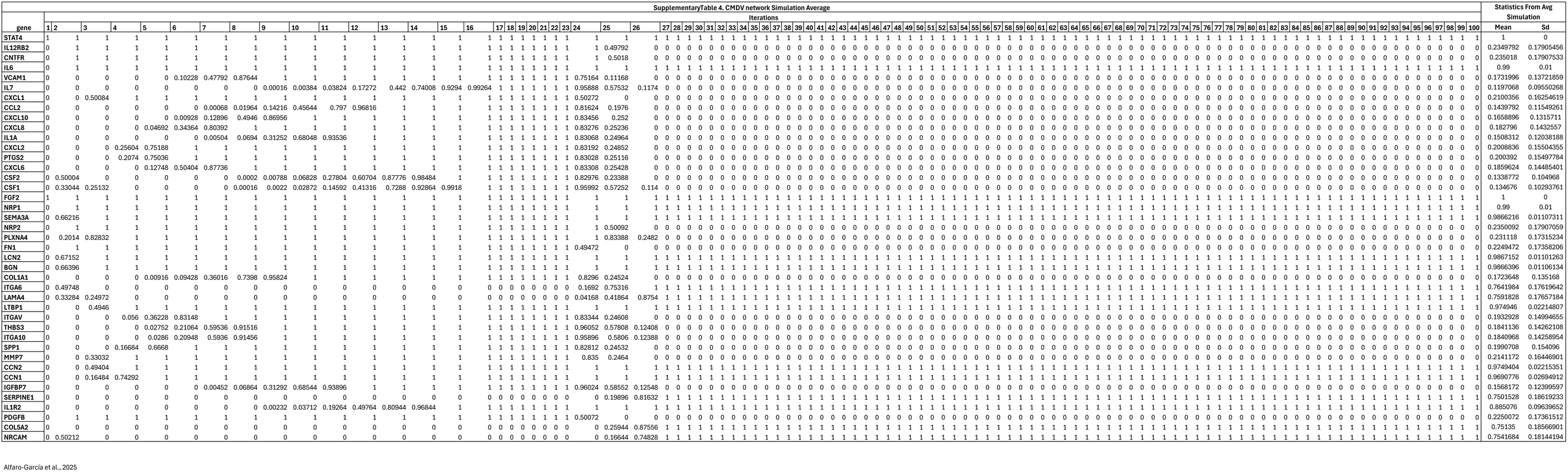
CMDV network Simulation Average.

**Supplementary table 5.**
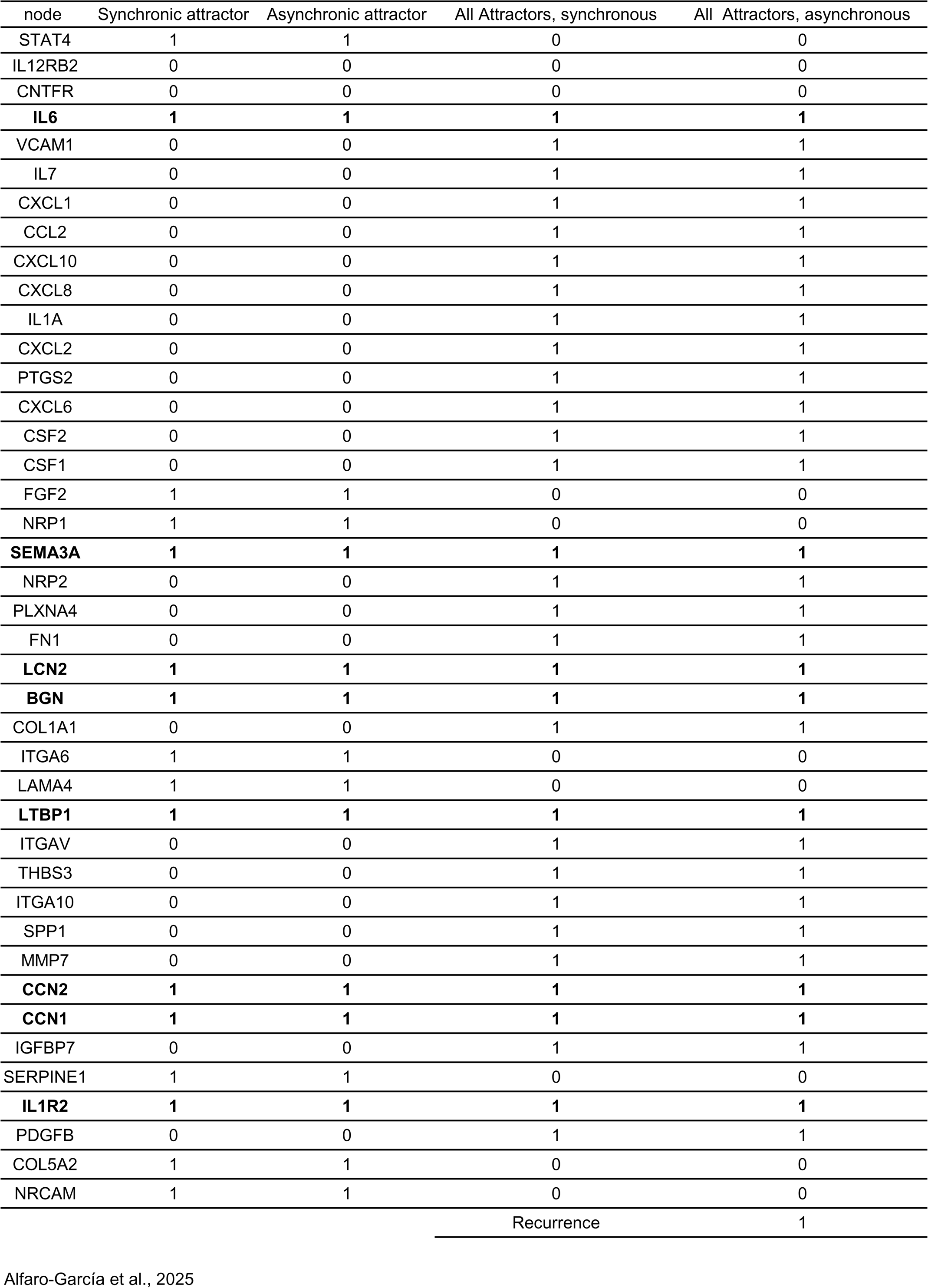
Attractors Analysis.

| node | Synchronic attractor | Asynchronic attractor | All Attractors, synchronous | All Attractors, asynchronous |
| --- | --- | --- | --- | --- |
| STAT4 | 1 | 1 | 0 | 0 |
| IL12RB2 | 0 | 0 | 0 | 0 |
| CNTFR | 0 | 0 | 0 | 0 |
| **IL6** | **1** | **1** | **1** | **1** |
| VCAM1 | 0 | 0 | 1 | 1 |
| IL7 | 0 | 0 | 1 | 1 |
| CXCL1 | 0 | 0 | 1 | 1 |
| CCL2 | 0 | 0 | 1 | 1 |
| CXCL10 | 0 | 0 | 1 | 1 |
| CXCL8 | 0 | 0 | 1 | 1 |
| IL1A | 0 | 0 | 1 | 1 |
| CXCL2 | 0 | 0 | 1 | 1 |
| PTGS2 | 0 | 0 | 1 | 1 |
| CXCL6 | 0 | 0 | 1 | 1 |
| CSF2 | 0 | 0 | 1 | 1 |
| CSF1 | 0 | 0 | 1 | 1 |
| FGF2 | 1 | 1 | 0 | 0 |
| NRP1 | 1 | 1 | 0 | 0 |
| **SEMA3A** | **1** | **1** | **1** | **1** |
| NRP2 | 0 | 0 | 1 | 1 |
| PLXNA4 | 0 | 0 | 1 | 1 |
| FN1 | 0 | 0 | 1 | 1 |
| **LCN2** | **1** | **1** | **1** | **1** |
| **BGN** | **1** | **1** | **1** | **1** |
| COL1A1 | 0 | 0 | 1 | 1 |
| ITGA6 | 1 | 1 | 0 | 0 |
| LAMA4 | 1 | 1 | 0 | 0 |
| **LTBP1** | **1** | **1** | **1** | **1** |
| ITGAV | 0 | 0 | 1 | 1 |
| THBS3 | 0 | 0 | 1 | 1 |
| ITGA10 | 0 | 0 | 1 | 1 |
| SPP1 | 0 | 0 | 1 | 1 |
| MMP7 | 0 | 0 | 1 | 1 |
| **CCN2** | **1** | **1** | **1** | **1** |
| **CCN1** | **1** | **1** | **1** | **1** |
| IGFBP7 | 0 | 0 | 1 | 1 |
| SERPINE1 | 1 | 1 | 0 | 0 |
| **IL1R2** | **1** | **1** | **1** | **1** |
| PDGFB | 0 | 0 | 1 | 1 |
| COL5A2 | 1 | 1 | 0 | 0 |
| NRCAM | 1 | 1 | 0 | 0 |
|  |  |  | Recurrence | 1 |

**Supplementary Table 6.**
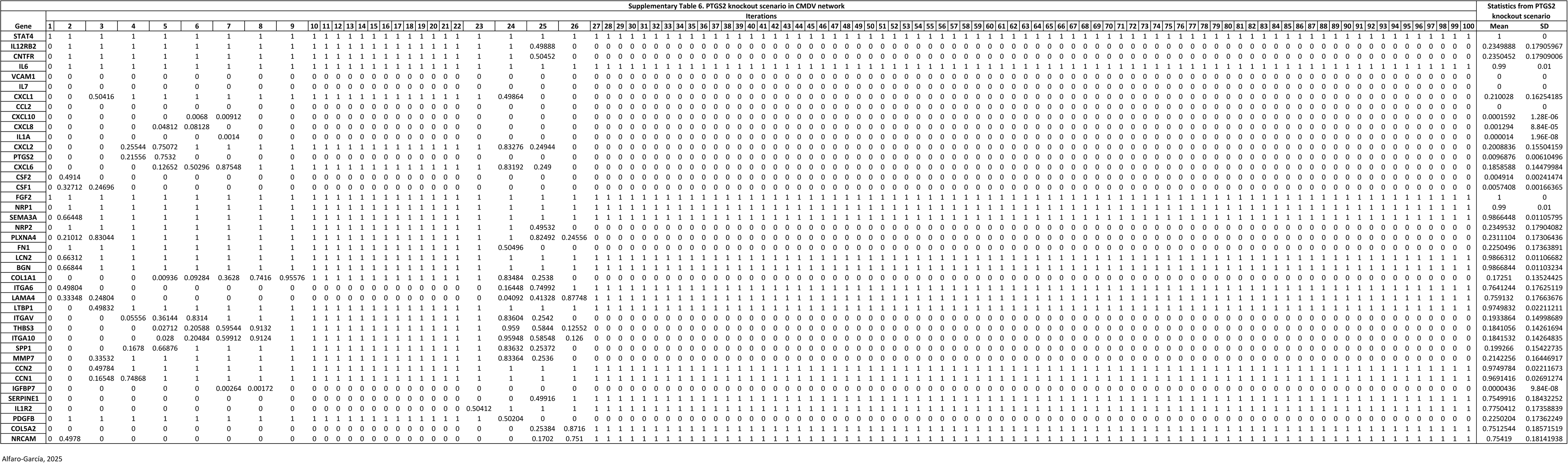
PTGS2 knockout scenario in CMDV network.

**Supplementary Table 7.**
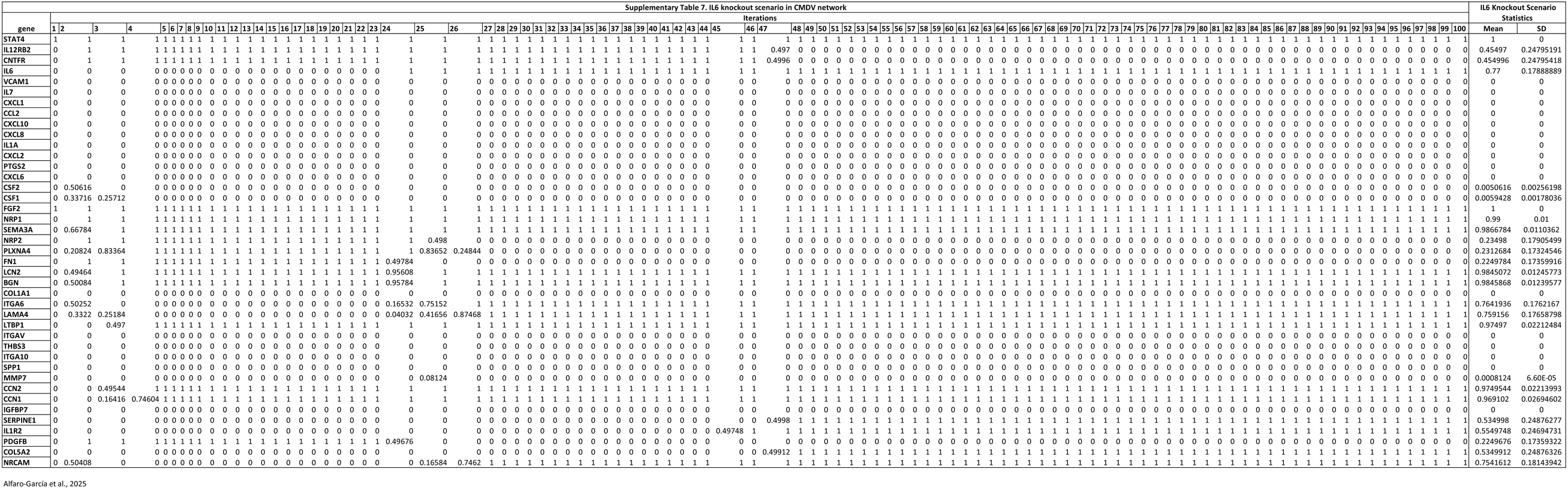
IL6 knockout scenario in CMDV network.

**Supplementary Table 8.**
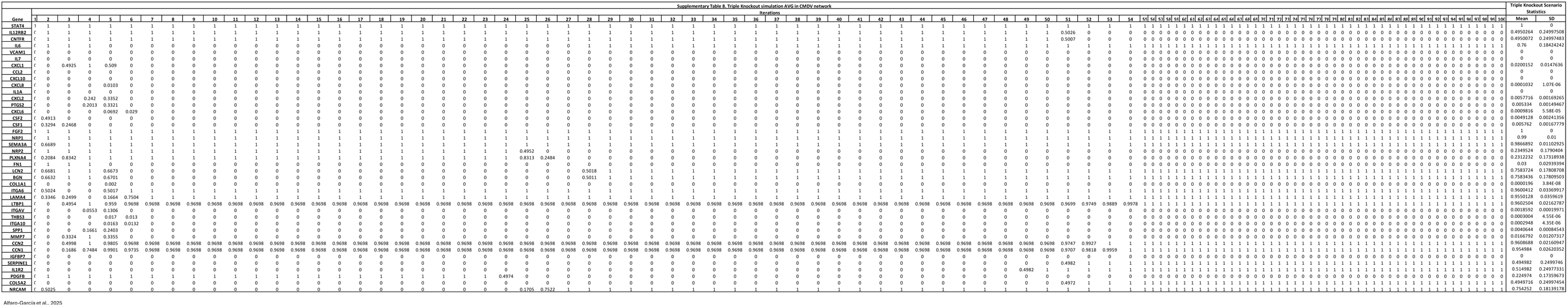
Triple Knockout simulation AVG in CMDV network.

**Supplementary Table 9.**
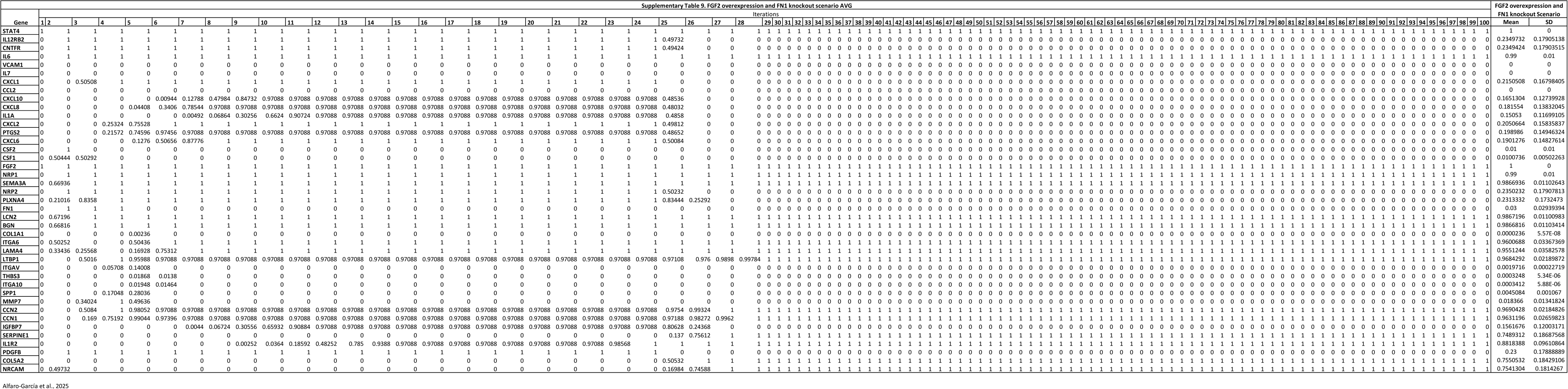
FGF2 overexpression and FN1 knockout scenario AVG.

## Bibliography

1. Brady OJ, Gething PW, Bhatt S, Messina JP, Brownstein JS, Hoen AG, et al. Refining the global spatial limits of dengue virus transmission by evidence-based consensus. PLoS Negl Trop Dis. 2012;6(8):e1760. Epub 2012/08/11. doi: 10.1371/journal.pntd.0001760. PubMed PMID: 22880140; PubMed Central PMCID: PMCPMC3413714.

2. Akinsulie OC, Idris I. Global re-emergence of dengue fever: The need for a rapid response and surveillance. The Microbe. 2024;4:100107. doi: 10.1016/j.microb.2024.100107.

3. Pan American Health Organization (PAHO); World Health Organization (WHO) Dengue Situation Report Region of the Americas. 2023. p. 2019.

4. Tejo AM, Hamasaki DT, Menezes LM, Ho YL. Severe dengue in the intensive care unit. J Intensive Med. 2024;4(1):16–33. Epub 2024/01/24. doi: 10.1016/j.jointm.2023.07.007. PubMed PMID: 38263966; PubMed Central PMCID: PMCPMC10800775.

5. Singh RK, Tiwari A, Satone PD, Priya T, Meshram RJ. Updates in the Management of Dengue Shock Syndrome: A Comprehensive Review. Cureus. 2023;15(10):e46713. Epub 2023/11/29. doi: 10.7759/cureus.46713. PubMed PMID: 38021722; PubMed Central PMCID: PMCPMC10631559.

6. Bhatt P, Sabeena SP, Varma M, Arunkumar G. Current Understanding of the Pathogenesis of Dengue Virus Infection. Curr Microbiol. 2021;78(1):17–32. Epub 2020/11/25. doi: 10.1007/s00284-020-02284-w. PubMed PMID: 33231723; PubMed Central PMCID: PMCPMC7815537.

7. Roy SK, Bhattacharjee S. Dengue virus: epidemiology, biology, and disease aetiology. Can J Microbiol. 2021;67(10):687–702. Epub 2021/06/26. doi: 10.1139/cjm-2020-0572. PubMed PMID: 34171205.

8. Niranjan R, Kishor S, Kumar A. Matrix metalloproteinases in the pathogenesis of dengue viral disease: Involvement of immune system and newer therapeutic strategies. J Med Virol. 2021;93(8):4629–37. Epub 2021/02/27. doi: 10.1002/jmv.26903. PubMed PMID: 33634515.

9. Guzman MG, Harris E. Dengue. Lancet. 2015;385(9966):453–65. Epub 2014/09/19. doi: 10.1016/S0140-6736(14)60572-9. PubMed PMID: 25230594.

10. Malavige GN, Ogg GS. Pathogenesis of vascular leak in dengue virus infection. Immunology. 2017;151(3):261–9. Epub 2017/04/25. doi: 10.1111/imm.12748. PubMed PMID: 28437586; PubMed Central PMCID: PMCPMC5461104.

11. Fiestas Solorzano VE, da Costa Faria NR, Dos Santos CF, Correa G, Cipitelli MDC, Dornelas Ribeiro M, et al. Different Profiles of Cytokines, Chemokines and Coagulation Mediators Associated with Severity in Brazilian Patients Infected with Dengue Virus. Viruses. 2021;13(9). Epub 2021/09/29. doi: 10.3390/v13091789. PubMed PMID: 34578370; PubMed Central PMCID: PMCPMC8473164.

12. Fiestas Solorzano VE, de Lima RC, de Azeredo EL. The Role of Growth Factors in the Pathogenesis of Dengue: A Scoping Review. Pathogens. 2022;11(10). Epub 2022/10/28. doi: 10.3390/pathogens11101179. PubMed PMID: 36297236; PubMed Central PMCID: PMCPMC9608673.

13. Cuartas-Lopez AM, Hernandez-Cuellar CE, Gallego-Gomez JC. Disentangling the role of PI3K/Akt, Rho GTPase and the actin cytoskeleton on dengue virus infection. Virus Res. 2018;256:153–65. Epub 2018/08/22. doi: 10.1016/j.virusres.2018.08.013. PubMed PMID: 30130602.

14. Escudero-Florez M, Torres-Hoyos D, Miranda-Brand Y, Boudreau RL, Gallego-Gomez JC, Vicente-Manzanares M. Dengue Virus Infection Alters Inter-Endothelial Junctions and Promotes Endothelial-Mesenchymal-Transition-Like Changes in Human Microvascular Endothelial Cells. Viruses. 2023;15(7). Epub 2023/07/29. doi: 10.3390/v15071437. PubMed PMID: 37515125; PubMed Central PMCID: PMCPMC10386726.

15. Alvarez-Diaz DA, Gutierrez-Diaz AA, Orozco-Garcia E, Puerta-Gonzalez A, Bermudez-Santana CI, Gallego-Gomez JC. Dengue virus potentially promotes migratory responses on endothelial cells by enhancing pro-migratory soluble factors and miRNAs. Virus Res. 2019;259:68–76. Epub 2018/10/28. doi: 10.1016/j.virusres.2018.10.018. PubMed PMID: 30367889.

16. Alfaro-Garcia JP, Orozco-Castano CA, Sanchez-Rendon JA, Casanova-Yepes HF, Vicente-Manzanares M, Gallego-Gomez JC. Characterization of the Temporal Dynamics of the Endothelial-Mesenchymal-like Transition Induced by Soluble Factors from Dengue Virus Infection in Microvascular Endothelial Cells. Int J Mol Sci. 2025;26(5). Epub 2025/03/13. doi: 10.3390/ijms26052139. PubMed PMID: 40076764; PubMed Central PMCID: PMCPMC11900998.

17. Piera-Velazquez S, Jimenez SA. Endothelial to Mesenchymal Transition: Role in Physiology and in the Pathogenesis of Human Diseases. Physiol Rev. 2019;99(2):1281–324. Epub 2019/03/14. doi: 10.1152/physrev.00021.2018. PubMed PMID: 30864875; PubMed Central PMCID: PMCPMC6734087.

18. Huang X, Pan L, Pu H, Wang Y, Zhang X, Li C, et al. Loss of caveolin-1 promotes endothelial-mesenchymal transition during sepsis: a membrane proteomic study. Int J Mol Med. 2013;32(3):585–92. Epub 2013/07/10. doi: 10.3892/ijmm.2013.1432. PubMed PMID: 23836408.

19. Stasi A, Intini A, Divella C, Franzin R, Montemurno E, Grandaliano G, et al. Emerging role of Lipopolysaccharide binding protein in sepsis-induced acute kidney injury. Nephrol Dial Transplant. 2017;32(1):24–31. Epub 2016/07/09. doi: 10.1093/ndt/gfw250. PubMed PMID: 27387474.

20. Feng J, Li K, Xie F, Han L, Wu Y. IL-35 ameliorates lipopolysaccharide-induced endothelial dysfunction by inhibiting endothelial-to-mesenchymal transition. Int Immunopharmacol. 2024;129:111567. Epub 2024/02/10. doi: 10.1016/j.intimp.2024.111567. PubMed PMID: 38335651.

21. Naipauer J, Mesri EA. The Kaposi’s sarcoma progenitor enigma: KSHV-induced MEndT-EndMT axis. Trends Mol Med. 2023;29(3):188–200. Epub 2023/01/13. doi: 10.1016/j.molmed.2022.12.003. PubMed PMID: 36635149; PubMed Central PMCID: PMCPMC9957928.

22. Ciszewski WM, Wozniak LA, Sobierajska K. Diverse roles of SARS-CoV-2 Spike and Nucleocapsid proteins in EndMT stimulation through the TGF-beta-MRTF axis inhibited by aspirin. Cell Commun Signal. 2024;22(1):296. Epub 2024/05/29. doi: 10.1186/s12964-024-01665-z. PubMed PMID: 38807115; PubMed Central PMCID: PMCPMC11134719.

23. Ro Y-T, Patterson JL. Transcriptional induction of TGF-β1 and endothelial-to-mesenchymal transition cell markers in human umbilical vein endothelial cells by Ebola virus infection. Genes & Genomics. 2022;44(12):1499–507. doi: 10.1007/s13258-022-01333-x.

24. Bischoff J. Endothelial-to-Mesenchymal Transition. Circ Res. 2019;124(8):1163–5. Epub 2019/04/12. doi: 10.1161/CIRCRESAHA.119.314813. PubMed PMID: 30973806; PubMed Central PMCID: PMCPMC6540806.

25. Dejana E, Lampugnani MG. Endothelial cell transitions. Science. 2018;362(6416):746–7. Epub 2018/11/18. doi: 10.1126/science.aas9432. PubMed PMID: 30442789.

26. Islam S, Bostrom KI, Di Carlo D, Simmons CA, Tintut Y, Yao Y, et al. The Mechanobiology of Endothelial-to-Mesenchymal Transition in Cardiovascular Disease. Frontiers in physiology. 2021;12:734215. Epub 2021/09/28. doi: 10.3389/fphys.2021.734215. PubMed PMID: 34566697; PubMed Central PMCID: PMCPMC8458763.

27. Dalrymple NA, Mackow ER. Roles for endothelial cells in dengue virus infection. Adv Virol. 2012;2012:840654. Epub 2012/09/07. doi: 10.1155/2012/840654. PubMed PMID: 22952474; PubMed Central PMCID: PMCPMC3431041.

28. Rathi KR, Arora MM, Sahai K, Tripathi S, Singh SP, Raman DK, et al. Autopsy findings in fatal dengue haemorrhagic fever – 06 Cases. Med J Armed Forces India. 2013;69(3):254–9. Epub 2014/03/07. doi: 10.1016/j.mjafi.2012.08.021. PubMed PMID: 24600119; PubMed Central PMCID: PMCPMC3862602.

29. Povoa TF, Alves AM, Oliveira CA, Nuovo GJ, Chagas VL, Paes MV. The pathology of severe dengue in multiple organs of human fatal cases: histopathology, ultrastructure and virus replication. PLoS One. 2014;9(4):e83386. Epub 2014/04/17. doi: 10.1371/journal.pone.0083386. PubMed PMID: 24736395; PubMed Central PMCID: PMCPMC3987999.

30. Cooley BC, Nevado J, Mellad J, Yang D, St Hilaire C, Negro A, et al. TGF-beta signaling mediates endothelial-to-mesenchymal transition (EndMT) during vein graft remodeling. Sci Transl Med. 2014;6(227):227ra34. Epub 2014/03/14. doi: 10.1126/scitranslmed.3006927. PubMed PMID: 24622514; PubMed Central PMCID: PMCPMC4181409.

31. Wang X, Bleher R, Wang L, Garcia JGN, Dudek SM, Shekhawat GS, et al. Imatinib Alters Agonists-mediated Cytoskeletal Biomechanics in Lung Endothelium. Sci Rep. 2017;7(1):14152. Epub 2017/10/28. doi: 10.1038/s41598-017-14722-0. PubMed PMID: 29075042; PubMed Central PMCID: PMCPMC5658337.

32. Teixeira GS, Andrade AA, Torres LR, Couto-Lima D, Moreira OC, Abreu R, et al. Suppression of TGF-beta/Smad2 signaling by GW788388 enhances DENV-2 clearance in macrophages. J Med Virol. 2022;94(9):4359–68. Epub 2022/05/21. doi: 10.1002/jmv.27879. PubMed PMID: 35596058; PubMed Central PMCID: PMCPMC9544077.

33. Irurzun-Arana I, Pastor JM, Trocóniz IF, Gómez-Mantilla JD. Advanced Boolean modeling of biological networks applied to systems pharmacology. Bioinformatics. 2017;33(7):1040–8. doi: 10.1093/bioinformatics/btw747.

34. Knecht E, Aguado C, Carcel J, Esteban I, Esteve JM, Ghislat G, et al. Intracellular protein degradation in mammalian cells: recent developments. Cellular and molecular life sciences: CMLS. 2009;66(15):2427–43. Epub 2009/04/29. doi: 10.1007/s00018-009-0030-6. PubMed PMID: 19399586; PubMed Central PMCID: PMCPMC11115841.

35. Mellenius H. Speed and accuracy in transcription and translation. Uppsala 2015.

36. Pickett JR, Wu Y, Zacchi LF, Ta HT. Targeting endothelial vascular cell adhesion molecule-1 in atherosclerosis: drug discovery and development of vascular cell adhesion molecule-1-directed novel therapeutics. Cardiovasc Res. 2023;119(13):2278–93. Epub 2023/08/18. doi: 10.1093/cvr/cvad130. PubMed PMID: 37595265; PubMed Central PMCID: PMCPMC10597632.

37. Koizumi M, King N, Lobb R, Benjamin C, Podolsky DK. Expression of vascular adhesion molecules in inflammatory bowel disease. Gastroenterology. 1992;103(3):840–7. Epub 1992/09/01. doi: 10.1016/0016-5085(92)90015-q. PubMed PMID: 1379955.

38. Youssef KK, Nieto MA. Epithelial-mesenchymal transition in tissue repair and degeneration. Nature reviews Molecular cell biology. 2024;25(9):720–39. Epub 2024/04/30. doi: 10.1038/s41580-024-00733-z. PubMed PMID: 38684869.

39. Gupta A, Rijhwani P, Pahadia MR, Kalia A, Choudhary S, Bansal DP, et al. Prevalence of Dengue Serotypes and Its Correlation With the Laboratory Profile at a Tertiary Care Hospital in Northwestern India. Cureus. 2021;13(5):e15029. Epub 2021/06/18. doi: 10.7759/cureus.15029. PubMed PMID: 34136322; PubMed Central PMCID: PMCPMC8199925.

40. Senaratne UTN, Murugananthan K, Sirisena P, Carr JM, Noordeen F. Dengue virus co-infections with multiple serotypes do not result in a different clinical outcome compared to mono-infections. Epidemiol Infect. 2020;148:e119. Epub 2020/07/01. doi: 10.1017/S0950268820000229. PubMed PMID: 32594967; PubMed Central PMCID: PMCPMC7325333.

41. Suppiah J, Ching SM, Amin-Nordin S, Mat-Nor LA, Ahmad-Najimudin NA, Low GK, et al. Clinical manifestations of dengue in relation to dengue serotype and genotype in Malaysia: A retrospective observational study. PLoS Negl Trop Dis. 2018;12(9):e0006817. Epub 2018/09/19. doi: 10.1371/journal.pntd.0006817. PubMed PMID: 30226880; PubMed Central PMCID: PMCPMC6161924.

42. Tsai JJ, Chang K, Chen CH, Liao CL, Chen LJ, Tsai YY, et al. Dengue virus serotype did not contribute to clinical severity or mortality in Taiwan’s largest dengue outbreak in 2015. Eur J Med Res. 2023;28(1):482. Epub 2023/11/07. doi: 10.1186/s40001-023-01454-3. PubMed PMID: 37932817; PubMed Central PMCID: PMCPMC10626727.

43. Morgan RN, Ismail NSM, Alshahrani MY, Aboshanab KM. Multi-epitope peptide vaccines targeting dengue virus serotype 2 created via immunoinformatic analysis. Sci Rep. 2024;14(1):17645. Epub 2024/08/01. doi: 10.1038/s41598-024-67553-1. PubMed PMID: 39085250; PubMed Central PMCID: PMCPMC11291903.

44. Sitthisuwannakul K, Sukthai R, Zhu Z, Nagashima K, Chattrairat K, Phanthanawiboon S, et al. Urinary dengue NS1 detection on Au-decorated ZnO nanowire platform. Biosens Bioelectron. 2024;254:116218. Epub 2024/03/23. doi: 10.1016/j.bios.2024.116218. PubMed PMID: 38518559.

45. Puerta-Guardo H, Glasner DR, Espinosa DA, Biering SB, Patana M, Ratnasiri K, et al. Flavivirus NS1 Triggers Tissue-Specific Vascular Endothelial Dysfunction Reflecting Disease Tropism. Cell Rep. 2019;26(6):1598–613 e8. Epub 2019/02/07. doi: 10.1016/j.celrep.2019.01.036. PubMed PMID: 30726741; PubMed Central PMCID: PMCPMC6934102.

46. Glasner DR, Ratnasiri K, Puerta-Guardo H, Espinosa DA, Beatty PR, Harris E. Dengue virus NS1 cytokine-independent vascular leak is dependent on endothelial glycocalyx components. PLoS Pathog. 2017;13(11):e1006673. Epub 2017/11/10. doi: 10.1371/journal.ppat.1006673. PubMed PMID: 29121099; PubMed Central PMCID: PMCPMC5679539.

47. Subauste MC, Pertz O, Adamson ED, Turner CE, Junger S, Hahn KM. Vinculin modulation of paxillin-FAK interactions regulates ERK to control survival and motility. The Journal of cell biology. 2004;165(3):371–81. Epub 2004/05/13. doi: 10.1083/jcb.200308011. PubMed PMID: 15138291; PubMed Central PMCID: PMCPMC2172187.

48. Tsubouchi A, Sakakura J, Yagi R, Mazaki Y, Schaefer E, Yano H, et al. Localized suppression of RhoA activity by Tyr31/118-phosphorylated paxillin in cell adhesion and migration. The Journal of cell biology. 2002;159(4):673–83. Epub 2002/11/26. doi: 10.1083/jcb.200202117. PubMed PMID: 12446743; PubMed Central PMCID: PMCPMC2173105.

49. Parsons SA, Sharma R, Roccamatisi DL, Zhang H, Petri B, Kubes P, et al. Endothelial paxillin and focal adhesion kinase (FAK) play a critical role in neutrophil transmigration. Eur J Immunol. 2012;42(2):436–46. Epub 2011/11/19. doi: 10.1002/eji.201041303. PubMed PMID: 22095445.

50. Meng ZZ, Liu W, Xia Y, Yin HM, Zhang CY, Su D, et al. The pro-inflammatory signalling regulator Stat4 promotes vasculogenesis of great vessels derived from endothelial precursors. Nat Commun. 2017;8:14640. Epub 2017/03/04. doi: 10.1038/ncomms14640. PubMed PMID: 28256502; PubMed Central PMCID: PMCPMC5338034.

51. Xu H, Pumiglia K, LaFlamme SE. Laminin-511 and alpha6 integrins regulate the expression of CXCR4 to promote endothelial morphogenesis. J Cell Sci. 2020;133(11). Epub 2020/05/16. doi: 10.1242/jcs.246595. PubMed PMID: 32409567.

52. Beguin EP, Janssen EFJ, Hoogenboezem M, Meijer AB, Hoogendijk AJ, van den Biggelaar M. Flow-induced Reorganization of Laminin-integrin Networks Within the Endothelial Basement Membrane Uncovered by Proteomics. Mol Cell Proteomics. 2020;19(7):1179–92. Epub 2020/04/26. doi: 10.1074/mcp.RA120.001964. PubMed PMID: 32332107; PubMed Central PMCID: PMCPMC7338090.

53. Pussell BA, Peake PW, Brown MA, Charlesworth JA. Human fibronectin metabolism. J Clin Invest. 1985;76(1):143–8. Epub 1985/07/01. doi: 10.1172/JCI111937. PubMed PMID: 4019773; PubMed Central PMCID: PMCPMC423729.

54. Qi Y, Qadir MMF, Hastreiter AA, Fock RA, Machi JF, Morales AA, et al. Endothelial c-Myc knockout enhances diet-induced liver inflammation and fibrosis. FASEB journal: official publication of the Federation of American Societies for Experimental Biology. 2022;36(1):e22077. Epub 2021/12/09. doi: 10.1096/fj.202101086R. PubMed PMID: 34878671; PubMed Central PMCID: PMCPMC11367571.

55. See KC. Dengue-Associated Hemophagocytic Lymphohistiocytosis: A Narrative Review of Its Identification and Treatment. Pathogens. 2024;13(4). Epub 2024/04/26. doi: 10.3390/pathogens13040332. PubMed PMID: 38668287; PubMed Central PMCID: PMCPMC11053942.

56. Weinstein N, Mendoza L, Álvarez-Buylla ER. A Computational Model of the Endothelial to Mesenchymal Transition. Frontiers in genetics. 2020;11:40. Epub 2020/04/01. doi: 10.3389/fgene.2020.00040. PubMed PMID: 32226439; PubMed Central PMCID: PMCPmc7080988.

57. Aguilar-Briseno JA, Moser J, Rodenhuis-Zybert IA. Understanding immunopathology of severe dengue: lessons learnt from sepsis. Curr Opin Virol. 2020;43:41–9. Epub 2020/09/09. doi: 10.1016/j.coviro.2020.07.010. PubMed PMID: 32896675.

58. Zhang YY, Ning BT. Signaling pathways and intervention therapies in sepsis. Signal Transduct Target Ther. 2021;6(1):407. Epub 2021/11/27. doi: 10.1038/s41392-021-00816-9. PubMed PMID: 34824200; PubMed Central PMCID: PMCPMC8613465.

59. Li Y, Zhao J, Yin Y, Li K, Zhang C, Zheng Y. The Role of IL-6 in Fibrotic Diseases: Molecular and Cellular Mechanisms. Int J Biol Sci. 2022;18(14):5405–14. Epub 2022/09/24. doi: 10.7150/ijbs.75876. PubMed PMID: 36147459; PubMed Central PMCID: PMCPMC9461670.

60. Takagaki Y, Lee SM, Dongqing Z, Kitada M, Kanasaki K, Koya D. Endothelial autophagy deficiency induces IL6 – dependent endothelial mesenchymal transition and organ fibrosis. Autophagy. 2020;16(10):1905–14. Epub 2020/01/23. doi: 10.1080/15548627.2020.1713641. PubMed PMID: 31965901; PubMed Central PMCID: PMCPMC8386622.

61. Mahmoud MM, Kim HR, Xing R, Hsiao S, Mammoto A, Chen J, et al. TWIST1 Integrates Endothelial Responses to Flow in Vascular Dysfunction and Atherosclerosis. Circ Res. 2016;119(3):450–62. Epub 2016/06/02. doi: 10.1161/CIRCRESAHA.116.308870. PubMed PMID: 27245171; PubMed Central PMCID: PMCPMC4959828.

62. Martinez-Gutierrez M, Correa-Londono LA, Castellanos JE, Gallego-Gomez JC, Osorio JE. Lovastatin delays infection and increases survival rates in AG129 mice infected with dengue virus serotype 2. PLoS One. 2014;9(2):e87412. Epub 2014/03/04. doi: 10.1371/journal.pone.0087412. PubMed PMID: 24586275; PubMed Central PMCID: PMCPMC3931612.

63. Bolger AM, Lohse M, Usadel B. Trimmomatic: a flexible trimmer for Illumina sequence data. Bioinformatics. 2014;30(15):2114–20. Epub 2014/04/04. doi: 10.1093/bioinformatics/btu170. PubMed PMID: 24695404; PubMed Central PMCID: PMCPMC4103590.

64. Andrews S. FastQC: A Quality Control Tool for High Throughput Sequence Data. Babraham Institute: https://www.bioinformatics.babraham.ac.uk/projects/fastqc/; 2019.

65. Dobin A, Davis CA, Schlesinger F, Drenkow J, Zaleski C, Jha S, et al. STAR: ultrafast universal RNA-seq aligner. Bioinformatics. 2013;29(1):15–21. Epub 2012/10/30. doi: 10.1093/bioinformatics/bts635. PubMed PMID: 23104886; PubMed Central PMCID: PMCPMC3530905.

66. Liao Y, Smyth GK, Shi W. featureCounts: an efficient general purpose program for assigning sequence reads to genomic features. Bioinformatics. 2014;30(7):923–30. Epub 2013/11/15. doi: 10.1093/bioinformatics/btt656. PubMed PMID: 24227677.

67. Mudge JM, Carbonell-Sala S, Diekhans M, Martinez JG, Hunt T, Jungreis I, et al. GENCODE 2025: reference gene annotation for human and mouse. Nucleic Acids Res. 2025;53(D1):D966-D75. Epub 2024/11/20. doi: 10.1093/nar/gkae1078. PubMed PMID: 39565199; PubMed Central PMCID: PMCPMC11701607.

68. Love MI, Huber W, Anders S. Moderated estimation of fold change and dispersion for RNA-seq data with DESeq2. Genome Biol. 2014;15(12):550. Epub 2014/12/18. doi: 10.1186/s13059-014-0550-8. PubMed PMID: 25516281; PubMed Central PMCID: PMCPMC4302049.

69. Subramanian A, Tamayo P, Mootha VK, Mukherjee S, Ebert BL, Gillette MA, et al. Gene set enrichment analysis: a knowledge-based approach for interpreting genome-wide expression profiles. Proc Natl Acad Sci U S A. 2005;102(43):15545–50. Epub 2005/10/04. doi: 10.1073/pnas.0506580102. PubMed PMID: 16199517; PubMed Central PMCID: PMCPMC1239896.

70. Coebergh van den Braak RRJ, Sieuwerts AM, Lalmahomed ZS, Smid M, Wilting SM, Bril SI, et al. Confirmation of a metastasis-specific microRNA signature in primary colon cancer. Sci Rep. 2018;8(1):5242. Epub 2018/03/29. doi: 10.1038/s41598-018-22532-1. PubMed PMID: 29588449; PubMed Central PMCID: PMCPMC5869672.

71. Wilkinson DJ. Stochastic Modelling for Systems Biology Third Edition. Lin X, Singh M, Britton NF, Tramontano A, Schneider MV, Mulder N, editors. Boca Raton, FL: CRC Press, Taylor & Francis; 2018.

72. Alfaro-Garcia JP, Granados-Alzate MC, Vicente-Manzanares M, Gallego-Gomez JC. An Integrated View of Virus-Triggered Cellular Plasticity Using Boolean Networks. Cells. 2021;10(11). Epub 2021/11/28. doi: 10.3390/cells10112863. PubMed PMID: 34831086; PubMed Central PMCID: PMCPMC8616224.

73. Schwab JD, Kuhlwein SD, Ikonomi N, Kuhl M, Kestler HA. Concepts in Boolean network modeling: What do they all mean? Comput Struct Biotechnol J. 2020;18:571–82. Epub 2020/04/08. doi: 10.1016/j.csbj.2020.03.001. PubMed PMID: 32257043; PubMed Central PMCID: PMCPMC7096748.

## Supplementary bibliography

1. Meng ZZ, Liu W, Xia Y, Yin HM, Zhang CY, Su D, et al. The pro-inflammatory signalling regulator Stat4 promotes vasculogenesis of great vessels derived from endothelial precursors. Nat Commun [Internet]. 2017;8:1–12. Available from: 10.1038/ncomms14640

2. Nguyen HN, Noss EH, Mizoguchi F, Huppertz C, Wei KS, Watts GFM, et al. Autocrine Loop Involving IL-6 Family Member LIF, LIF Receptor, and STAT4 Drives Sustained Fibroblast Production of Inflammatory Mediators. Immunity [Internet]. 2017;46(2):220–32. Available from: 10.1016/j.immuni.2017.01.004

3. Yao BB, Niu P, Surowy CS, Faltynek CR. Direct interaction of STAT4 with the IL-12 receptor. Arch Biochem Biophys. 1999;368(1):147–55.

4. Zhang L, Xiang Y, Cao C, Tan J, Li F, Yang X. Ciliary neurotrophic factor promotes the development of homocysteine-induced vascular endothelial injury through inflammation mediated by the JAK2/STAT3 signaling pathway. Exp Cell Res [Internet]. 2024;440(1):114103. Available from: https://www.sciencedirect.com/science/article/pii/S0014482724001940

5. Vachher H, Metgud T, Srikanth B. IL-6 Levels in Prediction of Severity of Dengue Fever. Indian J Pediatr. 2023;90(5):518.

6. Jovanović M, Vićovac L. Interleukin-6 Stimulates Cell Migration, Invasion and Integrin Expression in HTR-8/SVneo Cell Line. Placenta [Internet]. 2009;30(4):320–8. Available from: 10.1016/j.placenta.2009.01.013

7. Jusof FF, Lim CK, Aziz FN, Soe HJ, Raju CS, Sekaran SD, et al. The Cytokines CXCL10 and CCL2 and the Kynurenine Metabolite Anthranilic Acid Accurately Predict Patients at Risk of Developing Dengue with Warning Signs. J Infect Dis [Internet]. 2022;226(11):1964–73. Available from: 10.1093/infdis/jiac273

8. Cook-Mills JM, Marchese ME, Abdala-Valencia H. Vascular cell adhesion molecule-1 expression and signaling during disease: Regulation by reactive oxygen species and antioxidants. Antioxidants Redox Signal. 2011;15(6):1607–38.

9. Pickett JR, Wu Y, Zacchi LF, Ta HT. Targeting endothelial vascular cell adhesion molecule-1 in atherosclerosis: drug discovery and development of vascular cell adhesion molecule-1-directed novel therapeutics. Cardiovasc Res [Internet]. 2023;119(13):2278–93. Available from: 10.1093/cvr/cvad130

10. Winer H, Rodrigues GOL, Hixon JA, Aiello FB, Hsu TC, Wachter BT, et al. IL-7: Comprehensive review. Cytokine [Internet]. 2022;160:156049. Available from: https://www.sciencedirect.com/science/article/pii/S1043466622002587

11. Ariel A, Hershkoviz R, Cahalon L, Williams DE, Akiyama SK, Yamada KM, et al. Induction of T cell adhesion to extracellular matrix or endothelial cell ligands by soluble or matrix-bound interleukin-7. Eur J Immunol. 1997;27(10):2562–70.

12. Boro M, Balaji KN. CXCL1 and CXCL2 Regulate NLRP3 Inflammasome Activation via G-Protein–Coupled Receptor CXCR2. J Immunol. 2017;199(5):1660–71.

13. Simone TM, Higgins CE, Czekay RP, Law BK, Higgins SP, Archambeault J, et al. SERPINE1: A Molecular Switch in the Proliferation-Migration Dichotomy in Wound-“Activated” Keratinocytes. Adv Wound Care. 2014;3(3):281–90.

14. Zhou C, Gao Y, Ding P, Wu T, Ji G. The role of CXCL family members in different diseases. Cell Death Discov. 2023;9(1):1–12.

15. Ji HZ, Chen L, Ren M, Li S, Liu TY, Chen HJ, et al. CXCL8 Promotes Endothelial-to-Mesenchymal Transition of Endothelial Cells and Protects Cells from Erastin-Induced Ferroptosis via CXCR2-Mediated Activation of the NF-κB Signaling Pathway. Pharmaceuticals. 2023;16(9).

16. Cambier S, Gouwy M, Proost P. The chemokines CXCL8 and CXCL12: molecular and functional properties, role in disease and efforts towards pharmacological intervention. Cell Mol Immunol. 2023;20(3):217–51.

17. Liu X, Zhang H, He S, Mu X, Hu G, Dong H. Endothelial-Derived Interleukin-1α Activates Innate Immunity by Promoting the Bactericidal Activity of Transendothelial Neutrophils. Front Cell Dev Biol. 2020;8(July):1–9.

18. Cavalli G, Colafrancesco S, Emmi G, Imazio M, Lopalco G, Maggio MC, et al. Interleukin 1α: a comprehensive review on the role of IL-1α in the pathogenesis and treatment of autoimmune and inflammatory diseases. Autoimmun Rev [Internet]. 2021;20(3):102763. Available from: 10.1016/j.autrev.2021.102763

19. Kulesza A, Paczek L, Burdzinska A. The Role of COX-2 and PGE2 in the Regulation of Immunomodulation and Other Functions of Mesenchymal Stromal Cells. Biomedicines. 2023;11(2).

20. Neil JR, Johnson KM, Nemenoff RA, Schiemann WP. Cox-2 inactivates Smad signaling and enhances EMT stimulated by TGF-β through a PGE2-dependent mechanisms. Carcinogenesis. 2008;29(11):2227–35.

21. Lin CK, Tseng CK, Wu YH, Liaw CC, Lin CY, Huang CH, et al. Cyclooxygenase-2 facilitates dengue virus replication and serves as a potential target for developing antiviral agents. Sci Rep. 2017;7(August 2016):1–15.

22. Peters VA, Joesting JJ, Freund GG. IL-1 receptor 2 (IL-1R2) and its role in immune regulation. Brain Behav Immun [Internet]. 2013;32:1–8. Available from: 10.1016/j.bbi.2012.11.006

23. Park SR, Cho A, Kim JW, Lee HY, Hong IS. A Novel Endogenous Damage Signal, CSF-2, Activates Multiple Beneficial Functions of Adipose Tissue-Derived Mesenchymal Stem Cells. Mol Ther [Internet]. 2019;27(6):1087–100. Available from: 10.1016/j.ymthe.2019.03.010

24. Park KW, Kwon YW, Cho HJ, Shin JI, Kim YJ, Lee SE, et al. G-CSF exerts dual effects on endothelial cells-Opposing actions of direct eNOS induction versus indirect CRP elevation. J Mol Cell Cardiol [Internet]. 2008;45(5):670–8. Available from: 10.1016/j.yjmcc.2008.07.002

25. Tan Y, Qiao Y, Chen Z, Liu J, Guo Y, Tran T, et al. FGF2, an Immunomodulatory Factor in Asthma and Chronic Obstructive Pulmonary Disease (COPD). Front Cell Dev Biol. 2020;8(April):1–12.

26. Piera-Velazquez S, Jimenez SA. Endothelial to mesenchymal transition: Role in physiology and in the pathogenesis of human diseases. Physiol Rev. 2019;99(2):1281–324.

27. Eng E, Ballermann BJ. Diminished NF-κB activation and PDGF-B expression in glomerular endothelial cells subjected to chronic shear stress. Microvasc Res. 2003;65(3):137–44.

28. D’Amore PA, Sakurai MK. Angiogenesis, Angiogenic Growth Factors and Development Factors. Encycl Respir Med Vol 1-4. 2006;1–4:V1-110-V1-115.

29. Sharma S, Ehrlich M, Zhang M, Blobe GC, Henis YI. NRP1 interacts with endoglin and VEGFR2 to modulate VEGF signaling and endothelial cell sprouting. Commun Biol. 2024;7(1):1–15.

30. Issitt T, Bosseboeuf E, De Winter N, Dufton N, Gestri G, Senatore V, et al. Neuropilin-1 Controls Endothelial Homeostasis by Regulating Mitochondrial Function and Iron-Dependent Oxidative Stress. iScience [Internet]. 2019;11:205–23. Available from: 10.1016/j.isci.2018.12.005

31. Chikh A, Raimondi C. Endothelial Neuropilin-1: a multifaced signal transducer with an emerging role in inflammation and atherosclerosis beyond angiogenesis. Biochem Soc Trans. 2024;52(1):137–50.

32. Tokudome T, Otani K, Mao Y, Jensen LJ, Arai Y, Miyazaki T, et al. Endothelial Natriuretic Peptide Receptor 1 Play Crucial Role for Acute and Chronic Blood Pressure Regulation by Atrial Natriuretic Peptide. Hypertension. 2022;79(7):1409–22.

33. Sogawa-Fujiwara C, Fujiwara Y, Hanagata A, Yang Q, Mihara T, Kaji N, et al. Npr2 mutant mice show vasodilation and undeveloped adipocytes in mesentery. BMC Res Notes [Internet]. 2021;14(1):1–7. Available from: 10.1186/s13104-021-05853-9

34. Špiranec K, Chen W, Werner F, Nikolaev VO, Naruke T, Werner FKA, et al. Endothelial C-type natriuretic peptide acts on pericytes to regulate microcirculatory flow and blood pressure. Circulation. 2018;138(5):494–508.

35. Acevedo LM, Barillas S, Weis SM, Göthert JR, Cheresh DA. Semaphorin 3A suppresses VEGF-mediated angiogenesis yet acts as a vascular permeability factor. Blood. 2008;111(5):2674–80.

36. Eberhard D, Balkenhol S, Köster A, Follert P, Upschulte E, Ostermann P, et al. Semaphorin-3A regulates liver sinusoidal endothelial cell porosity and promotes hepatic steatosis. Nat Cardiovasc Res. 2024;3(6):734–53.

37. Reidy KJ, Villegas G, Teichman J, Veron D, Shen W, Jimenez J, et al. Semaphorin3a regulates endothelial cell number and podocyte differentiation during glomerular development. Development. 2009;136(23):3979–89.

38. Luo X, Jian W. Different roles of endothelial cell-derived fibronectin and plasma fibronectin in endothelial dysfunction. Turkish J Med Sci. 2023;53(6):1667–77.

39. Al-Yafeai Z, Yurdagul A, Peretik JM, Alfaidi M, Murphy PA, Orr AW. Endothelial FN (Fibronectin) deposition by α5β1 integrins drives atherogenic inflammation. Arterioscler Thromb Vasc Biol. 2018;38(11):2601–14.

40. Sivakumar K, Subbiah U. Computational analysis of non-synonymous SNPs in the human LCN2 gene. Egypt J Med Hum Genet [Internet]. 2024;25(1). Available from: 10.1186/s43042-024-00565-8

41. Guardado S, Ojeda-Juárez D, Kaul M, Nordgren TM. Comprehensive review of lipocalin 2-mediated effects in lung inflammation. Am J Physiol – Lung Cell Mol Physiol. 2021;321(4):L726–33.

42. Kim SL, Shin MW, Seo SY, Kim SW. Lipocalin 2 potentially contributes to tumorigenesis from colitis via IL-6/STAT3/NF-κB signaling pathway. Biosci Rep. 2022;42(5):1–14.

43. Gáspár R, Diószegi P, Nógrádi-Halmi D, Erdélyi-Furka B, Varga Z, Kahán Z, et al. The Proteoglycans Biglycan and Decorin Protect Cardiac Cells against Irradiation-Induced Cell Death by Inhibiting Apoptosis. Cells. 2024;13(10).

44. Zeltz C, Orgel J, Gullberg D. Molecular composition and function of integrin-based collagen glues – Introducing COLINBRIs. Biochim Biophys Acta – Gen Subj [Internet]. 2014;1840(8):2533–48. Available from: 10.1016/j.bbagen.2013.12.022

45. Singh D, Rai V, K Agrawal D. Regulation of Collagen I and Collagen III in Tissue Injury and Regeneration. Cardiol Cardiovasc Med. 2023;07(01):5–16.

46. Chen M, Zhu X, Zhang L, Zhao D. COL5A2 is a prognostic-related biomarker and correlated with immune infiltrates in gastric cancer based on transcriptomics and single-cell RNA sequencing. BMC Med Genomics [Internet]. 2023;16(1):1–20. Available from: 10.1186/s12920-023-01659-9

47. Yin W, Zhu H, Tan J, Xin Z, Zhou Q, Cao Y, et al. Identification of collagen genes related to immune infiltration and epithelial-mesenchymal transition in glioma. Cancer Cell Int [Internet]. 2021;21(1):1–18. Available from: 10.1186/s12935-021-01982-0

48. Xu H, Pumiglia K, LaFlamme SE. Laminin-511 and α6 integrins regulate the expression of CXCR4 to promote endothelial morphogenesis. J Cell Sci. 2020;133(11).

49. Béguin EP, Janssen EFJ, Hoogenboezem M, Meijer AB, Hoogendijk AJ, van den Biggelaar M. Flow-induced Reorganization of Laminin-integrin Networks Within the Endothelial Basement Membrane Uncovered by Proteomics. Mol Cell Proteomics. 2020;19(7):1179–92.

50. Cai R, Wang P, Zhao X, Lu X, Deng R, Wang X, et al. LTBP1 promotes esophageal squamous cell carcinoma progression through epithelial-mesenchymal transition and cancer-associated fibroblasts transformation. J Transl Med [Internet]. 2020;18(1):1–13. Available from: 10.1186/s12967-020-02310-2

51. Klingberg F, Chau G, Walraven M, Boo S, Koehler A, Chow ML, et al. The fibronectin ED-A domain enhances recruitment of latent TGF-β-binding protein-1 to the fibroblast matrix. J Cell Sci. 2018;131(5):1–12.

52. Zhang C, Wu M, Zhang L, Shang L ru, Fang J hong. Fibrotic microenvironment promotes the metastatic seeding of tumor cells via activating the fibronectin 1/secreted phosphoprotein 1-integrin signaling. Oncotarget. 2016;7(29).

53. Xu D, Li T, Wang R, Mu R. Expression and Pathogenic Analysis of Integrin Family Genes in Systemic Sclerosis. Front Med. 2021;8(July).

54. Pan H, Lu X, Ye D, Feng Y, Wan J, Ye J. The molecular mechanism of thrombospondin family members in cardiovascular diseases. Front Cardiovasc Med. 2024;11(March):1–12.

55. Wolpe AG, Ruddiman CA, Hall PJ, Isakson BE. Polarized Proteins in Endothelium and Their Contribution to Function. J Vasc Res. 2021;58(2):65–91.

56. Lemma SA, Kuusisto M, Haapasaari KM, Sormunen R, Lehtinen T, Klaavuniemi T, et al. Integrin alpha 10, CD44, PTEN, cadherin-11 and lactoferrin expressions are potential biomarkers for selecting patients in need of central nervous system prophylaxis in diffuse large B-cell lymphoma. Carcinogenesis. 2017;38(8):812–20.

57. Agnihotri R, Crawford HC, Haro H, Matrisian LM, Havrda MC, Liaw L. Osteopontin, a Novel Substrate for Matrix Metalloproteinase-3 (Stromelysin-1) and Matrix Metalloproteinase-7 (Matrilysin). J Biol Chem [Internet]. 2001;276(30):28261–7. Available from: 10.1074/jbc.M103608200

58. Zhao Y, Huang Z, Gao L, Ma H, Chang R. Osteopontin/SPP1: a potential mediator between immune cells and vascular calcification. Front Immunol. 2024;15(June):1–11.

59. Quintero-Fabián S, Arreola R, Becerril-Villanueva E, Torres-Romero JC, Arana-Argáez V, Lara-Riegos J, et al. Role of Matrix Metalloproteinases in Angiogenesis and Cancer. Front Oncol. 2019;9(December):1–21.

60. Ito TK, Ishii G, Saito S, Yano K, Hoshino A, Suzuki T, et al. Degradation of soluble VEGF receptor-1 by MMP-7 allows VEGF access to endothelial cells. Blood. 2009;113(10):2363–9.

61. Chaqour B. Caught between a “Rho” and a hard place: are CCN1/CYR61 and CCN2/CTGF the arbiters of microvascular stiffness? J Cell Commun Signal. 2020;14(1):21–9.

62. Mo FE. Shear-Regulated Extracellular Microenvironments and Endothelial Cell Surface Integrin Receptors Intertwine in Atherosclerosis. Front Cell Dev Biol. 2021;9(April).

63. Yu Y, Gao Y, Qin J, Kuang CY, Song MB, Yu SY, et al. CCN1 promotes the differentiation of endothelial progenitor cells and reendothelialization in the early phase after vascular injury. Basic Res Cardiol. 2010;105(6):713–24.

64. Hsu PL, Chen JS, Wang CY, Wu HL, Mo FE. Shear-Induced CCN1 Promotes Atheroprone Endothelial Phenotypes and Atherosclerosis. Circulation. 2019;139(25):2877–91.

65. Surolia R, Zmijewski JW. IGFBP7: When Endothelial gCap-ing Goes Wrong in Acute Lung Injury. Am J Respir Cell Mol Biol. 2024;71(1):21–2.

66. He R, Feng B, Zhang Y, Li Y, Wang D, Yu L. IGFBP7 promotes endothelial cell repair in the recovery phase of acute lung injury. Clin Sci. 2024;138(13):797–815.

